# Consensus substrate recognition of conserved bacterial virulence peptide-bond recombinase

**DOI:** 10.64898/2026.05.12.724320

**Authors:** Kathleen Westervelt, Thomas E. Wood, Erika N. Weiskopf, Tatum D. Mortimer, Marcia B. Goldberg

## Abstract

*Shigella* OspB, a conserved type 3 effector, is a cysteine protease and peptide recombinase. Developing a protease activity-based screen, we defined and validated an OspB consensus substrate recognition motif. We found that the P1 position is aspartic acid, although cysteine is tolerated, and the P6 position an uncharged nonpolar hydrophobic residue. We demonstrate their predicted proximity to OspB active site residues within a binding groove. These findings will facilitate identification of physiological substrates of OspB and its homologs.

## Introduction

Effector proteases secreted by bacterial pathogens subvert host cell processes through precise cleavage of specific target proteins. Identification of protease substrates remains a central challenge to understanding their function in bacterial pathogenesis. Substrate specificity varies dramatically, with some proteases targeting only a single or a few substrates and others dozens (1, 2). Moreover, low-abundance cleavage events or cleavage restricted to a specific subcellular location may not reach the threshold of detection by mass spectrometry.

We previously found that OspB, a conserved effector of the *Shigella* type 3 secretion system (3, 4), is structurally predicted to align with the cysteine protease domain of bacterial multi-domain MARTX toxins and is itself a cysteine protease (5). Like MARTX toxins, OspB depends on IP_6_ for activity (5). In *Saccharomyces cerevisiae*, OspB cleaves Tco89p, an intrinsically disordered component of yeast TORC1 (5), which has no homologs in humans, the natural host for *Shigella*. To inform the identification of OspB human substrates, we defined an OspB recognition site.

## Results and Discussion

We identified the cleavage site of OspB in Tco89p by mass spectrometry of LysC and chymotryptic products of the C-terminal Tco89p OspB cleavage product. The only non-LysC, non-chymotrypsin cleavage sites were in peptides containing N-terminal Asn-303 or Ile-304 (Fig. 1*A*), with extensive peptide coverage C-terminal to Asp-302 and no peptides N-terminal to Asn-303, identifying the site of OspB cleavage in Tco89p as Asp-302. Consistent with this, substitution of Asp-302 with Ala led to loss of OspB cleavage of Tco89p (Fig. 1*B*). Single alanine substitutions in the residues flanking Tco89p Asp-302 found that cleavage by OspB requires the P6, P1, and P2’ residues Leu-297, Asp-302, and Ile-304 (Fig. 1*B*).

**Figure 1.**
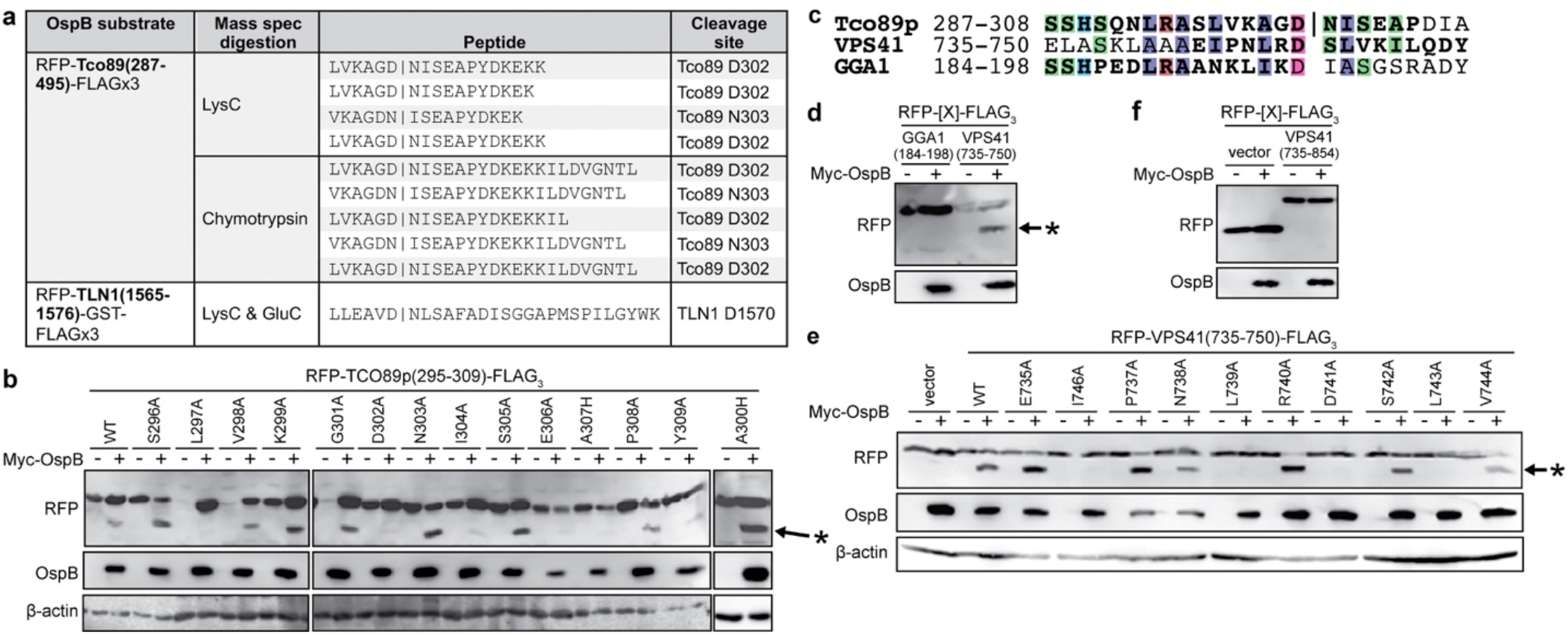
OspB substrate recognition motif identification. (*A*) OspB non-LysC and non-chymotryptic peptides of RFP-Tco89p(287-495)-FLAG_3_ and non-LysC peptides of RFP-TLN1(1565-1576)-GST-FLAG_3_ by mass spectrometry. (*B*) OspB processing of Tco89p(295-309) containing indicated alanine substitutions. *, Cleaved bands. (*C*) Alignment of BLASTP hit peptide sequences VPS41 and GGA1. Bold, peptide sequence; unbolded, vector sequence; vertical line, scissile bond. Similarity with Tco89p per Clustal2 color scheme: lavender, hydrophobic; red, positively-charged; magenta, negatively-charged; green, polar; cyan, aromatic. (*D*) OspB processing of indicated VPS41 and GGA1 peptides. (*E*) OspB processing of VPS41(735-750) containing indicated alanine substitutions. (*F*) OspB non-cleavage of 120-mer polypeptide of VPS41 (residues 735-854).

With BLASTP, we identified peptides similar to Tco89p 287-304 and tested OspB cleavage of a 16-mer peptide of VPS41 and a 15-mer peptide of GGA1, vesicular trafficking proteins. The VPS41 peptide, which contains the sequence Ile-X-X-X-X-Asp, was cleaved by OspB, but the GGA1 peptide, which contains the sequence Asn-X-X-X-X-Asp, was not (Fig. 1*C*-*D*). Alanine scanning mutagenesis of the VPS41 peptide identified several residues as required for OspB processing, including the Asp, the putative scissile bond, and the Ile of the motif (Fig. 1*E*). Yet, a 120-mer polypeptide of VPS41 that encompasses the cleavable 16-mer was not cleaved by OspB (Fig. 1*F*), indicating that OspB cleavage depends not only on a short recognition motif but on its context.

To identify an OspB consensus substrate recognition motif, using an activity-based *S. cerevisiae* assay (6), we developed an unbiased screen of OspB tolerance across the cleavage site and flanking residues within Tco89p. Oligonucleotide libraries were inserted between a Fas membrane tether and a Gal4 transcriptional activator (Fig. 2*A*-*B*). Upon introduction of the library into the reporter strain, wherein Gal4 activates transcription of *HIS3* and *URA3*, only those polypeptides that contain residues permissive to OspB cleavage should yield viable yeast on media lacking histidine and uracil. Proof-of-principle was established by good growth on selective media when the wild-type Tco89p peptide was inserted but no growth when the Asp-302-Ala substituted peptide was inserted (Fig. 2*C*).

**Figure 2.**
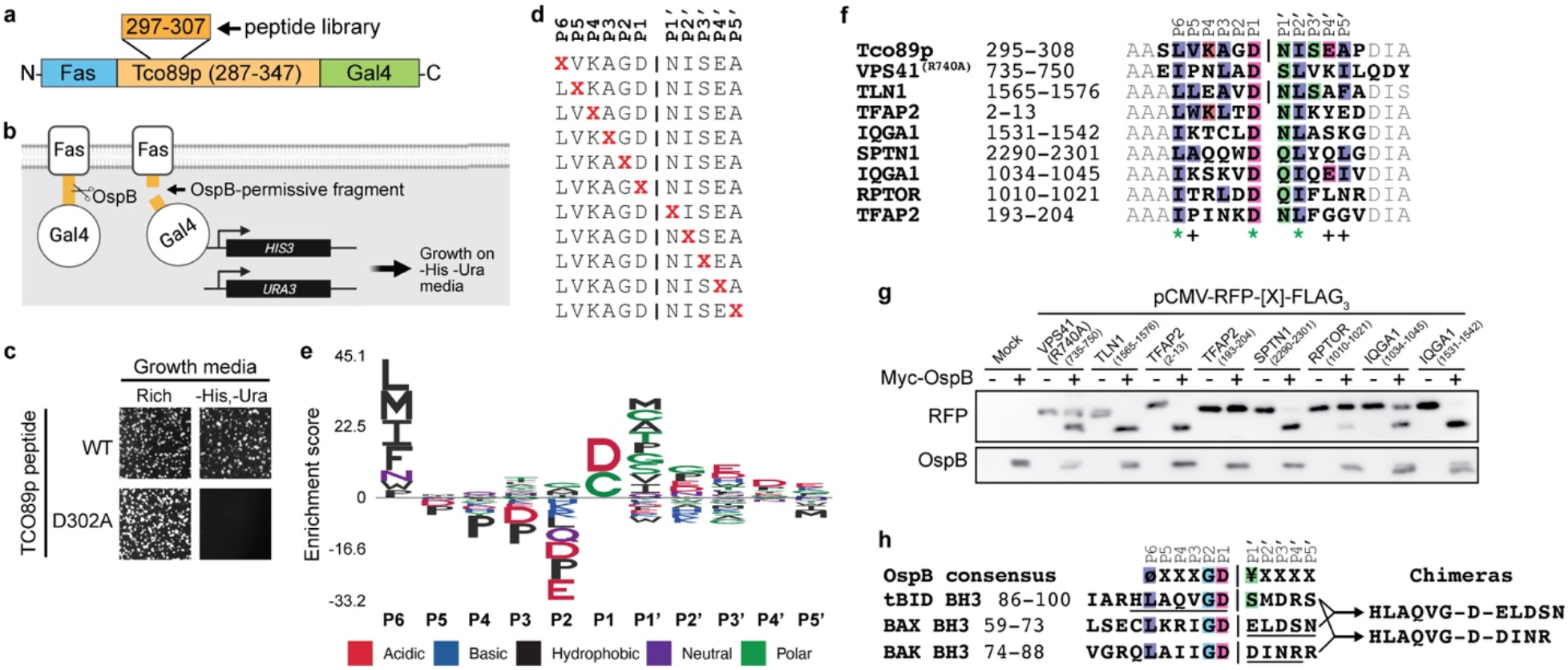
Unbiased activity-based screen identification of OspB consensus substrate recognition motif. (*A*) Peptide library screen construct. Oligonucleotide libraries of residues spanning Tco89p P6 to P5’ were inserted where indicated. (*B*) Schematic of yeast activity-based screen of OspB cleavage of peptide libraries. (*C*) Proof-of-principle of selectivity of yeast activity-based screen. Growth of yeast expressing reporter construct containing WT or D302A Tco89p(287-347) on selective media (lacking histidine and uracil, right panels) or rich media (left panels). (*D*) Design of oligonucleotide libraries surrounding the Tco89p D302-N303 scissile bond (vertical line). X, any amino acid. (*E*) Derived sequence logo for OspB substrate recognition visualized by enrichment score. (*F*) Alignment with Tco89p(296-308) of peptides of selected hits from pattern-matching algorithm. Bolded text, peptide; unbolded text, vector. (*G*) OspB cleavage of peptides in (*E*). (*H*) Alignment of OspB substrate recognition motif with OspB processing sites of tBID, BAX, BAK (4). Shown are the BH3 domains. Underlined, sequences recombined into each chimera, which is shown at right.

Each oligonucleotide library randomly encoded all 20 amino acids at a single position from P6 to P5’ surrounding the Tco89p D302-N303 scissile bond, while the remaining positions retained the wild-type Tco89p sequence (Fig. 2*D*). After transformation into the reporter strain and plating on selective and non-selective media, DNA was sequenced, with excellent library coverage. By comparing the frequencies of output variants to input variants, we generated a sequence logo for OspB substrate recognition (Fig. 2*E*).

At P1, Asp was strongly preferred (86.3%); Cys was also tolerated (13.4%). Uncharged nonpolar hydrophobic residues (Leu, Met, Ile, Phe) were strongly preferred at P6 and uncharged residues were preferred at P1’. In addition, at P2 and P1’, charged residues were deselected. At P2’, Ala was deselected, suggesting that poor cleavage with Ala in this position (Fig. 1*B*) is due to poor tolerance of Ala rather than preference for Ile.

To assess the validity of the sequence logo, we identified human proteins that contained its conserved features using a pattern-matching algorithm (7) (Supporting Dataset S1). Among those proteins identified, we tested OspB cleavage of peptides of cytoskeletal proteins TLN1 and SPTN1, the transcription factor TFAP2, the TORC1 component RPTOR, and the scaffolding protein IQGAP1; we previously found that OspB activated mTORC1 through interactions with IQGAP1 (8). All peptides except TFAP2 were cleaved by OspB, although RPTOR only weakly (Fig. 2*F*-*G*). To test whether cleavage by OspB occurred at the putative P1 Asp, we performed mass spectrometry on the C-terminal fragment of TLN1, which indicated Asp-1570 as the OspB cleavage site (Fig. 1*A*). These findings validate the strong preference for Asp at P1 and an uncharged nonpolar hydrophobic residue at P6.

Further validating this recognition site is the recent finding (4) that OspB cleaves within the BH3 domains of the apoptosis-associated BCL-2 proteins tBID, BAX, and BAK, each at an Asp. Analysis of the identified cleavage sites reveals that all three are processed at Leu-X-X-X-Gly-Asp (Fig. 2*H*). The absence of BAX and BAK in the pattern-matching search may be because acidic residues were deselected at P1’. OspB additionally mediates recombination of the N-terminal tBID cleavage product to the C-terminal BAX and BAK cleavages product, to yield chimeric fusions with the tBID Asp at the junction (4) (Fig. 2*H*). Whether OspB generates chimeras of cleavage products from substrates other than tBID, BAX, and BAK is currently unknown.

OspB homologs are present in 18 genera, several bacteriophages, and at least 41 distinct bacterial species (Supporting Dataset S2). Whereas OspB is homologous to the cysteine protease domain of MARTX toxins, which mediates auto-processing between MARTX effector domains (9-12), these auto-processing cleavage sites differ from our OspB consensus, indicating that despite similarity in the structural prediction of the OspB active site and shared dependence on IP_6_ (5, 11, 12), the OspB binding pocket differs from that of MARTX toxins. Modelling of OspB (13) showed minor conformational changes upon IP_6_ binding (Fig. 3*A*), and defines a binding groove that accommodates each substrate peptides in the same orientation and in proximity to active site residues Cys-184 and His-144 and to four additional residues required for OspB activity (Leu-59, Gly-63, Gly-246, and His-248; (4)) (Fig. 3*B*-*C*). The P1 Asp of Tco89p, tBID, and BAK approximate OspB active site residues, whereas those of TLN1 and BAX are rotated from them (Fig. 3*D*-*F*), suggesting active movement of the substrate with OspB activity.

**Figure 3.**
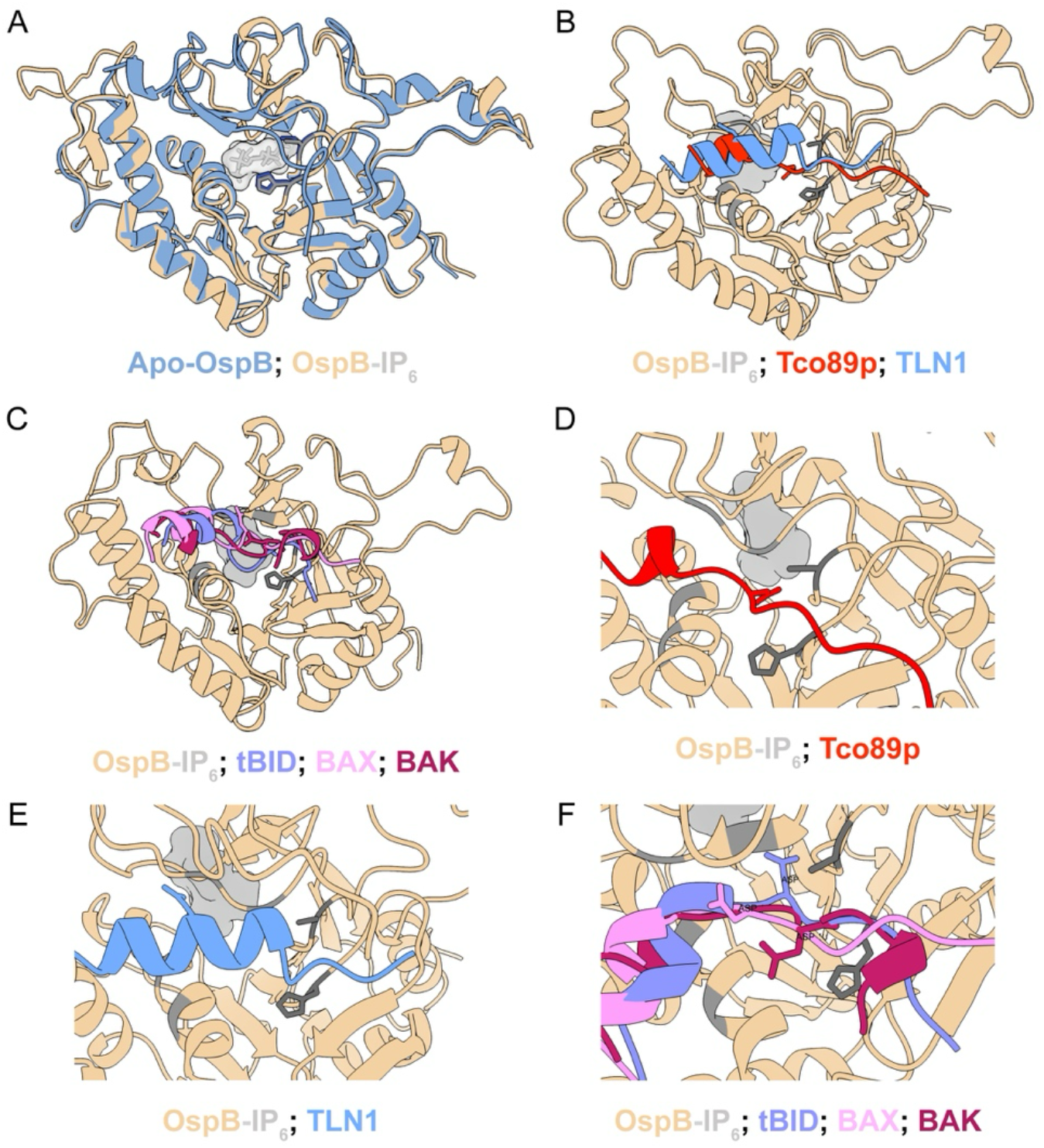
Structural modeling of OspB-substrate interface. (*A*) Overlay of predictions for apo-OspB and IP_6_-bound OspB. (*B-C*) Predicted accommodation of Tco89p and TLN1 (*B*) or tBID, BAX, and BAK (*C*) in groove of OspB. (*D-F*) Higher magnification view of substrate groove for indicated peptides. Predictions by Boltz-2 (13). Dark gray sticks, OspB active site residues Cys-184 and His-144; medium gray residues, OspB Leu-59, Gly-63, Gly-246, and His-248; light gray space-occupying shape, IP_6_; substrate sticks, P1 Asp.

These data define a consensus substrate recognition motif for OspB cysteine protease activity, define a predicted binding groove, and are consistent with OspB cleavage of multiple substrates during *Shigella* infection. Identification of protease substrates is substantially facilitated by narrowing candidates based on a substrate recognition motif. To our knowledge, this represents the first application of a yeast-based high-throughput screen for identification of a bacterial substrate recognition motif. This approach should be widely applicable to the numerous OspB homologs present in other gram-negative pathogens, as well as to other bacterial proteases that target mammalian cells.

## Acknowledgements

We thank Steven Gygi and Ross Tomaino of the Harvard Taplin Mass Spectrometry Facility for advice. This work was supported by NIH R01AI081724 (to M.B.G.), T32AI007529 (to K.W.), and T32AI1007061 (to E.N.W.).

## References

1. K. Koide, K. Ito, Y. Akiyama, Substrate recognition and binding by RseP, an Escherichia coli intramembrane protease. J Biol Chem 283, 9562–9570 (2008).

2. L. B. Evnin, J. R. Vasquez, C. S. Craik, Substrate specificity of trypsin investigated by using a genetic selection. Proc Natl Acad Sci U S A 87, 6659–6663 (1990).

3. D. V. Zurawski, K. L. Mumy, C. S. Faherty, B. A. McCormick, A. T. Maurelli, Shigella flexneri type III secretion system effectors OspB and OspF target the nucleus to downregulate the host inflammatory response via interactions with retinoblastoma protein. Mol Microbiol 71, 350–368 (2009).

4. Y. Shao et al., Bacterial effector OspB hijacks apoptosis through peptide-bond recombination of BH3 domain proteins. Cell Host Microbe 33, 1886–1900 e1889 (2025).

5. T. E. Wood et al., The Shigella Spp. Type III Effector Protein OspB Is a Cysteine Protease. mBio 10.1128/mbio.01270-22, e0127022 (2022).

6. H. Hayashi et al., Versatile assays for high throughput screening for activators or inhibitors of intracellular proteases and their cellular regulators. PLoS One 4, e7655 (2009).

7. I. Krystkowiak, J. Manguy, N. E. Davey, PSSMSearch: a server for modeling, visualization, proteome-wide discovery and annotation of protein motif specificity determinants. Nucleic Acids Res 46, W235–W241 (2018).

8. R. Lu et al., Shigella Effector OspB Activates mTORC1 in a Manner That Depends on IQGAP1 and Promotes Cell Proliferation. PLoS Pathog 11, e1005200 (2015).

9. K. L. Sheahan, C. L. Cordero, K. J. Satchell, Autoprocessing of the Vibrio cholerae RTX toxin by the cysteine protease domain. EMBO J 26, 2552–2561 (2007).

10. K. Prochazkova et al., Structural and molecular mechanism for autoprocessing of MARTX toxin of Vibrio cholerae at multiple sites. J Biol Chem 284, 26557–26568 (2009).

11. K. Prochazkova, K. J. Satchell, Structure-function analysis of inositol hexakisphosphate-induced autoprocessing of the Vibrio cholerae multifunctional autoprocessing RTX toxin. J Biol Chem 283, 23656–23664 (2008).

12. R. N. Pruitt et al., Structure-function analysis of inositol hexakisphosphate-induced autoprocessing in Clostridium difficile toxin A. J Biol Chem 284, 21934–21940 (2009).

13. S. Passaro et al., Boltz-2: Towards Accurate and Efficient Binding Affinity Prediction. bioRxiv 10.1101/2025.06.14.659707 (2025).

