## Supplemental Materials for "Consensus substrate recognition of conserved bacterial virulence peptide-bond recombinase"

**Strains and Media.** Strains used in this study are listed in Supplemental Table S3. Cloning and plasmid production were performed using *E. coli* DH10B, which was grown in Luria broth at 37°C. *Saccharomyces cerevisiae* strains were routinely cultured at 30°C in yeast extract-peptone-dextrose (YPD) broth or in synthetic selective media (MP Biomedicals) lacking leucine, histidine, and/or uracil for plasmid or library selection. Where appropriate, medium was supplemented with ampicillin (50-100 ug/ml). Yeast strains were transformed using the standard lithium acetate method. Yeast gene complementation analyses were conducted using low copy centromeric plasmids expressing the gene of interest from its native promoter unless stated otherwise. Overexpression of *TCO89* was achieved by constitutive expression of the construct from a multicopy vector. The plating efficiency of yeast strains was as described previously (1).

**Plasmid construction.** Plasmids used in this study are listed in Supplemental Table S4. All *TCO89* constructs were amplified from the *S. cerevisiae* BY4741 genome. The coding sequences for full-length Tco89p without its initiation methionine and all Tco89p fragments were amplified as BamHI-NotI fragments by PCR and cloned into pCMV-RFP-$FLAG_3$ (2).

pRS316GAL-OspB-FLAGx3-Hisx6 was generated by cloning the coding sequence for *ospB* without the initiating methionine into pRS316GAL- FLAGx3-Hisx6 (gift of C. Lesser). For complementation of the *S. cerevisiae* BY4741 *tco89*Δ strain, tagged Tco89p constructs were introduced into the single copy pRS316 plasmid under the control of the constitutive GPD promoter. First, the GAL promoter in pRS316GAL-OspB-FLAGx3-Hisx6 was replaced with the GPD promoter as a SacI-BamHI fragment from pAG413GPD-OspB (1). Then, the coding sequence for each tagged Tco89p derivative was amplified by PCR as a XbaI-ClaI fragment and cloned to replace the *ospB* gene in pRS316GPD-OspB-FLAG3-His6. The Tco89p D302A and N303S variants were generated by introducing the desired substitutions by SOEing PCR.

The correctness of all constructs was confirmed by sequencing. The sequences of primers used in this study will be made available upon request to the authors.

**Site-directed mutagenesis.** Alanine scanning site-directed mutagenesis of Tco89p and VPS41 was performed within the short Tco89p(295-309) and VPS41(735-750) regions as follows. Oligonucleotides encoding Tco89p(295-309) or VPS41(735-750) introducing an alanine-encoding codon at each triplet position therein were cloned as NotI-EcoRV fragments into NotI/EcoRV-digested pCMV-RFP-$FLAG_3$.

**Library screen.** The OspB-dependent transcriptional activator and reporter construct, which consists of the yeast transcription factor GAL4 fused at its N-terminus to the transmembrane domain of human Fas cell surface death receptor via a linker consisting of Tco89p(287-347), was based on Hayashi, et al. (3). The pRS315 vector was domesticated for compatibility with Golden Gate assembly by amplifying and introducing silent mutations via SOEing PCR into individual plasmid modules (bacterial origin, yeast centromeric sequence, ampicillin resistance cassette and leucine auxotrophic marker) to eliminate any native SapI or BsaI sites, using the approach described in Marillonnet, et al. (4). Modules were assembled into the Golden Gate-compatible pRS315GG plasmid using BsaI. A multiple cloning site (SphI-BglII-NotI-AscI-NcoI-KpnI-SacI-ApaI) was then added into the divergent SapI entry site using annealed oligonucleotides as the insert, to produce pRS315GG(MCS).

The GAL4 construct was amplified from the BY4741 genome using SOEing PCR to domesticate the construct for BsaI/SapI Golden Gate assembly, and subcloned into pCS2+MT. The Fas transmembrane anchor with a triple HA tag was codon-optimized for expression in yeast. A linker domain consisting of a ccdB cassette flanked with divergent BsaI sites was spliced between Tco89p residues 294 and 307 within Tco89p(287-347), to allow for counterselection of the parent plasmid upon insertion of variant library within the Tco89p(295-307) region. Both the Fas transmembrane and Tco89p modules were synthesized (Twist Biosciences). Transcription of the reporter plasmid was initially regulated by the GPD promoter and CYC1 terminator sequences, amplified from the plasmid pRS316GPD-ospB-FLAG3-$His_6$. The GPD promoter was PCR amplified, stitched with the Fas transmembrane anchor by SOEing PCR, and inserted into the SphI and NotI sites of pRS315GG(MCS). The Tco89p linker region was then inserted into the NotI and NcoI sites, before insertion of the domesticated GAL4 sequence, obtained from pCS2+MT-GAL4, between AscI and NcoI sites. The CYC1 terminator was PCR amplified and inserted between the NcoI and SacI sites. To reduce expression levels of the transcriptional reporter, the GPD promoter was replaced by the weak STE5 promoter. The STE5 promoter was amplified from the BY4741 genome and incorporated into pFHTFG between the SphI and BglII sites. pFHTFG was maintained in *E. coli* DB3.1, which is CcdB-resistant.

Individual site-saturated mutagenesis of Tco89p residues 297 to 307 was achieved using the NNK degenerate codon in each amino acid position, producing 11 pools of single-site variants, corresponding to the P6 to P5' positions surrounding the OspB cleavage site within Tco89p. Wild type Tco89p(295-307) and its D302A substitution were constructed as positive and negative controls, respectively. Variants of the Tco89p(287-347) region were incorporated into pFHTFG by BsaI-mediated Golden Gate assembly. This methodology eliminates the type IIS BsaI restriction sites and excises the ccdB cassette, to produce a seamless Tco89p(287-347) construct and licensing transformation of *E. coli* DH10B, which is CcdB-sensitive.

Individual single site variant libraries were recovered by pooling DH10B transformants and extraction of the plasmid libraries using a Midiprep kit (NucleoBond Xtra Midi EF, Macherey-Nagel). Libraries were transformed into the yeast reporter strain MaV103 (5), in which ATE1 and TCO89 had been deleted. ATE1, which arginylates proteins to target them to the ubiquitin pathway, was deleted to minimize degradation of the C-terminal cleavage product. For each variant library, colonies were plated on selective synthetic defined -His -Ura media and non-selective rich synthetic defined +His +Ura media. Plasmids were extracted from each plate, and amplicons were produced, barcoded, and pooled before next-generation sequencing. Sequencing resulted in sufficient sequencing depth to ensure that each variant constituted less than 1% of the total library, with an average of 1582 reads per variant. These parameters indicate excellent coverage of the library, allowing confident assessment of cleavage preferences across the mutagenized positions. Raw sequencing reads in FASTQ format were assessed for quality using FastQC version 0.11.9 (https://github.com/s-andrews/FastQC) on a per-sample basis and paired-end reads were then merged using BBMerge (6) version 38. Reads were mapped to a reference sequence of the amplicon using bwa (7) version 0.7.17. Mapping quality and coverage metrics were evaluated using bamQC from the Qualimap package (8) version 2.2.1. Individual FastQC and bamQC reports were generated for each sample, and aggregated summaries were compiled using MultiQC (9) version 1.9 for overview and comparison across all samples. As expected for targeted amplicon sequencing, FastQC flagged common warnings such as overrepresented sequences and PCR duplicates. These results reflect the non-random nature of the amplicon sequencing. Amplicon sequences from mapped reads were extracted using seqkit (10) version 2.3.1 Sequences were translated using Biopythonv 1.7.0. The frequency of output variants was compared to the input variants to build a sequence logo for the consensus sequence of the OspB cleavage site (using WebLogo18 and Logolas10). Yeast plasmid extraction, amplicon library preparation (e.g. target and index PCRs) and MiSeq Illumina sequencing were performed by Quintara Biosciences.

The resulting logo was used to identify motifs in the *H. sapiens* proteome using PSSMSearch (11) version 11. As input into the multiple sequence alignment, we used the frequencies of the amino acids at each position in the output for P6 to P5', thereby representing the experimentally observed residue preferences for this sequence.

**Mammalian cell culture and transfection.** HEK293T cells were maintained in high-glucose Dulbecco's Modified Eagle Medium (ThermoFisher 11965118) supplemented with 10% (v/v) fetal bovine serum (Sigma Aldrich F2442), at 37ºC and 5% $CO_2$. Cells were routinely tested for mycoplasma contamination using the PCR-based Mycoplasma Detection Kit (Southern Biotech 13100-01). Transient transfection was achieved using FuGENE6 transfection reagent (Promega E2692) following the manufacturer's protocol for reverse transfection. After 42 h media was replaced with DMEM containing 10 µM MG-132 (Selleck Chemicals S2619) to inhibit the proteasome and incubated for a further 6 h. Cells were washed once with cold HBSS (ThermoFisher 14025134) and lysed on ice with lysis buffer supplemented with protease and phosphatase inhibitors (10 mM HEPES pH 7.4, 150 mM NaCl, 1 mM EGTA, 100 µM MgCl2, 0.5% Triton X-100, protease inhibitor cocktail [Roche 111836170001], 1 mM PMSF, 1 mM sodium orthovanadate, 5 mM β-glycerophosphate, 1 mM sodium fluoride). Soluble lysates were separated from nuclei and cellular debris by centrifugation at 500 g for 5 min at 4ºC, and Laemmli sample buffer was added.

**SDS-PAGE analysis and western blotting.** Protein samples were analyzed by immunoblot analysis after separation by SDS-PAGE and transfer to nitrocellulose membranes using standard procedures. The extraction of protein samples from liquid yeast cultures was as described previously (1). The primary antibodies were horseradish peroxidase (HRP)-conjugated mouse anti-β-actin (Sigma, A3854; used at 1:10,000), rabbit anti-RFP (Abcam, ab62341; used at 1:1000), rabbit anti-OspB (used at 1:10,000), mouse anti-FLAG (Sigma, F3165; used at 1:5000), rabbit anti-FLAG (Sigma F7425; used at 1:1000), rat anti-α-tubulin (Santa Cruz, sc-53030; diluted to 1:200). HRP-conjugated secondary antibodies were goat anti-mouse (Jackson ImmunoResearch, 115-035-003), goat anti-rabbit (Jackson ImmunoResearch, 111-035-144), or goat anti-rat (Jackson ImmunoResearch, 112-035-175), each used at 1:5000.

**Immunoprecipitation of C-terminal cleavage products.** The C-terminal Tco89p-FLAG or TLN1-GST-FLAG cleavage product was generated in HEK293T cells by co-transfection of plasmids expressing RFP-Tco89p(287-347)-$FLAG_3$ or RFP-TLN1(1565-1576)-GST-$FLAG_3$ with Myc-OspB, as described above. Cells were collected with lysis buffer supplemented with protease and phosphatase inhibitors, and the soluble fraction was separated from the insoluble cellular debris and nuclei by pelleting at 500 g for 5 min at 4ºC. The total protein concentration of the soluble fraction was normalized to 1 mg/ml using a colorimetric γ-globulin standard (Protein Assay Kit I, Bio-Rad, 5000001). The C-terminal Tco89p or TLN1 fragment was immunoprecipitated by adding protein samples to anti-FLAG M2 magnetic beads (Millipore, M8823) equilibrated in lysis buffer and incubating with rotation at 4ºC for 2 h. Beads were washed extensively with cold immunoprecipitation buffer (10 mM HEPES pH 7.4, 150 mM NaCl, 1 mM EGTA, 100 µM MgCl2, 0.1% Triton X-100) before recovering the bound fraction by addition of 50 µl 2x concentrated Laemmli sample buffer to the beads and heating at 65ºC for 10 min.

**Mass spectrometry analysis.** Immunoprecipitated Tco89p and TLN1 samples were separated by gel electrophoresis before staining with GelCode Blue reagent (ThermoFisher 24590). The band corresponding to the OspB-mediated C-terminal Tco89p-$FLAG_3$ or TLN1-GST-$FLAG_3$ cleavage product was excised, washed in 50% (v/v) acetonitrile, and dried in a SpeedVac. Samples were reduced (20 mM TCEP, 25 mM TEAB) for 45 min at 37ºC before being cooled to room temperature and alkylated (10 mM iodoacetamide, 25 mM TEAB) in the dark for 45 min. The samples were washed in 10 mM TEAB before dehydration in acetonitrile and drying once again in a SpeedVac. The Tco89p sample was divided in two for independent digests with chymotrypsin (Promega, V1061) and LysC (VWR Pierce, PI90051); the TLN1 sample was digested with LysC and GluC, each overnight at 37ºC. Peptides were extracted using 5% formic acid in 50% acetonitrile at room temperature for 20 minutes.

LC-MS/MS analysis of the eluted peptides was performed on an Orbitrap Fusion Lumos equipped with dual pump Ultimate 3000 nanoLC (both ThermoFisher). Peptides were separated onto a microcapillary trapping column with a 100 µm inner diameter, which had been packed with approximately 5 cm of C18 Reprosil resin (5 µm, 100 Å, Dr. Maisch), followed by loading onto a 50 cm analytical µPAC column (PharmaFluidics, ESI Source Solutions). Peptide separation was achieved through applying a 5-27% acetonitrile gradient in 0.1% formic acid for 90 min at 200 nl/min. Electrospray ionization was enabled through applying a voltage of 1.8 kV using a home-made electrode junction at the end of the microcapillary column and sprayed from stainless steel emitters (ThermoFisher). The Orbitrap Fusion Lumos was operated in data-dependent mode for the mass spectrometry methods. The mass spectrometry survey scan was performed in the range of 395-1 800 m/z at a resolution of $6 \times 10^4$, followed by the selection of the thirty most intense ions (TOP30) for CID-MS2 fragmentation. Fragmentation in the ion trap used a precursor isolation width window of 2 m/z, an automatic gain control setting of 10,000, and a maximum ion accumulation of 200 ms. Singly charged ion species were not subjected to CID fragmentation. Normalized collision energy was set to 35 V and an activation time of 10 ms. Ions in a 10 ppm m/z window around ions selected for MS2 were excluded from further selection for fragmentation for 60 secs.

Raw data were submitted for analysis in Proteome Discoverer 2.4 (Thermo Scientific) software. Assignment of MS/MS spectra was performed using the Sequest HT algorithm by searching the data against a protein sequence database including all entries from the user provided database, as well as other known contaminants, such as human keratins and common lab contaminants. Sequest HT searches were performed using a 20 ppm precursor ion tolerance and requiring each peptides N-/C termini to adhere with Lys-C, chymotrypsin, and/or GluC protease specificity, as appropriate. Two missed cleavages were permitted. A MS2 spectra assignment false discovery rate (FDR) of 1% on both protein and peptide level was achieved by applying the target-decoy database search. Filtering was performed using a Percolator. Peptide N-termini and lysine residues (+229.162932 Da) were set as static modifications, whereas methionine oxidation (+15.99492 Da) was set as variable modification. Variable modifications of S/T/Y phosphorylation, acetylation, and oxidation were included in the search parameters. For quantification, a 0.02 m/z window centered on the theoretical m/z value of each the six reporter ions and the intensity of the signal closest to the theoretical m/z value was recorded. Reporter ion intensities were exported from Proteome Discoverer 2.4 search engine.

**Structural predictions.** Structural predictions were performed using the algorithm Boltz-2 (12).

### References


1. T. E. Wood *et al.*, The Shigella Spp. Type III Effector Protein OspB Is a Cysteine Protease. *mBio* 10.1128/mbio.01270-22, e0127022 (2022).
2. R. Lu *et al.*, Shigella Effector OspB Activates mTORC1 in a Manner That Depends on IQGAP1 and Promotes Cell Proliferation. *PLoS Pathog* **11**, e1005200 (2015).
3. H. Hayashi *et al.*, Versatile assays for high throughput screening for activators or inhibitors of intracellular proteases and their cellular regulators. *PLoS One* **4**, e7655 (2009).
4. S. Marillonnet, R. Grutzner, Synthetic DNA Assembly Using Golden Gate Cloning and the Hierarchical Modular Cloning Pipeline. *Curr Protoc Mol Biol* **130**, e115 (2020).
5. M. Vidal, R. K. Brachmann, A. Fattaey, E. Harlow, J. D. Boeke, Reverse two-hybrid and one-hybrid systems to detect dissociation of protein-protein and DNA-protein interactions. *Proc Natl Acad Sci U S A* **93**, 10315-10320 (1996).
6. B. Bushnell, J. Rood, E. Singer, BBMerge - Accurate paired shotgun read merging via overlap. *PLoS One* **12**, e0185056 (2017).
7. H. Li, Aligning sequence reads, clone sequences and assembly contigs with BWA-MEM. 1303.3997 http://dx.doi.org/arXiv:1303.3997.
8. K. Okonechnikov, A. Conesa, F. Garcia-Alcalde, Qualimap 2: advanced multi-sample quality control for high-throughput sequencing data. *Bioinformatics* **32**, 292-294 (2016).
9. P. Ewels, M. Magnusson, S. Lundin, M. Kaller, MultiQC: summarize analysis results for multiple tools and samples in a single report. *Bioinformatics* **32**, 3047-3048 (2016).
10. W. Shen, B. Sipos, L. Zhao, SeqKit2: A Swiss army knife for sequence and alignment processing. *Imeta* **3**, e191 (2024).
11. I. Krystkowiak, J. Manguy, N. E. Davey, PSSMSearch: a server for modeling, visualization, proteome-wide discovery and annotation of protein motif specificity determinants. *Nucleic Acids Res* **46**, W235-W241 (2018).
12. S. Passaro *et al.*, Boltz-2: Towards Accurate and Efficient Binding Affinity Prediction. *bioRxiv* 10.1101/2025.06.14.659707 (2025).

**Supporting Dataset S1. Pattern-matching algorithm search of human proteome**

| Rank | Protein accession number | GeneName | Protein name | Hit | Sequence start | Sequence stop | Position weighted matrix score | Position weighted matrix P value |
|---|---|---|---|---|---|---|---|---|
| 1 | P14923 | JUP | Junction plakoglobin | vpldpLEMHMDMDGDYpidty | 714 | 724 | 148.4 | 6.61E-07 |
| 2 | P61011 | SRP54 | Signal recognition particle 54 kDa protein | arlkkLMTIMDSMNDQeldst | 375 | 385 | 146.7 | 2.26E-06 |
| 3 | A6NFE2 | SMCO2 | Single-pass membrane and coiled-coil domain-containing protein 2 | nnkmsLQMKMDCQEQQltkkn | 14 | 24 | 146.6 | 2.36E-06 |
| 5 | Q12894 | IFRD2 | Interferon-related developmental regulator 2 | dleekLKEYVDCLTDKsaktr | 141 | 151 | 146.1 | 2.74E-06 |
| 4 | Q5VUA4 | ZNF318 | Zinc finger protein 318 | eeltaLGNLGDMPVDFcttrv | 2014 | 2024 | 146.3 | 2.74E-06 |
| 6 | Q7Z6Z7 | HUWE1 | E3 ubiquitin-protein ligase HUWE1 | vnqqqLQQLMDMGFTRehame | 1322 | 1332 | 146.1 | 2.83E-06 |
| 7 | Q8IZL8 | PELP1 | Proline-, glutamic acid- and leucine-rich protein 1 | dtaamLADFIDCPPDDekppp | 1109 | 1119 | 144.5 | 6.42E-06 |
| 8 | O75362 | ZNF217 | Zinc finger protein 217 | mptqsLLMYMDGPEVIgsslg | 14 | 24 | 144.1 | 7.64E-06 |
| 9 | Q9UBC1 | NFKBIL1 | NF-kappa-B inhibitor-like protein 1 | agrgsLWRFGDVPWPCpgggd | 294 | 304 | 144.0 | 8.21E-06 |
| 10 | Q96BJ8 | ELMO3 | Engulfment and cell motility protein 3 | spnhkLLQYGDMEEGAspptl | 572 | 582 | 143.6 | 1.04E-05 |
| 11 | P04264 | KRT1 | Keratin, type II cytoskeletal 1 | rldseLKNMQDMVEDYrnkye | 256 | 266 | 143.6 | 1.05E-05 |
| 12 | Q8TEP8 | CEP192 | Centrosomal protein of 192 kDa | pritsLCLLKDCEEIRdnren | 918 | 928 | 143.6 | 1.06E-05 |
| 13 | Q92545 | TMEM131 | Transmembrane protein 131 | sditsLIEAMDKDFDHhdspa | 1350 | 1360 | 143.4 | 1.11E-05 |
| 14 | Q0VF96 | CGNL1 | Cingulin-like protein 1 | klqreLEEQMDMNEHLqgqln | 1236 | 1246 | 142.7 | 1.44E-05 |
| 15 | P51531 | SMARCA2 | Probable global transcription activator SNF2L2 | eeefdLFMRMDMDRRRedarn | 1252 | 1262 | 142.6 | 1.45E-05 |
| 16 | Q15649 | ZNHIT3 | Zinc finger HIT domain-containing protein 3 | phlrqLMVNLDQGEDKaklmr | 113 | 123 | 142.5 | 1.51E-05 |
| 17 | Q96PX1 | RNF157 | E3 ubiquitin ligase RNF157 | rtcafLGMECDNNNDFdiasv | 611 | 621 | 142.2 | 1.71E-05 |
| 18 | Q9UID3 | VPS51 | Vacuolar protein sorting-associated protein 51 homolog | rrecpLAQLMDSETDMvrqir | 83 | 93 | 141.6 | 2.10E-05 |
| 19 | Q7Z2Y8 | GVINP1 | Interferon-induced very large GTPase 1 | treaeLRQAMDIPEEYwpspe | 130 | 140 | 141.6 | 2.11E-05 |
| 20 | Q9H6S3 | EPS8L2 | Epidermal growth factor receptor kinase substrate 8-like protein 2 | gnkdeLMQHMDEVNDElirki | 586 | 596 | 141.3 | 2.42E-05 |
| 21 | Q9UPN6 | SCAF8 | SR-related and CTD-associated factor 8 | sfnkkLMDRFDFGEDSehsee | 263 | 273 | 141.2 | 2.50E-05 |
| 22 | Q8TED9 | AFAP1L1 | Actin filament-associated protein 1-like 1 | efdcdLSDLRDMPEDDgepsk | 80 | 90 | 141.0 | 2.75E-05 |
| 23 | Q15303 | ERBB4 | Receptor tyrosine-protein kinase erbB-4 | ldeedLEDMMDAEEYLvpqaf | 1013 | 1023 | 141.0 | 2.81E-05 |
| 24 | Q96KC9 | CABS1 | Calcium-binding and spermatid-specific protein 1 | akkekLKSEDDMGTDFikstt | 75 | 85 | 141.0 | 2.81E-05 |
| 25 | P22223 | CDH3 | Cadherin-3 | wratyLIMGGDDGDHFtitth | 367 | 377 | 140.9 | 2.89E-05 |
| 26 | Q99583 | MNT | Max-binding protein MNT | elkheLSQWMDVLEIDrvlrq | 303 | 313 | 140.9 | 2.89E-05 |
| 27 | Q9NZQ9 | TMOD4 | Tropomodulin-4 | syqkeLEKYRDIDEDEilrtl | 8 | 18 | 140.9 | 2.91E-05 |
| 28 | Q8WUM9 | SLC20A1 | Sodium-dependent phosphate transporter 1 | lhklhLAKVGDCMGDSgdkpl | 397 | 407 | 140.7 | 3.10E-05 |
| 29 | Q8TDJ6 | DMXL2 | DmX-like protein 2 | dqyseLFQIQDIPTDDidlep | 1471 | 1481 | 140.7 | 3.17E-05 |
| 30 | Q6U841 | SLC4A10 | Sodium-driven chloride bicarbonate exchanger | mhantLEEIADMVLDQqvssg | 188 | 198 | 140.6 | 3.19E-05 |
| 31 | P59923 | ZNF445 | Zinc finger protein 445 | qrihsLQKQYDCHESEktpnv | 814 | 824 | 140.6 | 3.27E-05 |
| 32 | Q15910 | EZH2 | Histone-lysine N-methyltransferase EZH2 | elvnaLGQYNDDDDDDdgddp | 178 | 188 | 140.4 | 3.49E-05 |
| 33 | Q96N67 | DOCK7 | Dedicator of cytokinesis protein 7 | gsqenLRWRKDMTHWRqntek | 1442 | 1452 | 140.1 | 3.90E-05 |
| 34 | Q8IY92 | SLX4 | Structure-specific endonuclease subunit SLX4 | prsceLFSIIDVDADQepsqs | 1191 | 1201 | 140.0 | 3.91E-05 |
| 35 | P19013 | KRT4 | Keratin, type II cytoskeletal 4 | rlqseLKTMQDSVEDFktkye | 213 | 223 | 139.9 | 3.99E-05 |
| 36 | Q8NA69 | TEX45 | Testis-expressed protein 45 | qpppaLLFPMDPRWDReervs | 65 | 75 | 139.9 | 4.10E-05 |
| 37 | Q8N3X1 | FNBP4 | Formin-binding protein 4 | gglclLGAYADSDDDDndvse | 110 | 120 | 139.5 | 4.80E-05 |
| 38 | Q9HBH7 | BEX1 | Protein BEX1 | vrqpiLQYRWDMMHRLgepqa | 60 | 70 | 139.5 | 4.80E-05 |
| 39 | Q8NDV7 | TNRC6A | Trinucleotide repeat-containing gene 6A protein | glnsnLNVNMDMNSIKepqsr | 1519 | 1529 | 139.5 | 4.89E-05 |
| 40 | Q13009 | TIAM1 | T-lymphoma invasion and metastasis-inducing protein 1 | seikkLEQKIDMDEKMkkmge | 570 | 580 | 139.4 | 5.04E-05 |
| 41 | Q14692 | BMS1 | Ribosome biogenesis protein BMS1 homolog | dkkrkLKEMFDAEYDEgesty | 832 | 842 | 139.3 | 5.27E-05 |
| 42 | Q9H9L3 | ISG20L2 | Interferon-stimulated 20 kDa exonuclease-like 2 | shippLNRKADCPENAtmslk | 289 | 299 | 139.2 | 5.44E-05 |
| 43 | Q9NYZ3 | GTSE1 | G2 and S phase-expressed protein 1 | lidlpLIDFCDTPEAHvavgs | 650 | 660 | 139.2 | 5.52E-05 |
| 44 | Q9HCU4 | CELSR2 | Cadherin EGF LAG seven-pass G-type receptor 2 | geagrLEYTMDALFDSrsnqf | 216 | 226 | 139.2 | 5.64E-05 |
| 45 | O00555 | CACNA1A | Voltage-dependent P/Q-type calcium channel subunit alpha-1A | asreaLYNEMDPDERWkaayt | 793 | 803 | 139.1 | 5.78E-05 |
| 46 | Q4ZHG4 | FNDC1 | Fibronectin type III domain-containing protein 1 | stkmgLKVTWDPPKDAtsrpv | 55 | 65 | 139.0 | 5.97E-05 |
| 47 | Q16760 | DGKD | Diacylglycerol kinase delta | emdrqLRRLADTPWLCqsaep | 1083 | 1093 | 139.0 | 6.19E-05 |
| 48 | Q9NQZ2 | UTP3 | Something about silencing protein 10 | eeeevLALDMDDEDDEdggna | 83 | 93 | 138.9 | 6.37E-05 |
| 49 | Q96BU1 | S100PBP | S100P-binding protein | gisgeLCALMDQVHHMqhskw | 347 | 357 | 138.9 | 6.43E-05 |
| 50 | Q92930 | RAB8B | Ras-related protein Rab-8B | vermiLGNKCDMNDKRqvske | 119 | 129 | 138.9 | 6.47E-05 |
| 51 | Q8N7J2 | AMER2 | APC membrane recruitment protein 2 | ksfdsLTGCGDIIADQeeeag | 378 | 388 | 138.8 | 6.50E-05 |
| 52 | Q9H4T2 | ZSCAN16 | Zinc finger and SCAN domain-containing protein 16 | tkneeLFQKEDMPKDKeflge | 186 | 196 | 138.8 | 6.59E-05 |
| 53 | A6H8Y1 | BDP1 | Transcription factor TFIIIB component B" homolog | akkrsLTLRDDCQEYTtevhs | 2388 | 2398 | 138.8 | 6.60E-05 |
| 54 | P51815 | ZNF75D | Zinc finger protein 75D | vislgLKLKNDTGNDHpisvs | 276 | 286 | 138.8 | 6.63E-05 |
| 56 | Q6T4R5 | NHS | Nance-Horan syndrome protein | tagvlLSSHMDQKDDHqsssg | 617 | 627 | 138.8 | 6.65E-05 |
| 55 | Q7L0X2 | ERICH6 | Glutamate-rich protein 6 | fseeyLWKVTDIGDYDddfpd | 78 | 88 | 138.8 | 6.65E-05 |
| 57 | Q6P2P2 | PRMT9 | Protein arginine N-methyltransferase 9 | ltpekLYQTMDTHCQNemssg | 553 | 563 | 138.7 | 6.81E-05 |
| 58 | Q9ULM3 | YEATS2 | YEATS domain-containing protein 2 | pdpesLRNDGDSIEDVltqid | 1243 | 1253 | 138.7 | 6.81E-05 |
| 59 | O76039 | CDKL5 | Cyclin-dependent kinase-like 5 | rpalqLPGQMDPGWHVssvtr | 903 | 913 | 138.6 | 7.04E-05 |
| 60 | P11488 | GNAT1 | Guanine nucleotide-binding protein G(t) subunit alpha-1 | ddarkLMHMADTIEEGtmpke | 103 | 113 | 138.6 | 7.11E-05 |
| 61 | P25054 | APC | Adenomatous polyposis coli protein | nvsqsLCQEDDYEDDKptnys | 1129 | 1139 | 138.6 | 7.11E-05 |
| 62 | Q99961 | SH3GL1 | Endophilin-A2 | gdvitLTNQIDENWYEgmldg | 335 | 345 | 138.6 | 7.12E-05 |
| 63 | Q9Y3R0 | GRIP1 | Glutamate receptor-interacting protein 1 | pisshLSDLGDVEEDSspaqk | 772 | 782 | 138.3 | 7.64E-05 |
| 64 | Q96L96 | ALPK3 | Alpha-protein kinase 3 | gehglLTYICDAMELGpqral | 495 | 505 | 138.3 | 7.70E-05 |
| 65 | P15884 | TCF4 | Transcription factor 4 | hqqslLGGDMDMGNPGtlspt | 127 | 137 | 138.3 | 7.89E-05 |
| 66 | Q01851 | POU4F1 | POU domain, class 4, transcription factor 1 | vgaagLASICDSDTDPrelea | 255 | 265 | 138.2 | 7.93E-05 |
| 67 | O60244 | MED14 | Mediator of RNA polymerase II transcription subunit 14 | glktfLNMFVDSNQDArrrsv | 963 | 973 | 138.2 | 7.98E-05 |
| 70 | O14641 | DVL2 | Segment polarity protein dishevelled homolog DVL-2 | epgdmLLQVNDMNFENmsndd | 317 | 327 | 138.2 | 8.04E-05 |
| 69 | P32314 | FOXN2 | Forkhead box protein N2 | qesdsLATSIDPKEDHnysas | 292 | 302 | 138.2 | 8.04E-05 |
| 68 | Q96L73 | NSD1 | Histone-lysine N-methyltransferase, H3 lysine-36 specific | kfstlLMMLKDMHDSKtkeqr | 911 | 921 | 138.2 | 8.04E-05 |
| 71 | Q5MIZ7 | PPP4R3B | Serine/threonine-protein phosphatase 4 regulatory subunit 3B | atkgsLVGLVDYPDDEeedee | 821 | 831 | 138.1 | 8.33E-05 |
| 72 | O75386 | TULP3 | Tubby-related protein 3 | qpadnLLGDIDDLEDFvyspa | 181 | 191 | 138.1 | 8.34E-05 |
| 73 | O75309 | CDH16 | Cadherin-16 | rtyqlLVQVKDMGDQAsghqa | 210 | 220 | 138.1 | 8.35E-05 |
| 74 | Q86UD4 | ZNF329 | Zinc finger protein 329 | pclssLGDGWDCENQEghlrq | 34 | 44 | 138.1 | 8.44E-05 |
| 75 | P49257 | LMAN1 | Protein ERGIC-53 | leikqLNRQLDMILDEqrryv | 348 | 358 | 138.1 | 8.52E-05 |
| 76 | Q9UBH6 | XPR1 | Xenotropic and polytropic retrovirus receptor 1 | ddqtlLEQMMDQDDGVrnrqk | 642 | 652 | 138.0 | 8.64E-05 |
| 77 | P07949 | RET | Proto-oncogene tyrosine-protein kinase receptor Ret | wienkLYGMSDPNWPGespvp | 1061 | 1071 | 138.0 | 8.82E-05 |
| 78 | O76090 | BEST1 | Bestrophin-1 | nihttLKDHMDPYWALenrde | 567 | 577 | 138.0 | 8.87E-05 |
| 79 | Q06787 | FMR1 | Synaptic functional regulator FMR1 | ttdgsLQIRVDCNNERsvhtk | 578 | 588 | 137.9 | 8.88E-05 |
| 80 | Q2WGJ9 | FER1L6 | Fer-1-like protein 6 | meqgrLQMWVDMFPKDmpqpg | 1568 | 1578 | 137.9 | 8.88E-05 |
| 81 | Q8IY92 | SLX4 | Structure-specific endonuclease subunit SLX4 | flnsaLWDVWDGEEQRppetp | 1510 | 1520 | 137.9 | 9.06E-05 |
| 82 | Q9NP71 | MLXIPL | Carbohydrate-responsive element-binding protein | plqpsLDDFMDISDFFtnsrl | 293 | 303 | 137.8 | 9.15E-05 |
| 85 | O15370 | SOX12 | Transcription factor SOX-12 | aerlrLKHMADYPDYKyrprk | 98 | 108 | 137.8 | 9.19E-05 |

| | | | | | | | | |
|---|---|---|---|---|---|---|---|---|
| 84 | P35716 | SOX11 | Transcription factor SOX-11 | aerlrLKHMADYPDYKyrprk | 107 | 117 | 137.8 | 9.19E-05 |
| 83 | Q06945 | SOX4 | Transcription factor SOX-4 | aerlrLKHMADYPDYKyrprk | 117 | 127 | 137.8 | 9.19E-05 |
| 86 | Q5H9J7 | BEX5 | Protein BEX5 | dvpnrLVDNIDMIDGDgddme | 54 | 64 | 137.8 | 9.35E-05 |
| 87 | Q9P1Z2 | CALCOCO1 | Calcium-binding and coiled-coil domain-containing protein 1 | sdkdaLEDHMDGHFFFstqdp | 672 | 682 | 137.7 | 9.50E-05 |
| 88 | Q9C099 | LRRCC1 | Leucine-rich repeat and coiled-coil domain-containing protein 1 | kltveLMKAKDQQEDHlkhlr | 507 | 517 | 137.7 | 9.55E-05 |
| 89 | Q14257 | RCN2 | Reticulocalbin-2 | eealhLIDEMDLNGDKklsee | 271 | 281 | 137.7 | 9.57E-05 |
| 90 | A6NJ78 | METTL15 | 12S rRNA N4-methylcytidine (m4C) methyltransferase | rkdgpLDMRMDGGRYPdmpta | 192 | 202 | 137.7 | 9.60E-05 |
| 91 | O14793 | MSTN | Growth/differentiation factor 8 | pplreLIDQYDVQRDDssdgs | 90 | 100 | 137.6 | 9.85E-05 |
| 92 | Q5VV43 | KIAA0319 | Dyslexia-associated protein KIAA0319 | qrpaqLLDYGDMMLNRgspsg | 113 | 123 | 137.6 | 9.90E-05 |
| 93 | Q9HC84 | MUC5B | Mucin-5B | mpleeLGQQVDCDRMRglmca | 1393 | 1403 | 137.6 | 9.91E-05 |
| 94 | Q9BZA0 | TTTY10 | Putative transcript Y 10 protein | mkLQTLMDWEEAHeknrk | 3 | 13 | 137.5 | 9.96E-05 |
| 95 | Q96AY3 | FKBP10 | Peptidyl-prolyl cis-trans isomerase FKBP10 | dppanLFEDMDLNKDGevppe | 505 | 515 | 137.5 | 0.000100305 |
| 96 | Q9NRE2 | TSHZ2 | Teashirt homolog 2 | gsvaqLQGGNDTGTDEeletg | 42 | 52 | 137.5 | 0.0001004 |
| 97 | Q6ZN54 | DEF8 | Differentially expressed in FDCP 8 homolog | hlrlkLQELKDPNEDEpnirv | 181 | 191 | 137.5 | 0.000101438 |
| 98 | O60733 | PLA2G6 | 85/88 kDa calcium-independent phospholipase A2 | paelhLFRNYDAPETVreprf | 598 | 608 | 137.5 | 0.000102004 |
| 100 | Q96SB4 | SRPK1 | SRSF protein kinase 1 | rhkedLHNANDCDVQNlnqes | 380 | 390 | 137.5 | 0.000102664 |
| 99 | Q9UGR2 | ZC3H7B | Zinc finger CCCH domain-containing protein 7B | gtsngLGSIDDIETDCyvdpr | 200 | 210 | 137.5 | 0.000102664 |
| 101 | Q9NYT0 | PLEK2 | Pleckstrin-2 | pphisLHRIVDKMHDSntgir | 129 | 139 | 137.4 | 0.000103797 |
| 102 | Q3KR16 | PLEKHG6 | Pleckstrin homology domain-containing family G member 6 | tiiphLVVTEDTDEDAplvpd | 568 | 578 | 137.4 | 0.000103891 |
| 103 | Q68DQ2 | CRYBG3 | Very large A-kinase anchor protein | rvkthLFRSEDCNETMeienv | 1444 | 1454 | 137.4 | 0.000104457 |
| 104 | Q8N9W8 | FAM71D | Protein FAM71D | hdprdLQNMLDGGEYApfvsp | 29 | 39 | 137.4 | 0.000105212 |
| 105 | P55287 | CDH11 | Cadherin-11 | kkladLYGSKDTFDDDs | 785 | 795 | 137.3 | 0.000105967 |
| 106 | A2AJT9 | BCLAF3 | BCLAF1 and THRAP3 family member 3 | eseqtLIKIIDPNDLRhdier | 545 | 555 | 137.3 | 0.000106533 |
| 107 | Q9Y485 | DMXL1 | DmX-like protein 1 | fqddsLELKWDSDNDEenedv | 1964 | 1974 | 137.3 | 0.000108232 |
| 108 | Q6UXH8 | CCBE1 | Collagen and calcium-binding EGF domain-containing protein 1 | sfdflLLMLADIRNDItelqe | 339 | 349 | 137.3 | 0.000108515 |
| 109 | Q9UI17 | DMGDH | Dimethylglycine dehydrogenase, mitochondrial | apvtsLKARSDGTWDVetpqg | 222 | 232 | 137.2 | 0.000109175 |
| 110 | Q9H7U1 | CCSER2 | Serine-rich coiled-coil domain-containing protein 2 | ledddLMLDVDLPEDAplenv | 554 | 564 | 137.2 | 0.000109458 |
| 111 | Q6IN85 | PPP4R3A | Serine/threonine-protein phosphatase 4 regulatory subunit 3A | ttkggLVGLVDYPDDDeddde | 800 | 810 | 137.2 | 0.000110119 |
| 112 | P08865 | RPSA | 40S ribosomal protein SA | evmpdLYFYRDPEEIEkeeqa | 201 | 211 | 137.2 | 0.000113422 |
| 113 | Q8WZ42 | TTN | Titin | nrkmcLLNWSDPEDDGgseit | 16141 | 16151 | 137.1 | 0.000114365 |
| 114 | Q9H9D4 | ZNF408 | Zinc finger protein 408 | qplgpLLQDGDVDEECpaqaq | 255 | 265 | 137.1 | 0.000115026 |
| 115 | P49746 | THBS3 | Thrombospondin-3 | sdndgLGDECDGDDDNdgipd | 655 | 665 | 137.1 | 0.000116724 |
| 116 | P54826 | GAS1 | Growth arrest-specific protein 1 | gsdggLDDYYDEDYDDeqrtg | 261 | 271 | 137.1 | 0.000117668 |
| 117 | Q9HCK1 | ZDBF2 | DBF4-type zinc finger-containing protein 2 | dsvrnLKKAKDVIEDNpdepv | 1801 | 1811 | 137.0 | 0.000117857 |
| 118 | Q9BPZ3 | PAIP2 | Polyadenylate-binding protein-interacting protein 2 | ipardLPQTMDQIQDQfndlv | 80 | 90 | 137.0 | 0.000120027 |
| 119 | Q12879 | GRIN2A | Glutamate receptor ionotropic, NMDA 2A | daclrMGNLYDIDEDQmlqet | 1242 | 1252 | 137.0 | 0.000120782 |
| 120 | Q9HAS0 | C17orf75 | Protein Njmu-R1 | mlpsLQESMDGDEKElesse | 5 | 15 | 137.0 | 0.000121537 |
| 121 | Q15027 | ACAP1 | Arf-GAP with coiled-coil, ANK repeat and PH domain-containing protein 1 | tmadaLAHGADVNWVNggqdn | 592 | 602 | 136.9 | 0.000122858 |
| 122 | Q8WW62 | TMED6 | Transmembrane emp24 domain-containing protein 6 | kerkqLNDTLDAIEDGtqkvq | 155 | 165 | 136.9 | 0.000123329 |
| 123 | Q96BJ8 | ELMO3 | Engulfment and cell motility protein 3 | admraLLTGKDCPHVRekgsg | 602 | 612 | 136.9 | 0.000123896 |
| 124 | Q9NYL9 | TMOD3 | Tropomodulin-3 | pfrkdLEKYKDLDEDEllgnl | 9 | 19 | 136.9 | 0.000124084 |
| 125 | Q8IZQ1 | WDFY3 | WD repeat and FYVE domain-containing protein 3 | epaevLEMQEDCPEAQigqea | 3263 | 3273 | 136.9 | 0.000124273 |
| 126 | Q9ULH0 | KIDINS220 | Kinase D-interacting substrate of 220 kDa | grksfLMKRGDVIDYSssgvs | 1451 | 1461 | 136.9 | 0.000126066 |
| 127 | Q3YEC7 | RABL6 | Rab-like protein 6 | iaaqmLSFVMDDPDFEsegsd | 566 | 576 | 136.8 | 0.000127576 |
| 128 | Q6UB99 | ANKRD11 | Ankyrin repeat domain-containing protein 11 | kdeksLKRIKDTNKDIsrsfr | 718 | 728 | 136.8 | 0.000128142 |
| 129 | Q5QJE6 | DNTTIP2 | Deoxynucleotidyltransferase terminal-interacting protein 2 | ikasdLTKFGDCGGSDdeees | 388 | 398 | 136.8 | 0.000128991 |
| 131 | P02585 | TNNC2 | Troponin C, skeletal muscle | eeiesLMKDGDKNNDGridfd | 135 | 145 | 136.7 | 0.000129935 |
| 130 | P63316 | TNNC1 | Troponin C, slow skeletal and cardiac muscles | ddieeLMKDGDKNNDGridyd | 136 | 146 | 136.7 | 0.000129935 |
| 133 | P24844 | MYL9 | Myosin regulatory light polypeptide 9 | hlrelLTTMGDRFTDEevdem | 127 | 137 | 136.7 | 0.000130973 |
| 132 | P28289 | TMOD1 | Tropomodulin-1 | syrreLEKYRDLDEDEilgal | 7 | 17 | 136.7 | 0.000130973 |
| 134 | Q3T8J9 | GON4L | GON-4-like protein | seseaLMLVWDASETEklpgt | 2029 | 2039 | 136.7 | 0.000131728 |
| 135 | Q2TAC6 | KIF19 | Kinesin-like protein KIF19 | kvdeqMVVLMDPMEDPddilr | 41 | 51 | 136.6 | 0.000135313 |
| 136 | Q4VC44 | FLYWCH1 | FLYWCH-type zinc finger-containing protein 1 | fsklvLLTASDQDEDGvgskp | 44 | 54 | 136.6 | 0.000135691 |
| 137 | Q96C92 | ENTR1 | Endosome-associated-trafficking regulator 1 | rhlrtLQISYDALKDEnsklr | 267 | 277 | 136.6 | 0.000138238 |
| 138 | O14646 | CHD1 | Chromodomain-helicase-DNA-binding protein 1 | ssdrhLTQYHDHHKDRhqgds | 1550 | 1560 | 136.5 | 0.000138616 |
| 140 | Q14563 | SEMA3A | Semaphorin-3A | rngdpLTHCSDLHHDNhhghs | 564 | 574 | 136.5 | 0.000138805 |
| 139 | Q8TE12 | LMX1A | LIM homeobox transcription factor 1-alpha | sddtsLSNLGDCFLATseagp | 345 | 355 | 136.5 | 0.000138805 |
| 141 | O60861 | GAS7 | Growth arrest-specific protein 7 | gelitLLQVPDGGWWEgeked | 31 | 41 | 136.5 | 0.000139465 |
| 142 | Q9Y5S2 | CDC42BPB | Serine/threonine-protein kinase MRCK beta | svtvpLRSMSDPDQDFdkepd | 1655 | 1665 | 136.5 | 0.000140597 |
| 144 | Q6P995 | FAM171B | Protein FAM171B | sndtsLDSGVDMNELHssrkl | 705 | 715 | 136.5 | 0.000140692 |
| 143 | Q9NY74 | ETAA1 | Ewing's tumor-associated antigen 1 | vdddlLYQACDDIERLtqqqd | 610 | 620 | 136.5 | 0.000140692 |
| 145 | Q8TEW0 | PARD3 | Partitioning defective 3 homolog | akkgmLKGLGDMFRFGkhrkd | 1017 | 1027 | 136.5 | 0.000142768 |
| 147 | Q5T7V8 | GORAB | RAB6-interacting golgin | kkleeLMQQLDVEADEetlel | 295 | 305 | 136.4 | 0.000143428 |
| 146 | Q8IZY2 | ABCA7 | Phospholipid-transporting ATPase ABCA7 | leevfLYFSKDQGKDEdteeq | 2098 | 2108 | 136.4 | 0.000143428 |
| 148 | Q9UFE4 | CCDC39 | Coiled-coil domain-containing protein 39 | qekeeLQREGDCLDAKinkae | 674 | 684 | 136.4 | 0.000144183 |
| 149 | Q8NDB2 | BANK1 | B-cell scaffold protein with ankyrin repeats | iqqekLRQLRDCIIGKrpeee | 719 | 729 | 136.4 | 0.000144655 |
| 150 | Q9NQ31 | AKIP1 | A-kinase-interacting protein 1 | rgeskLHMCLDIGNGQrkdrk | 126 | 136 | 136.4 | 0.000144844 |
| 151 | Q96GX5 | MASTL | Serine/threonine-protein kinase greatwall | niedpLIVTPDCQEKTspkgv | 608 | 618 | 136.4 | 0.000145315 |
| 152 | Q8N7J2 | AMER2 | APC membrane recruitment protein 2 | agtgtLAADMDLHCDCaaetp | 39 | 49 | 136.3 | 0.000146165 |
| 153 | Q9Y608 | LRRFIP2 | Leucine-rich repeat flightless-interacting protein 2 | neknnLIYQVDTLKDVieeqe | 396 | 406 | 136.3 | 0.000146731 |
| 154 | Q9UQN3 | CHMP2B | Charged multivesicular body protein 2b | mindtLDDIFDGSDDEeesqd | 146 | 156 | 136.3 | 0.000148429 |
| 155 | Q155Q3 | DIXDC1 | Dixin | lkqelLRANMDKDELHnqnvd | 393 | 403 | 136.3 | 0.000148524 |
| 156 | O75976 | CPD | Carboxypeptidase D | dgfhrLRQHHDEYEDEirmms | 1337 | 1347 | 136.3 | 0.000149279 |
| 157 | O95391 | SLU7 | Pre-mRNA-splicing factor SLU7 | hvqpqLMFDYDGKRDRwngyn | 159 | 169 | 136.3 | 0.0001506 |
| 161 | O43812 | DUX1 | Double homeobox protein 1 | atkeeLAQGIDIPEPRvqiwf | 52 | 62 | 136.2 | 0.000151449 |
| 158 | O94830 | DDHD2 | Phospholipase DDHD2 | sekdsLNIVMDQGDTPtleed | 378 | 388 | 136.2 | 0.000151449 |
| 160 | Q96PT3 | DUX5 | Double homeobox protein 5 | atkeeLAQGIDIPEPRvqiwf | 79 | 89 | 136.2 | 0.000151449 |
| 159 | Q96PT4 | DUX3 | Putative double homeobox protein 3 | atkeqLAQGIDIPEPRvqiwf | 79 | 89 | 136.2 | 0.000151449 |
| 162 | Q9Y597 | KCTD3 | BTB/POZ domain-containing protein KCTD3 | iqmwdLTTAMDMVNKSedkdv | 569 | 579 | 136.2 | 0.000152204 |
| 163 | Q9H799 | CPLANE1 | Ciliogenesis and planar polarity effector 1 | klthnLFEQGDAGHLQllkvk | 2437 | 2447 | 136.2 | 0.000153053 |
| 164 | Q9H9C1 | VIPAS39 | Spermatogenesis-defective protein 39 homolog | ravnsLRDFVDDDDDDdlerv | 42 | 52 | 136.2 | 0.000153902 |
| 165 | Q96K80 | ZC3H10 | Zinc finger CCCH domain-containing protein 10 | grrhdLYDIYDLPDRGfedhe | 189 | 199 | 136.2 | 0.000154185 |
| 166 | P17023 | ZNF19 | Zinc finger protein 19 | palisLLERGDMAWGLeaqdd | 65 | 75 | 136.2 | 0.000154563 |
| 167 | Q8IYT3 | CCDC170 | Coiled-coil domain-containing protein 170 | kheefLTQLRDCLDPDerndk | 172 | 182 | 136.1 | 0.000155412 |
| 168 | Q9H6T3 | RPAP3 | RNA polymerase II-associated protein 3 | eetgnLIQTIDVPDSTtaaap | 443 | 453 | 136.1 | 0.000155506 |
| 169 | Q7Z333 | SETX | Probable helicase senataxin | dkknpLGNCGDINLVRlgpek | 2035 | 2045 | 136.1 | 0.000156073 |
| 170 | Q2VIQ3 | KIF4B | Chromosome-associated kinesin KIF4B | pnskiLKEMCDMEQVLskkta | 1169 | 1179 | 136.1 | 0.000156922 |
| 171 | O14730 | RIOK3 | Serine/threonine-protein kinase RIO3 | mlaqmLQMEYDREYDAqlrre | 84 | 94 | 136.1 | 0.000157205 |

| | | | | | | | | |
|---|---|---|---|---|---|---|---|---|
| 172 | Q13033 | STRN3 | Striatin-3 | etfnfLENADDSDEDEendmi | 251 | 261 | 136.1 | 0.000157488 |
| 173 | Q6N021 | TET2 | Methylcytosine dioxygenase TET2 | lhvlpLYKVSDVDEFGsveaq | 1420 | 1430 | 136.1 | 0.000158243 |
| 174 | Q8IYK8 | REM2 | GTP-binding protein REM 2 | mhtdLDTDMDMDTETtalcp | 5 | 15 | 136.1 | 0.00015947 |
| 175 | Q9HCE6 | ARHGEF10L | Rho guanine nucleotide exchange factor 10-like protein | rdmqeLKHKYDCKMTQlmkaa | 235 | 245 | 136.1 | 0.000160224 |
| 176 | Q2UY09 | COL28A1 | Collagen alpha-1(XXVIII) chain | eisesLSVTRDQDEDDkapep | 1023 | 1033 | 136.0 | 0.000161829 |
| 177 | Q13164 | MAPK7 | Mitogen-activated protein kinase 7 | dleefLNQSFDMGVADgpqdg | 751 | 761 | 136.0 | 0.000162112 |
| 178 | Q09666 | AHNAK | Neuroblast differentiation-associated protein AHNAK | veapsLDVHMDSPDINiegpd | 4954 | 4964 | 136.0 | 0.000162583 |
| 179 | Q5VUB5 | FAM171A1 | Protein FAM171A1 | sndasLDSGVDMNEPKsarkg | 740 | 750 | 136.0 | 0.000164188 |
| 180 | A2RUR9 | CCDC144A | Coiled-coil domain-containing protein 144A | deynhLKERMDQCEKEkagrk | 1109 | 1119 | 136.0 | 0.000164754 |
| 181 | Q8IYA2 | CCDC144CP | Putative coiled-coil domain-containing protein 144C | deynhLKERMDQCEKEkagrk | 919 | 929 | 136.0 | 0.000164754 |
| 182 | Q8N108 | MIER1 | Mesoderm induction early response protein 1 | gnkkpLHADMDTNGYEtdnlt | 437 | 447 | 136.0 | 0.000165603 |
| 184 | Q7RTP6 | MICAL3 | [F-actin]-monooxygenase MICAL3 | haadpLEIQADVHWTHirere | 1034 | 1044 | 135.9 | 0.000167868 |
| 183 | Q96JM7 | L3MBTL3 | Lethal(3)malignant brain tumor-like protein 3 | kkkpkLSLKADTKEDGeerdd | 177 | 187 | 135.9 | 0.000167868 |
| 185 | Q6ZVL6 | KIAA1549L | UPF0606 protein KIAA1549L | evnkaLKQKSDIEHYRnklrl | 1395 | 1405 | 135.9 | 0.000169849 |
| 186 | Q13905 | RAPGEF1 | Rap guanine nucleotide exchange factor 1 | msrltLKQEGDDGPDVrggsg | 682 | 692 | 135.9 | 0.00017051 |
| 187 | Q9H4R4 | NCOR1P1 | Putative nuclear receptor corepressor 1-like protein NCOR1P1 | lskeeLIECMDRVDREiakve | 72 | 82 | 135.9 | 0.000171076 |
| 188 | Q9P2B7 | CFAP97 | Cilia- and flagella-associated protein 97 | hkqkvLHDTMDLNHLLkaflq | 340 | 350 | 135.8 | 0.000171453 |
| 189 | Q6UXB8 | PI16 | Peptidase inhibitor 16 | rrgenLFAITDEGMDVplame | 89 | 99 | 135.8 | 0.00017202 |
| 190 | P0CE71 | OCM2 | Putative oncomodulin-2 | setksLMAAADNDGDGkigae | 86 | 96 | 135.8 | 0.000172397 |
| 191 | P0CE72 | OCM | Oncomodulin-1 | setksLMAAADNDGDGkigae | 86 | 96 | 135.8 | 0.000172397 |
| 192 | O96001 | PPP1R17 | Protein phosphatase 1 regulatory subunit 17 | vfsehLIKRYDVQERHpkgkm | 85 | 95 | 135.8 | 0.000172491 |
| 194 | Q5T7N2 | L1TD1 | LINE-1 type transposase domain-containing protein 1 | srmdiLEERIDSLEDQieefs | 659 | 669 | 135.8 | 0.000174284 |
| 193 | Q8TEP8 | CEP192 | Centrosomal protein of 192 kDa | nraspLEQAQDSPIDFhlqsw | 151 | 161 | 135.8 | 0.000174284 |
| 195 | Q96MW1 | CCDC43 | Coiled-coil domain-containing protein 43 | rkaalLAQYADVTDEEdeade | 132 | 142 | 135.8 | 0.000174945 |
| 196 | O15061 | SYNM | Synemin | sdkveLGVIGDSVHMEglpgs | 1289 | 1299 | 135.8 | 0.000175888 |
| 197 | Q9H446 | RWDD1 | RWD domain-containing protein 1 | evdesLFQEMDDLELEddedd | 214 | 224 | 135.8 | 0.000175888 |
| 198 | Q02818 | NUCB1 | Nucleobindin-1 | rlrmlLKAKMDAEQDPnvqvd | 115 | 125 | 135.8 | 0.000176077 |
| 199 | Q96RU2 | USP28 | Ubiquitin carboxyl-terminal hydrolase 28 | ankevLAKVIDLTHDNkddlq | 88 | 98 | 135.8 | 0.000176926 |
| 201 | Q5SW79 | CEP170 | Centrosomal protein of 170 kDa | gkiriLFKDKDRNWDDieskl | 1442 | 1452 | 135.7 | 0.000177021 |
| 200 | Q96L14 | CEP170P1 | Cep170-like protein | gkiriLFKDKDRNWDDieskl | 151 | 161 | 135.7 | 0.000177021 |
| 202 | Q8N5C8 | TAB3 | TGF-beta-activated kinase 1 and MAP3K7-binding protein 3 | smnrqLQINVDCTLKEvdllq | 584 | 594 | 135.7 | 0.000177115 |
| 203 | Q9BTW9 | TBCD | Tubulin-specific chaperone D | seqkpLILTEDDDEDDdvpeg | 342 | 352 | 135.7 | 0.000177398 |
| 204 | Q92665 | MRPS31 | 28S ribosomal protein S31, mitochondrial | rsrpeLRIQFDEGYDNypgqe | 224 | 234 | 135.7 | 0.000177587 |
| 205 | Q6IN85 | PPP4R3A | Serine/threonine-protein phosphatase 4 regulatory subunit 3A | gglvgLVDYPDDDEDDdeded | 803 | 813 | 135.7 | 0.00018004 |
| 206 | Q8NFU7 | TET1 | Methylcytosine dioxygenase TET1 | lhvlpLYKLSDTDEFGskegm | 1710 | 1720 | 135.7 | 0.000181078 |
| 207 | Q92597 | NDRG1 | Protein NDRG1 | ptkttLLKMADCGGLPqisqp | 283 | 293 | 135.7 | 0.000181172 |
| 208 | Q13474 | DRP2 | Dystrophin-related protein 2 | svserLKQLQDAHRDFgpgsq | 332 | 342 | 135.7 | 0.000181267 |
| 209 | Q8TC99 | FNDC8 | Fibronectin type III domain-containing protein 8 | maseaLHQVGDGEEAVlkken | 6 | 16 | 135.7 | 0.00018155 |
| 210 | O94964 | SOGA1 | Protein SOGA1 | venrgLRAEMDDMKDHgggcg | 236 | 246 | 135.7 | 0.000181739 |
| 211 | Q15057 | ACAP2 | Arf-GAP with coiled-coil, ANK repeat and PH domain-containing protein 2 | kmaeaLAHGADVNWANseenk | 626 | 636 | 135.6 | 0.000183248 |
| 212 | Q92540 | SMG7 | Protein SMG7 | stsrnLSNNCDTGEKPvvtfk | 533 | 543 | 135.6 | 0.000183248 |
| 214 | O15079 | SNPH | Syntaphilin | eassILSSGVDCGTEEtslhs | 264 | 274 | 135.6 | 0.000185702 |
| 213 | Q9BQI3 | EIF2AK1 | Eukaryotic translation initiation factor 2-alpha kinase 1 | ielpsLEVLSDQEEDReqcgv | 254 | 264 | 135.6 | 0.000185702 |
| 215 | Q9BQ70 | TCF25 | Transcription factor 25 | nnrfeLINIDDLEDDPvvnge | 64 | 74 | 135.6 | 0.000185985 |
| 216 | Q03518 | TAP1 | Antigen peptide transporter 1 | sfisgLPQGYDTEVDEagsql | 688 | 698 | 135.6 | 0.000186079 |
| 217 | Q96B67 | ARRDC3 | Arrestin domain-containing protein 3 | flpppLYSEIDPNPDQsaddr | 393 | 403 | 135.6 | 0.000186362 |
| 218 | Q9Y6I9 | TEX264 | Testis-expressed protein 264 | wkwrgLVEAIDTQVDGtgadt | 205 | 215 | 135.6 | 0.000186457 |
| 219 | Q9H254 | SPTBN4 | Spectrin beta chain, non-erythrocytic 4 | geqelLMMSEDKGKDEqstlq | 1643 | 1653 | 135.5 | 0.00019108 |
| 220 | Q9BXN1 | ASPN | Asporin | lknmmLKDMEDTDDDDdddd | 31 | 41 | 135.5 | 0.000191269 |
| 221 | Q15858 | SCN9A | Sodium channel protein type 9 subunit alpha | drdddLLNKKDMAFDNvnens | 1926 | 1936 | 135.5 | 0.000191835 |
| 222 | Q9BZE4 | GTPBP4 | Nucleolar GTP-binding protein 1 | kemrsLGVDMDDKDDAhyavq | 527 | 537 | 135.5 | 0.000192496 |
| 223 | Q6ZU65 | UBN2 | Ubinuclein-2 | hrkdrLQDLIDIGFGYdetdp | 199 | 209 | 135.5 | 0.000192873 |
| 224 | Q8NER1 | TRPV1 | Transient receptor potential cation channel subfamily V member 1 | ksrtrLFGKGDSEEAFpvdcp | 49 | 59 | 135.5 | 0.000193345 |
| 225 | Q9NYV4 | CDK12 | Cyclin-dependent kinase 12 | mappdLPHWQDCHELWskkrr | 1033 | 1043 | 135.5 | 0.000193628 |
| 226 | Q9HA90 | EFCC1 | EF-hand and coiled-coil domain-containing protein 1 | daaaeLATDGDSDTDEearla | 117 | 127 | 135.5 | 0.000193911 |
| 227 | P0DN79 | CBSL | Cystathionine beta-synthase-like protein | kdrisLRMIEDAERDGtlkpg | 124 | 134 | 135.4 | 0.000195798 |
| 228 | P35520 | CBS | Cystathionine beta-synthase | kdrisLRMIEDAERDGtlkpg | 124 | 134 | 135.4 | 0.000195798 |
| 229 | Q92681 | RSC1A1 | Regulatory solute carrier protein family 1 member 1 | hdqeyLCNIGDLELPEerqqn | 190 | 200 | 135.4 | 0.000196931 |
| 230 | Q06730 | ZNF33A | Zinc finger protein 33A | higenLMNEMDIRNFQpqvsl | 775 | 785 | 135.4 | 0.00019778 |
| 231 | Q14004 | CDK13 | Cyclin-dependent kinase 13 | mpppdLPLWQDCHELWskkrr | 1011 | 1021 | 135.4 | 0.00019778 |
| 232 | P22223 | CDH3 | Cadherin-3 | gtgtlLLTLIDVNDHGpvpep | 535 | 545 | 135.4 | 0.000199195 |
| 233 | Q05193 | DNM1 | Dynamin-1 | alkeaLSIIGDINTTTvstpm | 739 | 749 | 135.4 | 0.000199573 |
| 234 | O95153 | TSPOAP1 | Peripheral-type benzodiazepine receptor-associated protein 1 | regqiLKVFGDKDADGfyqge | 1660 | 1670 | 135.3 | 0.00019995 |
| 235 | Q8NDI1 | EHBP1 | EH domain-binding protein 1 | lvekrLRYLMDTGRNTeeeea | 1099 | 1109 | 135.3 | 0.000200422 |
| 236 | P51572 | BCAP31 | B-cell receptor-associated protein 31 | eehakLQAAVDGPMDKkee | 233 | 243 | 135.3 | 0.000201743 |
| 237 | Q8IUW3 | SPATA2L | Spermatogenesis-associated protein 2-like protein | ayrapLDLYRDLQEDEgseda | 225 | 235 | 135.3 | 0.000202687 |
| 238 | Q9BX66 | SORBS1 | Sorbin and SH3 domain-containing protein 1 | tsqasLHMNGDGGVHTpssgi | 1072 | 1082 | 135.3 | 0.000203347 |
| 239 | Q9Y4F4 | TOGARAM1 | TOG array regulator of axonemal microtubules protein 1 | keekeLFHNKDCEKKEknswe | 1191 | 1201 | 135.3 | 0.000203347 |
| 240 | Q9H8W4 | PLEKHF2 | Pleckstrin homology domain-containing family F member 2 | slkspLNDMSDDDDDDdssd | 235 | 245 | 135.3 | 0.000203442 |
| 242 | Q03164 | KMT2A | Histone-lysine N-methyltransferase 2A | qnpsrLAVISDSGEKRvtite | 2949 | 2959 | 135.3 | 0.000203725 |
| 241 | Q9Y618 | NCOR2 | Nuclear receptor corepressor 2 | lskeeLIQNMDRVDREitmve | 172 | 182 | 135.3 | 0.000203725 |
| 243 | Q71F56 | MED13L | Mediator of RNA polymerase II transcription subunit 13-like | pslhdLDNIFDNSDDDelgav | 810 | 820 | 135.3 | 0.000204102 |
| 244 | Q8TF72 | SHROOM3 | Protein Shroom3 | dpdsrLKTTMDLMEGLfprdv | 1692 | 1702 | 135.2 | 0.000207782 |
| 245 | Q96S82 | UBL7 | Ubiquitin-like protein 7 | qwqpqLQQLRDMGIQDdelsl | 342 | 352 | 135.2 | 0.000207877 |
| 246 | P21709 | EPHA1 | Ephrin type-A receptor 1 | kaqgeLGWLLDPPKDGwseqq | 40 | 50 | 135.2 | 0.000208254 |
| 247 | P46940 | IQGAP1 | Ras GTPase-activating-like protein IQGAP1 | evsltLTNKFDVPGDEnaemd | 1365 | 1375 | 135.2 | 0.000208537 |
| 248 | Q9P2D1 | CHD7 | Chromodomain-helicase-DNA-binding protein 7 | rgtdmLADGGDGGEFDreded | 1839 | 1849 | 135.2 | 0.000208537 |
| 249 | P40763 | STAT3 | Signal transducer and activator of transcription 3 | fnyktLKSQGDMQDLNgnnqs | 179 | 189 | 135.2 | 0.000208631 |
| 250 | Q6UWT4 | C5orf46 | Uncharacterized protein C5orf46 | llvlfLTCYADDKPDKpddkp | 19 | 29 | 135.2 | 0.000208726 |
| 251 | O43525 | KCNQ3 | Potassium voltage-gated channel subfamily KQT member 3 | srikyLQTRIDMIFTPgppst | 564 | 574 | 135.2 | 0.000209009 |
| 252 | Q8N806 | UBR7 | Putative E3 ubiquitin-protein ligase UBR7 | teddgLVRNIDGIGDQevikp | 211 | 221 | 135.2 | 0.000210236 |
| 253 | P12107 | COL11A1 | Collagen alpha-1(XI) chain | grdsdLLVDGDLGEYDfyeyk | 372 | 382 | 135.2 | 0.000211368 |
| 254 | Q9H3T2 | SEMA6C | Semaphorin-6C | lpvkhLRAAGDPWEWNqnrnn | 712 | 722 | 135.2 | 0.000212783 |
| 255 | O60284 | ST18 | Suppression of tumorigenicity 18 protein | sliqeLSVAYDCSMAKkrtae | 28 | 38 | 135.2 | 0.000213633 |
| 256 | P29375 | KDM5A | Lysine-specific demethylase 5A | aqleeLMMVGDLLEVSldetq | 1448 | 1458 | 135.1 | 0.000213916 |
| 257 | P29558 | RBMS1 | RNA-binding motif, single-stranded-interacting protein 1 | pteplLCKFADGGQKKrqnpn | 220 | 230 | 135.1 | 0.000214765 |
| 258 | Q6XE24 | RBMS3 | RNA-binding motif, single-stranded-interacting protein 3 | psepILCKFADGGQKKrqnqs | 219 | 229 | 135.1 | 0.000214765 |
| 259 | Q96I76 | GPATCH3 | G patch domain-containing protein 3 | kggsgLVFYTDAQFWQeeegd | 336 | 346 | 135.1 | 0.000214765 |

| | | | | | | | | |
|---|---|---|---|---|---|---|---|---|
| 260 | Q86VP3 | PACS2 | Phosphofurin acidic cluster sorting protein 2 | edldlLYDTLDMEHPSdsgpd | 289 | 299 | 135.1 | 0.000215992 |
| 261 | O00534 | VWA5A | von Willebrand factor A domain-containing protein 5A | laaksLLQTKDMGLREtpasd | 553 | 563 | 135.1 | 0.000216275 |
| 262 | P49454 | CENPF | Centromere protein F | lenseLKKSLDCMHKDqveke | 2674 | 2684 | 135.1 | 0.000216463 |
| 263 | Q8NFU7 | TET1 | Methylcytosine dioxygenase TET1 | plskgLEKQHDCDYKIlpalg | 134 | 144 | 135.1 | 0.000217313 |
| 264 | Q13206 | DDX10 | Probable ATP-dependent RNA helicase DDX10 | eeeafLDWSDDDDDDdgfdp | 777 | 787 | 135.1 | 0.000217784 |
| 265 | Q14137 | BOP1 | Ribosome biogenesis protein BOP1 | vgnvpLEWYDDFPHVGydldg | 141 | 151 | 135.1 | 0.000218256 |
| 266 | Q86UP3 | ZFHX4 | Zinc finger homeobox protein 4 | lhspfLERPMDMPYMIfdpnn | 2484 | 2494 | 135.1 | 0.000218728 |
| 267 | P31629 | HIVEP2 | Transcription factor HIVEP2 | irvtqLMTPSDSCEDTqmtey | 1983 | 1993 | 135.1 | 0.000219955 |
| 268 | Q05193 | DNM1 | Dynamin-1 | ellanLYSCGDQNTLMeesae | 705 | 715 | 135.0 | 0.000223069 |
| 269 | P10124 | SRGN | Serglycin | eqdyqLVDESDAFHDNlrsld | 122 | 132 | 135.0 | 0.000224107 |
| 270 | Q12955 | ANK3 | Ankyrin-3 | tsekeLCKMADSFFGTdtile | 2095 | 2105 | 135.0 | 0.000224295 |
| 271 | Q8IYB7 | DIS3L2 | DIS3-like exonuclease 2 | mapdtLQKQADHCNDRrmask | 732 | 742 | 134.9 | 0.000227409 |
| 272 | Q14789 | GOLGB1 | Golgin subfamily B member 1 | ktgqeLQSACDALKDQnskll | 391 | 401 | 134.9 | 0.000227598 |
| 273 | O75095 | MEGF6 | Multiple epidermal growth factor-like domains protein 6 | delpqLFQDDDVGADEeeael | 484 | 494 | 134.9 | 0.000227787 |
| 274 | Q9C0E4 | GRIP2 | Glutamate receptor-interacting protein 2 | rksgsLSETSDADEDPadalk | 750 | 760 | 134.9 | 0.000228825 |
| 275 | Q9C099 | LRRCC1 | Leucine-rich repeat and coiled-coil domain-containing protein 1 | qaenkLMDYIDELHKHaneke | 447 | 457 | 134.9 | 0.000229485 |
| 276 | Q86Y25 | ZNF354C | Zinc finger protein 354C | qekkpLRQMIDSHEKTisedg | 143 | 153 | 134.9 | 0.000229768 |
| 277 | Q9BX26 | SYCP2 | Synaptonemal complex protein 2 | llpkkLCKIEDADHHIhkmse | 1317 | 1327 | 134.9 | 0.000230806 |
| 279 | G9CGD6 | CNK3/IPCEF1 | CNK3/IPCEF1 fusion protein | gkprpLSMPADGNWMGivdpf | 432 | 442 | 134.9 | 0.000231278 |
| 278 | Q6P9H4 | CNKSR3 | Connector enhancer of kinase suppressor of ras 3 | gkprpLSMPADGNWMGivdpf | 432 | 442 | 134.9 | 0.000231278 |
| 280 | Q8WXI7 | MUC16 | Mucin-16 | vptptLAKITDMDTNLepvtr | 6259 | 6269 | 134.9 | 0.000231278 |
| 281 | Q9HCK1 | ZDBF2 | DBF4-type zinc finger-containing protein 2 | dsdipLYSVIDQPEVAvyeee | 798 | 808 | 134.9 | 0.00023175 |
| 283 | P18627 | LAG3 | Lymphocyte activation gene 3 protein | gggpdLLVTGDNGDFTlrled | 307 | 317 | 134.9 | 0.000232505 |
| 282 | Q13387 | MAPK8IP2 | C-Jun-amino-terminal kinase-interacting protein 2 | tddcgLGLSYDSDHCEkdsls | 46 | 56 | 134.9 | 0.000232505 |
| 284 | Q9Y2K5 | R3HDM2 | R3H domain-containing protein 2 | leqeiLEFINDNNNQFkkfpq | 180 | 190 | 134.9 | 0.000232599 |
| 285 | Q9UPX8 | SHANK2 | SH3 and multiple ankyrin repeat domains protein 2 | dkknmLIDIMDTSQQKsagll | 920 | 930 | 134.9 | 0.000233543 |
| 286 | Q6ZSZ6 | TSHZ1 | Teashirt homolog 1 | lresaLMDISDMVKNLtgrlt | 837 | 847 | 134.9 | 0.000233637 |
| 287 | Q15434 | RBMS2 | RNA-binding motif, single-stranded-interacting protein 2 | psdplLCKFADGGPKKrqnqg | 214 | 224 | 134.8 | 0.000237128 |
| 288 | Q16623 | STX1A | Syntaxin-1A | drtqeLRTAKDSDDDDdvavt | 8 | 18 | 134.8 | 0.000238261 |
| 289 | P29317 | EPHA2 | Ephrin type-A receptor 2 | eqlkpLKTYVDPHTYEdpnqa | 585 | 595 | 134.8 | 0.000239016 |
| 290 | Q8IWZ8 | SUGP1 | SURP and G-patch domain-containing protein 1 | keqqeMQQMYDMIMQHkramq | 451 | 461 | 134.8 | 0.000240431 |
| 291 | O15015 | ZNF646 | Zinc finger protein 646 | kapspLGVAGDAMEMVvdsvl | 993 | 1003 | 134.8 | 0.00024128 |
| 292 | Q9NV56 | MRGBP | MRG/MORF4L-binding protein | viwdhLSTMYDMQALHeseil | 81 | 91 | 134.8 | 0.00024128 |
| 293 | Q9ULK6 | RNF150 | RING finger protein 150 | lsdveLSTDQDCEEVKs | 427 | 437 | 134.8 | 0.00024128 |
| 294 | Q9Y577 | TRIM17 | E3 ubiquitin-protein ligase TRIM17 | stqgpLQMLQDMKEPLsrknn | 248 | 258 | 134.8 | 0.000241658 |
| 295 | P35908 | KRT2 | Keratin, type II cytoskeletal 2 epidermal | sqnseLNNMQDLVEDYkkkye | 254 | 264 | 134.8 | 0.000242696 |
| 296 | O75410 | TACC1 | Transforming acidic coiled-coil-containing protein 1 | srsspLKLEFDFTEDTgniea | 318 | 328 | 134.7 | 0.000243073 |
| 297 | Q9UKA4 | AKAP11 | A-kinase anchor protein 11 | sswssLGLEGDLYEDNlsfpt | 1768 | 1778 | 134.7 | 0.000243073 |
| 298 | P55289 | CDH12 | Cadherin-12 | ssidsLTTEADQDYDYltdwg | 757 | 767 | 134.7 | 0.000245998 |
| 299 | P20020 | ATP2B1 | Plasma membrane calcium-transporting ATPase 1 | ephipLIDDTDAEDDAptkrn | 1161 | 1171 | 134.7 | 0.000246659 |
| 300 | O43896 | KIF1C | Kinesin-like protein KIF1C | leqqrLYADSDSGDDSdkrsc | 670 | 680 | 134.7 | 0.000247319 |
| 301 | Q96J84 | KIRREL1 | Kin of IRRE-like protein 1 | vnrepLTMHSDREDDTasvst | 548 | 558 | 134.7 | 0.000247697 |
| 302 | Q8WYR1 | PIK3R5 | Phosphoinositide 3-kinase regulatory subunit 5 | sfvsgLSDGMDSGYVEdsees | 383 | 393 | 134.7 | 0.000247885 |
| 305 | O94986 | CEP152 | Centrosomal protein of 152 kDa | dtlplLVENADPEWKKrnmae | 1105 | 1115 | 134.7 | 0.000248074 |
| 303 | Q99715 | COL12A1 | Collagen alpha-1(XII) chain | vswksLYDDVDTGEKNlpeda | 942 | 952 | 134.7 | 0.000248074 |
| 306 | Q9HBR0 | SLC38A10 | Putative sodium-coupled neutral amino acid transporter 10 | gqaqaLEEAGDLPEDPqkvpe | 567 | 577 | 134.7 | 0.000248074 |
| 304 | Q9Y4F4 | TOGARAM1 | TOG array regulator of axonemal microtubules protein 1 | grsnhLAHGADTDWLLagnrt | 604 | 614 | 134.7 | 0.000248074 |
| 307 | Q8TCN5 | ZNF507 | Zinc finger protein 507 | ltranLGHYGDINLLDpdtsq | 515 | 525 | 134.7 | 0.000249867 |
| 308 | O76024 | WFS1 | Wolframin | vgkhyLQLAGDTDEELnscta | 111 | 121 | 134.7 | 0.000251566 |
| 309 | Q14999 | CUL7 | Cullin-7 | qpflaLMQSLDTPETNrtlhl | 669 | 679 | 134.6 | 0.000251943 |
| 310 | Q00975 | CACNA1B | Voltage-dependent N-type calcium channel subunit alpha-1B | asceaLYSEMDPEERLrfatt | 787 | 797 | 134.6 | 0.000252604 |
| 311 | Q9C0K0 | BCL11B | B-cell lymphoma/leukemia 11B | qcggsLGACYDKALDKdsppp | 85 | 95 | 134.6 | 0.00025317 |
| 312 | Q5VZK9 | CARMIL1 | F-actin-uncapping protein LRRC16A | geqngLMGRVDEGVDEfftkk | 1018 | 1028 | 134.6 | 0.000253547 |
| 313 | P48634 | PRRC2A | Protein PRRC2A | pvdpkLAWVGDVFTATpaepr | 789 | 799 | 134.6 | 0.000255057 |
| 314 | O60304 | ZNF500 | Zinc finger protein 500 | gpgiqLEDGGDGREDAplrme | 259 | 269 | 134.6 | 0.000255151 |
| 315 | Q96RN5 | MED15 | Mediator of RNA polymerase II transcription subunit 15 | eplrrMINKIDKNEDRkkdls | 549 | 559 | 134.6 | 0.000255434 |
| 316 | Q8TD57 | DNAH3 | Dynein heavy chain 3, axonemal | pevfgLHENADITKDNqetnq | 3801 | 3811 | 134.6 | 0.000257888 |
| 317 | A5PLL1 | ANKRD34B | Ankyrin repeat domain-containing protein 34B | fkdleLAGSNDDTWDPgspvr | 216 | 226 | 134.6 | 0.000257982 |
| 318 | Q7Z6Z7 | HUWE1 | E3 ubiquitin-protein ligase HUWE1 | seqlsLDELWDMLGEClkele | 3853 | 3863 | 134.6 | 0.000259869 |
| 319 | Q8NFP9 | NBEA | Neurobeachin | kdngpLITLADEKEDLpnsst | 1129 | 1139 | 134.5 | 0.000260058 |
| 320 | Q8TAB5 | C1orf216 | UPF0500 protein C1orf216 | dqqarLQESFDTILDNrkeli | 198 | 208 | 134.5 | 0.000261002 |
| 321 | Q03001 | DST | Dystonin | tiqlfLAKHGDKMTDEernel | 1499 | 1509 | 134.5 | 0.000261285 |
| 322 | Q13936 | CACNA1C | Voltage-dependent L-type calcium channel subunit alpha-1C | vttqeLADACDMTIEEmesaa | 2146 | 2156 | 134.5 | 0.0002627 |
| 323 | Q5T7N2 | L1TD1 | LINE-1 type transposase domain-containing protein 1 | nsvddLSSRMDILEERidsle | 652 | 662 | 134.5 | 0.000263832 |
| 324 | Q2M3C6 | TMEM266 | Transmembrane protein 266 | dssqtLGSSMDCSTAReepss | 405 | 415 | 134.5 | 0.000264021 |
| 325 | Q13023 | AKAP6 | A-kinase anchor protein 6 | eisssLGRLNDCYKEKsrlkk | 482 | 492 | 134.5 | 0.000264682 |
| 326 | Q14690 | PDCD11 | Protein RRP5 homolog | palppLAESSDSEEDEkphqa | 1553 | 1563 | 134.5 | 0.000264776 |
| 327 | Q8WZ42 | TTN | Titin | keerkLRMPYDVPEPRkykqt | 32895 | 32905 | 134.5 | 0.000265342 |
| 328 | Q9ULV0 | MYO5B | Unconventional myosin-Vb | keysqLEQRYDNLRDEmtiik | 1077 | 1087 | 134.5 | 0.000265342 |
| 329 | Q9Y2D4 | EXOC6B | Exocyst complex component 6B | cigptLRSVYDGEEHGrfmek | 37 | 47 | 134.5 | 0.00026572 |
| 330 | A6NKG5 | RTL1 | Retrotransposon-like protein 1 | sepseLQQAGDSDHSEtfyec | 595 | 605 | 134.5 | 0.000267135 |
| 331 | Q13637 | RAB32 | Ras-related protein Rab-32 | ipavlLANKCDQNKDSsqsps | 141 | 151 | 134.5 | 0.000267324 |
| 332 | O15055 | PER2 | Period circadian protein homolog 2 | ipslgLSEVSDTKEDEngspl | 1232 | 1242 | 134.4 | 0.000268173 |
| 333 | Q9HBD1 | RC3H2 | Roquin-2 | rphleLLANIDPNPDAvsptw | 356 | 366 | 134.4 | 0.000268456 |
| 334 | A2RUR9 | CCDC144A | Coiled-coil domain-containing protein 144A | dtgvvLLSGNDTLHDLcqsql | 198 | 208 | 134.4 | 0.00027006 |
| 335 | Q3MJ40 | CCDC144B | Coiled-coil domain-containing protein 144B | dtgvvLLSGNDTLHDLcqsql | 198 | 208 | 134.4 | 0.00027006 |
| 336 | Q6ZWC4 | | Putative uncharacterized protein LOC100128429 | cpshrLMHSTDGPLDPeplst | 195 | 205 | 134.4 | 0.00027006 |
| 337 | Q5VZ89 | DENND4C | DENN domain-containing protein 4C | sisnvLFSTQDPVEDAvfgea | 1008 | 1018 | 134.4 | 0.000270626 |
| 338 | Q92558 | WASF1 | Wiskott-Aldrich syndrome protein family member 1 | tlpipLQETYDVCEQPpplni | 121 | 131 | 134.4 | 0.000271193 |
| 339 | Q9ULG1 | INO80 | Chromatin-remodeling ATPase INO80 | lyahfMSRKRDMGHDGiqeei | 399 | 409 | 134.4 | 0.000271193 |
| 340 | Q5H8A4 | PIGG | GPI ethanolamine phosphate transferase 2 | plpnlLVLCGDHGMSEtgshg | 263 | 273 | 134.4 | 0.000271759 |
| 341 | Q6ZVT6 | CFAP20DC | Protein CFAP20DC | mspeeLSFILDLKEDNsvtsr | 375 | 385 | 134.4 | 0.000272702 |
| 342 | Q14151 | SAFB2 | Scaffold attachment factor B2 | gtdglLDSFCDSKEYVaaqlr | 156 | 166 | 134.4 | 0.000273174 |
| 343 | Q9Y6Q9 | NCOA3 | Nuclear receptor coactivator 3 | dskspLGFYCDQNPVEssmcq | 571 | 581 | 134.4 | 0.000273174 |
| 344 | O15535 | ZSCAN9 | Zinc finger and SCAN domain-containing protein 9 | tedreLVLRKDCPKIVephgk | 200 | 210 | 134.3 | 0.000276194 |
| 345 | P58417 | NXPH1 | Neurexophilin-1 | endtdLDLRYDTPEPYseqdl | 72 | 82 | 134.3 | 0.000278553 |
| 346 | Q659A1 | ICE2 | Little elongation complex subunit 2 | qkekqLVTGMDGGPEEcknkd | 470 | 480 | 134.3 | 0.000279496 |
| 347 | Q9ULL8 | SHROOM4 | Protein Shroom4 | kskgpLSQLCDTKEPVeetqe | 610 | 620 | 134.3 | 0.000280157 |

| | | | | | | | | |
|---|---|---|---|---|---|---|---|---|
| 348 | Q7Z6G8 | ANKS1B | Ankyrin repeat and sterile alpha motif domain-containing protein 1B | hrkriLASLGDRLHDDppqkp | 940 | 950 | 134.3 | 0.000282233 |
| 349 | Q5T5C0 | STXBP5 | Syntaxin-binding protein 5 | levrlLYEINDVETPEgeqpp | 548 | 558 | 134.3 | 0.000282421 |
| 350 | Q9H799 | CPLANE1 | Ciliogenesis and planar polarity effector 1 | sqesnLRGCGDVEDSNknlke | 2100 | 2110 | 134.3 | 0.00028261 |
| 351 | Q8WXI7 | MUC16 | Mucin-16 | ftitnLQYGEDMGHPGsrkfn | 12399 | 12409 | 134.3 | 0.000282799 |
| 352 | O14513 | NCKAP5 | Nck-associated protein 5 | tfvydLDSHVDADDDPstlal | 472 | 482 | 134.2 | 0.000284214 |
| 353 | P17480 | UBTF | Nucleolar transcription factor 1 | MNGEADCPTDLemaap | 1 | 11 | 134.2 | 0.000286196 |
| 354 | Q7Z570 | ZNF804A | Zinc finger protein 804A | qdhrsLVLQNDMKHMSqnqav | 739 | 749 | 134.2 | 0.000286196 |
| 355 | P0C7P4 | UQCRFS1P1 | Putative cytochrome b-c1 complex subunit Rieske-like protein 1 | eaaveLSQLRDPQHDLdrvkk | 201 | 211 | 134.2 | 0.000286951 |
| 356 | P47985 | UQCRFS1 | Cytochrome b-c1 complex subunit Rieske, mitochondrial | eaaveLSQLRDPQHDLdrvkk | 192 | 202 | 134.2 | 0.000286951 |
| 357 | Q8IZM8 | ZNF654 | Zinc finger protein 654 | nvfkpLTECGDDYEEEdedeg | 146 | 156 | 134.2 | 0.000286951 |
| 358 | Q96K83 | ZNF521 | Zinc finger protein 521 | inqcqLTDGVDVEDDPtcswp | 72 | 82 | 134.2 | 0.000287328 |
| 359 | Q96NW7 | LRRC7 | Leucine-rich repeat-containing protein 7 | psdynLGNYGDKPSDNsdlkt | 1237 | 1247 | 134.2 | 0.000289121 |
| 360 | Q8TED9 | AFAP1L1 | Actin filament-associated protein 1-like 1 | sfvesLFEEFDCDLSDlrdmp | 72 | 82 | 134.2 | 0.000289687 |
| 361 | Q8NEF9 | SRFBP1 | Serum response factor-binding protein 1 | gsdssLSGNSDGGEEFceeek | 243 | 253 | 134.2 | 0.00028997 |
| 362 | Q9HAU0 | PLEKHA5 | Pleckstrin homology domain-containing family A member 5 | qiqkeLWRIQDVMEGLskhkq | 762 | 772 | 134.2 | 0.000290536 |
| 363 | P67936 | TPM4 | Tropomyosin alpha-4 chain | rkiqaLQQQADEAEDRaqglq | 17 | 27 | 134.2 | 0.000291669 |
| 364 | Q15911 | ZFHX3 | Zinc finger homeobox protein 3 | dqlrvLRQYFDINNSPseeqi | 2160 | 2170 | 134.1 | 0.000293745 |
| 365 | O75376 | NCOR1 | Nuclear receptor corepressor 1 | lskeeLIQSMDRVDREiakve | 181 | 191 | 134.1 | 0.000293839 |
| 366 | Q9GZU2 | PEG3 | Paternally-expressed gene 3 protein | ynkekLCDFTDGRDAFmqsse | 684 | 694 | 134.1 | 0.000294971 |
| 367 | P51532 | SMARCA4 | Transcription activator BRG1 | eeefdLFMRMDLDRRReearn | 1315 | 1325 | 134.1 | 0.000295255 |
| 368 | D6RGH6 | MCIDAS | Multicilin | eadfnLQDFRDTVDDLisdss | 115 | 125 | 134.1 | 0.000298746 |
| 369 | Q9BRK5 | SDF4 | 45 kDa calcium-binding protein | vtaeeLESYMDPMNEYnalne | 303 | 313 | 134.0 | 0.000302237 |
| 370 | Q8N108 | MIER1 | Mesoderm induction early response protein 1 | seiedLAREGDMPIHEllsly | 58 | 68 | 134.0 | 0.000303747 |
| 371 | Q9ULI0 | ATAD2B | ATPase family AAA domain-containing protein 2B | geverLRMWTDTEFENmdmys | 217 | 227 | 134.0 | 0.000304313 |
| 373 | Q86VM9 | ZC3H18 | Zinc finger CCCH domain-containing protein 18 | vrdtvLEPYADPYYDYeierf | 364 | 374 | 134.0 | 0.000304974 |
| 372 | Q86YT6 | MIB1 | E3 ubiquitin-protein ligase MIB1 | htlsqLRQLQDMQDVGkvdaa | 715 | 725 | 134.0 | 0.000304974 |
| 374 | Q96BD0 | SLCO4A1 | Solute carrier organic anion transporter family member 4A1 | mpLHQLGDKPLTFpspns | 3 | 13 | 134.0 | 0.000305068 |
| 375 | Q6UN15 | FIP1L1 | Pre-mRNA 3'-end-processing factor FIP1 | ngvplLEVDLDSFEDKpwrkp | 153 | 163 | 134.0 | 0.000305162 |
| 376 | Q2VPK5 | CTU2 | Cytoplasmic tRNA 2-thiolation protein 2 | qrawgLQEIRDCLIEDsddea | 497 | 507 | 134.0 | 0.000305257 |
| 377 | O14686 | KMT2D | Histone-lysine N-methyltransferase 2D | gdefdLLAYTDPELDTgdkkd | 3049 | 3059 | 134.0 | 0.000306483 |
| 378 | P38398 | BRCA1 | Breast cancer type 1 susceptibility protein | psqeeLIKVVDVEEQQleesg | 1528 | 1538 | 134.0 | 0.000306955 |
| 379 | Q68DC2 | ANKS6 | Ankyrin repeat and SAM domain-containing protein 6 | pafqlLLRACDQGDTEtarrl | 13 | 23 | 134.0 | 0.000308748 |
| 380 | Q8N2Y8 | RUSC2 | Iporin | tgqgpLAQLMDPGPALpgspa | 643 | 653 | 134.0 | 0.000309503 |
| 381 | Q8NEM0 | MCPH1 | Microcephalin | trhdvLDDSCDGFKDLikphe | 620 | 630 | 134.0 | 0.000309503 |
| 382 | P0DPQ3 | PRR20G | Proline-rich protein 20G | gpgmyLVHLNDHGELGyqgql | 102 | 112 | 133.9 | 0.000310447 |
| 383 | Q96HS1 | PGAM5 | Serine/threonine-protein phosphatase PGAM5, mitochondrial | ngrvaLRTLGDTGFMPpdkit | 272 | 282 | 133.9 | 0.000310635 |
| 384 | P35613 | BSG | Basigin | lhienLNMEADPGQYRcngts | 290 | 300 | 133.9 | 0.00031073 |
| 385 | P21333 | FLNA | Filamin-A | ikvsgLGEKVDVGKDQeftvk | 979 | 989 | 133.9 | 0.000311201 |
| 386 | P00451 | F8 | Coagulation factor VIII | hfrpqLHHSGDMVFTPesglq | 860 | 870 | 133.9 | 0.000311296 |
| 387 | P34741 | SDC2 | Syndecan-2 | esraeLTSDKDMYLDNssiee | 24 | 34 | 133.9 | 0.000311296 |
| 388 | Q9BY07 | SLC4A5 | Electrogenic sodium bicarbonate cotransporter 4 | temdtLQHDGDQMEWKesarw | 120 | 130 | 133.9 | 0.000312994 |
| 389 | Q9NY46 | SCN3A | Sodium channel protein type 3 subunit alpha | kgridLPIKQDMIIDKlngns | 1938 | 1948 | 133.9 | 0.000312994 |
| 390 | Q8NF91 | SYNE1 | Nesprin-1 | sakeeLHRWSDMSGDSsatqk | 2863 | 2873 | 133.9 | 0.000313372 |
| 391 | Q9H501 | ESF1 | ESF1 homolog | qsvvqLIMTRDSDGYEnstdg | 216 | 226 | 133.9 | 0.000313938 |
| 392 | Q9BT88 | SYT11 | Synaptotagmin-11 | scidqLPIKMDYGEELrspit | 122 | 132 | 133.9 | 0.000316203 |
| 393 | A6NCI8 | C2orf78 | Uncharacterized protein C2orf78 | qvfhaLGKKIDMKTGFsssrt | 640 | 650 | 133.9 | 0.000317524 |
| 394 | O75157 | TSC22D2 | TSC22 domain family protein 2 | sdssvLTRSGDCIRHSstfdq | 195 | 205 | 133.9 | 0.000317712 |
| 395 | Q86XN8 | MEX3D | RNA-binding protein MEX3D | lsalgLGGAGDTDEEGaagdg | 71 | 81 | 133.9 | 0.000319033 |
| 396 | Q5TI25 | NBPF14 | Neuroblastoma breakpoint family member 14 | qqrvgLAVNMDEIEKYqevee | 330 | 340 | 133.8 | 0.000319505 |
| 397 | Q8N3P4 | VPS8 | Vacuolar protein sorting-associated protein 8 homolog | dmdkeLEFKNDLIDDKefdip | 45 | 55 | 133.8 | 0.000319977 |
| 398 | O14607 | UTY | Histone demethylase UTY | evrtqLLQPADENWDPtgtkk | 951 | 961 | 133.8 | 0.000320732 |
| 399 | O15550 | KDM6A | Lysine-specific demethylase 6A | evrtqLLQPADENWDPtgtkk | 1004 | 1014 | 133.8 | 0.000320732 |
| 400 | Q69YN4 | VIRMA | Protein virilizer homolog | ddpvpLPVSGDKEEDAphred | 201 | 211 | 133.8 | 0.000321959 |
| 401 | A6H8Y1 | BDP1 | Transcription factor TFIIIB component B'' homolog | prekiLDVIDDTIEMEtglka | 889 | 899 | 133.8 | 0.000322053 |
| 403 | Q6ZU65 | UBN2 | Ubinuclein-2 | eqrkkLIHTEDPFNDEhqerq | 159 | 169 | 133.8 | 0.00032243 |
| 402 | Q7Z736 | PLEKHH3 | Pleckstrin homology domain-containing family H member 3 | ygdgeLSGDGDEDEDEetfel | 29 | 39 | 133.8 | 0.00032243 |
| 404 | Q6MZP7 | LIN54 | Protein lin-54 homolog | perktLMHLADAAEVRvqqqt | 641 | 651 | 133.8 | 0.000322619 |
| 405 | Q6P3S6 | FBXO42 | F-box only protein 42 | qeetvLEGTMDQDEEPhpvle | 22 | 32 | 133.8 | 0.000323563 |
| 406 | Q92824 | PCSK5 | Proprotein convertase subtilisin/kexin type 5 | rthpdLMQNYDALASCdvngn | 181 | 191 | 133.8 | 0.00032394 |
| 407 | Q6P9F0 | CCDC62 | Coiled-coil domain-containing protein 62 | ensqeLIQMYDSKMEEskald | 309 | 319 | 133.8 | 0.000324318 |
| 408 | Q8N4L4 | SPEM1 | Spermatid maturation protein 1 | spavhLRCTMDPVMMTvsppp | 93 | 103 | 133.8 | 0.000325544 |
| 409 | Q9ULI3 | HEG1 | Protein HEG homolog 1 | fgltsLRWQNDSPTFGehqla | 410 | 420 | 133.8 | 0.000325639 |
| 410 | Q4J6C6 | PREPL | Prolyl endopeptidase-like | enekpLPENMDAFEKVrtkle | 86 | 96 | 133.8 | 0.000325922 |
| 411 | Q02930 | CREB5 | Cyclic AMP-responsive element-binding protein 5 | qlkqlLLTHKDCPITAmqkes | 435 | 445 | 133.8 | 0.000326865 |
| 412 | Q96M63 | ODAD1 | Outer dynein arm-docking complex subunit 1 | qqqkvLQQRMDKVHSEaerle | 346 | 356 | 133.8 | 0.00032696 |
| 413 | Q14692 | BMS1 | Ribosome biogenesis protein BMS1 homolog | dsemdLPAFADSDDDLerssa | 498 | 508 | 133.7 | 0.000329602 |
| 414 | Q9NXN4 | GDAP2 | Ganglioside-induced differentiation-associated protein 2 | vdvdtLPSWGDSCQDElnssd | 16 | 26 | 133.7 | 0.000330074 |
| 415 | Q9H074 | PAIP1 | Polyadenylate-binding protein-interacting protein 1 | engtdLSGAGDPYLDDiddem | 443 | 453 | 133.7 | 0.000330168 |
| 416 | Q8N187 | CARF | Calcium-responsive transcription factor | iipatLQWTTDSGNILketmt | 499 | 509 | 133.7 | 0.000332621 |
| 417 | A0AVK6 | E2F8 | Transcription factor E2F8 | dnrsgLPEAKDCIHEHlsgde | 90 | 100 | 133.7 | 0.000332999 |
| 418 | Q14980 | NUMA1 | Nuclear mitotic apparatus protein 1 | plessLDSLGDVFLDSgrktr | 1790 | 1800 | 133.7 | 0.000333471 |
| 419 | P08151 | GLI1 | Zinc finger protein GLI1 | giqdpLLGMLDGREDLereek | 211 | 221 | 133.7 | 0.00033432 |
| 420 | P41229 | KDM5C | Lysine-specific demethylase 5C | apleeLMMEGDLLEVTldenh | 1402 | 1412 | 133.7 | 0.000334792 |
| 421 | Q9BY66 | KDM5D | Lysine-specific demethylase 5D | apleeLMMEGDLLEVTldenh | 1386 | 1396 | 133.7 | 0.000334792 |
| 422 | O14526 | FCHO1 | F-BAR domain only protein 1 | srenyLNRCMDQERLRrests | 142 | 152 | 133.7 | 0.00033583 |
| 424 | Q641Q2 | WASHC2A | WASH complex subunit 2A | ppptgLFDDDDGDDDDdffsa | 450 | 460 | 133.7 | 0.000336113 |
| 426 | Q8N205 | SYNE4 | Nesprin-4 | tldqdLEVEGDSDWPGpggvw | 218 | 228 | 133.7 | 0.000336113 |
| 423 | Q8N6G2 | TEX26 | Testis-expressed protein 26 | lchhnLQPTDDPNWDSyattm | 18 | 28 | 133.7 | 0.000336113 |
| 425 | Q9Y4E1 | WASHC2C | WASH complex subunit 2C | ppptgLFDDDDGDDDDdffsa | 450 | 460 | 133.7 | 0.000336113 |
| 427 | Q9NPI1 | BRD7 | Bromodomain-containing protein 7 | eriqsLKQSIDFMADLqktrk | 240 | 250 | 133.7 | 0.000338377 |
| 428 | O43525 | KCNQ3 | Potassium voltage-gated channel subfamily KQT member 3 | dmgkkLDFLVDMHMQHmerlq | 636 | 646 | 133.7 | 0.000338755 |
| 429 | Q8TC05 | MDM1 | Nuclear protein MDM1 | hstkvLSENVDNGLDRllrkk | 159 | 169 | 133.6 | 0.000338849 |
| 431 | P04259 | KRT6B | Keratin, type II cytoskeletal 6B | rldseLRNMQDLVEDLknkye | 239 | 249 | 133.6 | 0.000339604 |
| 430 | P48668 | KRT6C | Keratin, type II cytoskeletal 6C | rldseLRNMQDLVEDLknkye | 239 | 249 | 133.6 | 0.000339604 |
| 432 | Q15058 | KIF14 | Kinesin-like protein KIF14 | ssrlkLHLKSDMSECEnddpl | 38 | 48 | 133.6 | 0.000340359 |
| 433 | P67870 | CSNK2B | Casein kinase II subunit beta | hyrqaLDMILDLEPDEeledn | 50 | 60 | 133.6 | 0.000340453 |
| 434 | Q6PF04 | ZNF613 | Zinc finger protein 613 | ksdfpLRQNHDTFDLHgkilk | 122 | 132 | 133.6 | 0.000340548 |
| 435 | Q01814 | ATP2B2 | Plasma membrane calcium-transporting ATPase 2 | qphipLIDDTDLEEDAalkqn | 1184 | 1194 | 133.6 | 0.000343001 |

| | | | | | | | | |
|---|---|---|---|---|---|---|---|---|
| 436 | O94885 | SASH1 | SAM and SH3 domain-containing protein 1 | ltaveLLQEYDSNSDQsgsqe | 693 | 703 | 133.6 | 0.000345926 |
| 437 | Q9Y6H5 | SNCAIP | Synphilin-1 | ktipeLCRRCDTQNEDrsvss | 31 | 41 | 133.6 | 0.000346115 |
| 438 | Q5KU26 | COLEC12 | Collectin-12 | dtlekLQASGDALVDRqsqlk | 139 | 149 | 133.5 | 0.000348002 |
| 440 | Q6UUV9 | CRTC1 | CREB-regulated transcription coactivator 1 | eltssLAGVGDVSFDSdsqfp | 581 | 591 | 133.5 | 0.00034838 |
| 441 | Q9BT81 | SOX7 | Transcription factor SOX-7 | lsqveLLGDMDRNEFDqylnt | 323 | 333 | 133.5 | 0.00034838 |
| 439 | Q9Y2G9 | SBNO2 | Protein strawberry notch homolog 2 | ssvdsLSDIVDTPDFLpadsl | 114 | 124 | 133.5 | 0.00034838 |
| 442 | Q99973 | TEP1 | Telomerase protein component 1 | aqeatLGRWFDSEEKKgaetq | 189 | 199 | 133.5 | 0.000348663 |
| 443 | Q96RD6 | PANX2 | Pannexin-2 | qeggfLSQAEDCGLGLapapi | 547 | 557 | 133.5 | 0.000350267 |
| 444 | P30530 | AXL | Tyrosine-protein kinase receptor UFO | epdeiLYVNMDEGGGYpeppg | 820 | 830 | 133.5 | 0.000351399 |
| 445 | Q9H892 | TTC12 | Tetratricopeptide repeat protein 12 | tekrlLLMEEDQEEDEcrttl | 46 | 56 | 133.5 | 0.000351399 |
| 446 | Q9UBK2 | PPARGC1A | Peroxisome proliferator-activated receptor gamma coactivator 1-alpha | tkrpsLRLFGDHDYCQsinsk | 378 | 388 | 133.5 | 0.000351777 |
| 447 | Q96BT7 | ALKBH8 | Alkylated DNA repair protein alkB homolog 8 | eeekmLLESVDWTEDTdnqns | 152 | 162 | 133.5 | 0.000356212 |
| 448 | Q6ZSB3 | LINC00299 | Putative uncharacterized protein encoded by LINC00299 | kdgnsLHRHGDQAWGKhrrqn | 56 | 66 | 133.4 | 0.000356683 |
| 449 | O43149 | ZZEF1 | Zinc finger ZZ-type and EF-hand domain-containing protein 1 | keinaLAEHGDLELDErgdre | 2420 | 2430 | 133.4 | 0.000356966 |
| 450 | Q92564 | DCUN1D4 | DCN1-like protein 4 | htlnkLNLTEDIGQDDhqtgs | 32 | 42 | 133.4 | 0.000357061 |
| 451 | P23560 | BDNF | Brain-derived neurotrophic factor | agsrgLTSLADTFEHVieell | 56 | 66 | 133.4 | 0.000357249 |
| 452 | Q8TEW0 | PARD3 | Partitioning defective 3 homolog | aehenLFRENDCIVRIndgdl | 317 | 327 | 133.4 | 0.000357533 |
| 453 | Q86UU1 | PHLDB1 | Pleckstrin homology-like domain family B member 1 | siekdLQEIMDSLVLEepgaa | 197 | 207 | 133.4 | 0.000357721 |
| 454 | Q13835 | PKP1 | Plakophilin-1 | ksdkmMNNNYDCPLPEeetnp | 522 | 532 | 133.4 | 0.000359514 |
| 455 | Q8IUM7 | NPAS4 | Neuronal PAS domain-containing protein 4 | wkcqeLDFLADPDNMFleetp | 694 | 704 | 133.4 | 0.00036159 |
| 456 | P0C671 | BNIP5 | Protein BNIP5 | mivelLKRVGDQWEEEqslas | 254 | 264 | 133.4 | 0.000361684 |
| 457 | Q9C0D6 | FHDC1 | FH2 domain-containing protein 1 | kdprpLFCISDTTDCSltldc | 770 | 780 | 133.4 | 0.000362251 |
| 458 | P19404 | NDUFV2 | NADH dehydrogenase [ubiquinone] flavoprotein 2, mitochondrial | gaggaLFVHRDTPENNpdtpf | 38 | 48 | 133.4 | 0.000362628 |
| 459 | Q9HBH7 | BEX1 | Protein BEX1 | nkgepLALPLDAGEYCvprgn | 34 | 44 | 133.4 | 0.000364232 |
| 460 | Q92828 | CORO2A | Coronin-2A | ekktwLTNGFDVFECPppkte | 465 | 475 | 133.4 | 0.000364704 |
| 461 | P16144 | ITGB4 | Integrin beta-4 | ttsgtLSTHMDQQFFQt | 1811 | 1821 | 133.3 | 0.000365742 |
| 462 | Q13253 | NOG | Noggin | llrslLGGHYDPGFMAtsppe | 70 | 80 | 133.3 | 0.000366025 |
| 463 | Q9H0J4 | QRICH2 | Glutamine-rich protein 2 | anrehLEMEIDVKADKsalat | 1315 | 1325 | 133.3 | 0.000367063 |
| 465 | O14640 | DVL1 | Segment polarity protein dishevelled homolog DVL-1 | epgdmLLQVNDVNFENmsndd | 301 | 311 | 133.3 | 0.000367157 |
| 464 | P54792 | DVL1P1 | Putative segment polarity protein dishevelled homolog DVL1P1 | epgdmLLQVNDVNFENmsndd | 301 | 311 | 133.3 | 0.000367157 |
| 466 | Q9NRE2 | TSHZ2 | Teashirt homolog 2 | pqkhaLSDIADMVKVLpkatt | 792 | 802 | 133.3 | 0.000367723 |
| 467 | O60282 | KIF5C | Kinesin heavy chain isoform 5C | eeissLYRQLDDKDDEinqqs | 424 | 434 | 133.3 | 0.000369045 |
| 468 | Q96N16 | JAKMIP1 | Janus kinase and microtubule-interacting protein 1 | gdmveLMGVQDQHMDErdvrr | 274 | 284 | 133.3 | 0.000369233 |
| 469 | Q96Q06 | PLIN4 | Perilipin-4 | mlqneLEGLGDIFHPMnaeeq | 1142 | 1152 | 133.3 | 0.000369799 |
| 470 | O60500 | NPHS1 | Nephrin | npepsLMWYKDSRTVTesrlp | 476 | 486 | 133.3 | 0.000370743 |
| 471 | Q8WWN8 | ARAP3 | Arf-GAP with Rho-GAP domain, ANK repeat and PH domain-containing prote | latvhLEQYADTFRRHglata | 18 | 28 | 133.3 | 0.000370743 |
| 472 | Q7Z6I5 | SPATA12 | Spermatogenesis-associated protein 12 | tcgstLEKSGDTWEMKaldss | 12 | 22 | 133.3 | 0.000371687 |
| 473 | Q9NYI0 | PSD3 | PH and SEC7 domain-containing protein 3 | iknekLEWAVDDEEKKkspse | 736 | 746 | 133.3 | 0.000372536 |
| 474 | P54257 | HAP1 | Huntingtin-associated protein 1 | lrdelLQLYSDSDEEDedeee | 253 | 263 | 133.3 | 0.000373574 |
| 475 | Q7Z570 | ZNF804A | Zinc finger protein 804A | nkstvLDMSNDCISVQattee | 363 | 373 | 133.3 | 0.00037414 |
| 476 | Q8IWU2 | LMTK2 | Serine/threonine-protein kinase LMTK2 | yppalLTTDMDNPERTgpels | 572 | 582 | 133.2 | 0.000374329 |
| 477 | Q8N157 | AHI1 | Jouberin | aetstLTISGDTVEGEqkkes | 252 | 262 | 133.2 | 0.000376216 |
| 478 | Q9BYB0 | SHANK3 | SH3 and multiple ankyrin repeat domains protein 3 | lesihLGEHRDRFEDHeiega | 1682 | 1692 | 133.2 | 0.000376971 |
| 479 | Q6NUN7 | JHY | Jhy protein homolog | fsyqqLHTLSDMDLNNlnels | 485 | 495 | 133.2 | 0.000379424 |
| 480 | Q8NBV8 | SYT8 | Synaptotagmin-8 | asgqpLQHWADMLAHArrpia | 356 | 366 | 133.2 | 0.000379707 |
| 481 | O95071 | UBR5 | E3 ubiquitin-protein ligase UBR5 | rygsaLASAGDPGHPNhplha | 1859 | 1869 | 133.2 | 0.000380934 |
| 482 | Q9BT81 | SOX7 | Transcription factor SOX-7 | kqakrLCKRVDPGFLLsslsr | 125 | 135 | 133.2 | 0.000381595 |
| 483 | B1APH4 | ZNF487 | Putative zinc finger protein 487 | psqshLDCCIDDDLMEkrqen | 108 | 118 | 133.2 | 0.000382727 |
| 485 | O75362 | ZNF217 | Zinc finger protein 217 | aqtknLKRFFDGAKDVtgspp | 557 | 567 | 133.2 | 0.00038367 |
| 484 | Q8NB14 | USP38 | Ubiquitin carboxyl-terminal hydrolase 38 | nptsgLWINGDPPLQKelmda | 964 | 974 | 133.2 | 0.00038367 |
| 486 | O60673 | REV3L | DNA polymerase zeta catalytic subunit | ggsqlLFKQKDMPLMGsavdh | 1261 | 1271 | 133.2 | 0.000384048 |
| 487 | Q8IZ41 | RASEF | Ras and EF-hand domain-containing protein | cfdsgLSTLRDPNEYDsevey | 440 | 450 | 133.2 | 0.000384803 |
| 489 | Q14789 | GOLGB1 | Golgin subfamily B member 1 | pldpeLHQESDMEFNNttqed | 34 | 44 | 133.2 | 0.000385558 |
| 488 | Q8WUY3 | PRUNE2 | Protein prune homolog 2 | sqlemLGFSADSTEWWkaspq | 1820 | 1830 | 133.2 | 0.000385558 |
| 490 | Q8N187 | CARF | Calcium-responsive transcription factor | vamdeLVEVGDVEDTGnlegt | 648 | 658 | 133.2 | 0.000385746 |
| 491 | Q92793 | CREBBP | CREB-binding protein | alsseLSLVGDTTGDTlekfv | 2424 | 2434 | 133.2 | 0.000386407 |
| 493 | O43290 | SART1 | U4/U6.U5 tri-snRNP-associated protein 1 | rqlqqLQQLRDSGEKVveivk | 515 | 525 | 133.1 | 0.000386596 |
| 492 | Q5SYE7 | NHSL1 | NHS-like protein 1 | prlgaLGPAGDMNGTFlyqrg | 348 | 358 | 133.1 | 0.000386596 |
| 494 | Q4LE39 | ARID4B | AT-rich interactive domain-containing protein 4B | kdievLSEDTDYEEDEvtkkr | 789 | 799 | 133.1 | 0.000387256 |
| 495 | Q68CZ2 | TNS3 | Tensin-3 | tlvpdLGLGMDGPYERertfg | 541 | 551 | 133.1 | 0.000387351 |
| 496 | Q9NR83 | SLC2A4RG | SLC2A4 regulator | fyyteLDVGVDTLTDGlsslt | 246 | 256 | 133.1 | 0.000387634 |
| 497 | O95447 | LCA5L | Lebercilin-like protein | yshriLKNLHDTEDYPkvsst | 342 | 352 | 133.1 | 0.000388955 |
| 498 | P49454 | CENPF | Centromere protein F | lavadLEKQRDCSQDLlkkre | 563 | 573 | 133.1 | 0.000389238 |
| 499 | A8MWX3 | WASH4P | Putative WAS protein family homolog 4 | giandLMYIADLGPGIapsap | 273 | 283 | 133.1 | 0.000390464 |
| 501 | C4AMC7 | WASH3P | Putative WAS protein family homolog 3 | gitndLMYIADLGPGIapsap | 258 | 268 | 133.1 | 0.000390464 |
| 500 | Q6VEQ5 | WASH2P | WAS protein family homolog 2 | giandLMYIADLGPGIapsap | 260 | 270 | 133.1 | 0.000390464 |
| 503 | A0A1B0GTK | FAM236D | Protein FAM236D | kdpeeLVAVSDTAEDPssgtg | 28 | 38 | 133.1 | 0.000390653 |
| 502 | P0DP71 | FAM236C | Protein FAM236C | kdpeeLVAVSDTAEDPssgtg | 28 | 38 | 133.1 | 0.000390653 |
| 504 | O00401 | WASL | Neural Wiskott-Aldrich syndrome protein | nldpeLKNLFDMCGISeaqlk | 232 | 242 | 133.1 | 0.000390936 |
| 506 | Q16829 | DUSP7 | Dual specificity protein phosphatase 7 | lglggLRISSDCSDGEsdrel | 210 | 220 | 133.1 | 0.000392446 |
| 505 | Q8TF72 | SHROOM3 | Protein Shroom3 | yktlrLVVRRDVCTDPghadt | 104 | 114 | 133.1 | 0.000392446 |
| 507 | O15027 | SEC16A | Protein transport protein Sec16A | sekagLSGQADFDDFCsspgl | 394 | 404 | 133.1 | 0.000395088 |
| 508 | Q12756 | KIF1A | Kinesin-like protein KIF1A | skleaLQKQMDSRYYPevnee | 674 | 684 | 133.1 | 0.000395937 |
| 509 | Q9UI42 | CPA4 | Carboxypeptidase A4 | vtiedLQALLDNEDDEmqhne | 94 | 104 | 133.1 | 0.00039622 |
| 510 | A2CJ06 | DYTN | Dystrotelin | elqelLSKLMDAFNLEtpsgp | 531 | 541 | 133.0 | 0.00039707 |
| 511 | Q8N8J7 | FAM241A | Uncharacterized protein FAM241A | csageLLRGGDGGERDedgda | 7 | 17 | 133.0 | 0.000397541 |
| 512 | Q14005 | IL16 | Pro-interleukin-16 | snrksLSQQLDCPAGKaagts | 176 | 186 | 133.0 | 0.000398579 |
| 513 | Q71RG4 | TMUB2 | Transmembrane and ubiquitin-like domain-containing protein 2 | litvrLKFLNDTEELAvarpe | 178 | 188 | 133.0 | 0.000399429 |
| 514 | Q96QP1 | ALPK1 | Alpha-protein kinase 1 | deegqLLDSMDVPCTNghgsh | 857 | 867 | 133.0 | 0.000399523 |
| 515 | Q8NE31 | FAM13C | Protein FAM13C | gtsrlLYHITDGDNPLlsprc | 226 | 236 | 133.0 | 0.000400278 |
| 516 | Q9UIF9 | BAZ2A | Bromodomain adjacent to zinc finger domain protein 2A | dvssfLETTADVEEITgeglt | 519 | 529 | 133.0 | 0.000401788 |
| 517 | Q13367 | AP3B2 | AP-3 complex subunit beta-2 | krhddLKEMLDTNKDSlklea | 42 | 52 | 133.0 | 0.000402165 |
| 518 | Q8NI22 | MCFD2 | Multiple coagulation factor deficiency protein 2 | msedeLINIIDGVLRDddknn | 117 | 127 | 133.0 | 0.000402543 |
| 519 | O95747 | OXSR1 | Serine/threonine-protein kinase OSR1 | gssgrLHKTEDGGWEWsddef | 328 | 338 | 133.0 | 0.000403486 |
| 520 | Q86VP3 | PACS2 | Phosphofurin acidic cluster sorting protein 2 | eaeedLDLLYDTLDMEhpsds | 286 | 296 | 133.0 | 0.000403581 |
| 521 | Q92574 | TSC1 | Hamartin | aaeerLDCCNDGCSDSmvghn | 993 | 1003 | 133.0 | 0.000403581 |
| 522 | Q8IUD2 | ERC1 | ELKS/Rab6-interacting/CAST family member 1 | qtqnrMKLMADNYEDDhfkss | 980 | 990 | 133.0 | 0.000404147 |
| 523 | Q68DE3 | USF3 | Basic helix-loop-helix domain-containing protein USF3 | pqkssLSDQMDHPDFSsenpk | 1022 | 1032 | 133.0 | 0.000404807 |

| | | | | | | | | |
|---|---|---|---|---|---|---|---|---|
| 524 | O75153 | CLUH | Clustered mitochondria protein homolog | MLLNGDCPESLkkeaa | 1 | 11 | 133.0 | 0.000404902 |
| 525 | Q16549 | PCSK7 | Proprotein convertase subtilisin/kexin type 7 | lesvpLCSSKDPDEVEtesrg | 725 | 735 | 133.0 | 0.000405185 |
| 526 | Q6RFH5 | WDR74 | WD repeat-containing protein 74 | qlnclLLSGRDNWEDEpqepq | 318 | 328 | 133.0 | 0.000405656 |
| 527 | Q15468 | STIL | SCL-interrupting locus protein | mkrygLLQSSDNSEDEeeppd | 1128 | 1138 | 133.0 | 0.00040594 |
| 528 | Q9Y618 | NCOR2 | Nuclear receptor corepressor 2 | iirkaLMGKYDQWEESpplsa | 2347 | 2357 | 133.0 | 0.000406034 |
| 529 | O75339 | CILP | Cartilage intermediate layer protein 1 | akeiaLGRCFDGTSDGssrim | 1092 | 1102 | 132.9 | 0.000407544 |
| 530 | P00451 | F8 | Coagulation factor VIII | lailtLEMTGDQREVGslgts | 1469 | 1479 | 132.9 | 0.000407921 |
| 531 | Q9Y4C1 | KDM3A | Lysine-specific demethylase 3A | qeeevLKTIQDGDSDEltikr | 1147 | 1157 | 132.9 | 0.00040962 |
| 532 | Q14C86 | GAPVD1 | GTPase-activating protein and VPS9 domain-containing protein 1 | wstdvLGSDFDPNIDEdrlqe | 660 | 670 | 132.9 | 0.000410469 |
| 533 | Q6ZUB1 | SPATA31E1 | Spermatogenesis-associated protein 31E1 | vseiaLIVQVDSEEQLpgrap | 1102 | 1112 | 132.9 | 0.000410563 |
| 535 | Q14681 | KCTD2 | BTB/POZ domain-containing protein KCTD2 | sflcrLCCQEDPELDSdkdet | 101 | 111 | 132.9 | 0.000410941 |
| 534 | Q5SYB0 | FRMPD1 | FERM and PDZ domain-containing protein 1 | aqeqkLFVELDLDPDFflgkq | 1200 | 1210 | 132.9 | 0.000410941 |
| 536 | Q12770 | SCAP | Sterol regulatory element-binding protein cleavage-activating protein | gdqpdLTCLIDTNFSAqprss | 873 | 883 | 132.9 | 0.000411129 |
| 537 | Q6KC79 | NIPBL | Nipped-B-like protein | klstlLNHNNDTEEEErlwrd | 1283 | 1293 | 132.9 | 0.000411979 |
| 538 | Q6RI45 | BRWD3 | Bromodomain and WD repeat-containing protein 3 | eqdlrLINEGDVPHLPvnray | 662 | 672 | 132.9 | 0.000412639 |
| 539 | P43307 | SSR1 | Translocon-associated protein subunit alpha | gprglLAVAQDLTEDEetved | 29 | 39 | 132.9 | 0.000413771 |
| 540 | Q6ZMI0 | PPP1R21 | Protein phosphatase 1 regulatory subunit 21 | qslrkLQQEMDSLTFRnlqla | 55 | 65 | 132.9 | 0.000414621 |
| 542 | Q86YV9 | HPS6 | Hermansky-Pudlak syndrome 6 protein | lsgpvLSPYEDILWDPstppp | 754 | 764 | 132.9 | 0.000415187 |
| 541 | Q96AV8 | E2F7 | Transcription factor E2F7 | tdslqLDVVGDSAVDEfekqr | 124 | 134 | 132.9 | 0.000415187 |
| 543 | Q04637 | EIF4G1 | Eukaryotic translation initiation factor 4 gamma 1 | qkpegLPHISDVVLDKanktp | 633 | 643 | 132.9 | 0.000415376 |
| 544 | Q8IWZ3 | ANKHD1 | Ankyrin repeat and KH domain-containing protein 1 | gksqeLNFVMDVNSSKypsll | 1614 | 1624 | 132.9 | 0.000415942 |
| 546 | Q7Z403 | TMC6 | Transmembrane channel-like protein 6 | maqpLAFILDVPETPgdqgq | 5 | 15 | 132.9 | 0.000416697 |
| 545 | Q9P2P5 | HECW2 | E3 ubiquitin-protein ligase HECW2 | grqdsLNDYLDAIEHNghsrp | 416 | 426 | 132.9 | 0.000416697 |
| 547 | Q92576 | PHF3 | PHD finger protein 3 | stvvgLDDIMDEGVVKesgnd | 85 | 95 | 132.9 | 0.00041698 |
| 548 | Q92786 | PROX1 | Prospero homeobox protein 1 | tanqrLQCFGDVIIPNpldtf | 409 | 419 | 132.9 | 0.00041698 |
| 549 | Q96P44 | COL21A1 | Collagen alpha-1(XXI) chain | fdvqkLRIYCDPEQNNretac | 406 | 416 | 132.9 | 0.000417168 |
| 550 | Q9C073 | FAM117A | Protein FAM117A | qgeeeLLRILDIPDGHrapap | 234 | 244 | 132.9 | 0.00041764 |
| 552 | P54132 | BLM | Bloom syndrome protein | sspdsLSTINDWDDMDdfdts | 148 | 158 | 132.8 | 0.000418112 |
| 551 | Q9NRR4 | DROSHA | Ribonuclease 3 | vgtsrLRDLYDKFEEElgsrq | 441 | 451 | 132.8 | 0.000418112 |
| 553 | Q15652 | JMJD1C | Probable JmjC domain-containing histone demethylation protein 2C | esqspLHWLADLAEQKareek | 2009 | 2019 | 132.8 | 0.00041915 |
| 554 | Q58WW2 | DCAF6 | DDB1- and CUL4-associated factor 6 | trdsaLQDTDDSDDDPvlipg | 651 | 661 | 132.8 | 0.000419716 |
| 555 | Q5T089 | MORN1 | MORN repeat-containing protein 1 | gpgqmLFQNGDKYDGDwvrdr | 148 | 158 | 132.8 | 0.000419716 |
| 556 | P00533 | EGFR | Epidermal growth factor receptor | nfyraLMDEEDMDDVVdadey | 1001 | 1011 | 132.8 | 0.000419999 |
| 557 | Q12840 | KIF5A | Kinesin heavy chain isoform 5A | eeirrLYKQLDDKDDEinqqs | 424 | 434 | 132.8 | 0.000421792 |
| 558 | Q9Y2J4 | AMOTL2 | Angiomotin-like protein 2 | keeqiLALEADMTKWEqkyle | 560 | 570 | 132.8 | 0.000421792 |
| 559 | Q01955 | COL4A3 | Collagen alpha-3(IV) chain | gipgsLGKCGDPGLPGpdgep | 679 | 689 | 132.8 | 0.000423113 |
| 560 | Q6ZV29 | PNPLA7 | Patatin-like phospholipase domain-containing protein 7 | kkpprLQESCDSDHGGgrpaa | 330 | 340 | 132.8 | 0.000423585 |
| 561 | Q8NET6 | CHST13 | Carbohydrate sulfotransferase 13 | ekrspLQKLYDLDQDPrstla | 52 | 62 | 132.8 | 0.000424057 |
| 563 | P98160 | HSPG2 | Basement membrane-specific heparan sulfate proteoglycan core protein | qlrlhLVSPADSGEYVcraas | 2980 | 2990 | 132.8 | 0.000425283 |
| 562 | Q15796 | SMAD2 | Mothers against decapentaplegic homolog 2 | tsdqqLNQSMDTGSPAelspt | 237 | 247 | 132.8 | 0.000425283 |
| 564 | Q9NPF7 | IL23A | Interleukin-23 subunit alpha | wsahpLVGHMDLREEGdeett | 50 | 60 | 132.8 | 0.000427265 |
| 565 | Q5FWF5 | ESCO1 | N-acetyltransferase ESCO1 | vkrkvLEVKSDSKEDEnlvin | 180 | 190 | 132.8 | 0.000428114 |
| 566 | Q86V81 | ALYREF | THO complex subunit 4 | MADKMDMSLDDiikln | 1 | 11 | 132.8 | 0.000428114 |
| 567 | Q96JQ0 | DCHS1 | Protocadherin-16 | sgtlrLAHALDCETQArhqlv | 2661 | 2671 | 132.8 | 0.00042868 |
| 568 | O95502 | NPTXR | Neuronal pentraxin receptor | avptgLHSKMDQLEGQllaqv | 231 | 241 | 132.7 | 0.000429341 |
| 569 | Q86Y26 | NUTM1 | NUT family member 1 | epvniLDVKDDCGLQLrvsed | 904 | 914 | 132.7 | 0.000429624 |
| 570 | Q15572 | TAF1C | TATA box-binding protein-associated factor RNA polymerase I subunit C | sssfsLSGHVDPSEDTssphs | 762 | 772 | 132.7 | 0.000429907 |
| 571 | Q8NET4 | RTL9 | Retrotransposon Gag-like protein 9 | tmstpLMRTSDPGERPslltr | 771 | 781 | 132.7 | 0.000429907 |
| 573 | P10071 | GLI3 | Transcriptional activator GLI3 | slktrLALLGDALEPGvalpp | 957 | 967 | 132.7 | 0.00043019 |
| 572 | Q14197 | MRPL58 | Peptidyl-tRNA hydrolase ICT1, mitochondrial | nladcLQKIRDMITEAsqtpk | 151 | 161 | 132.7 | 0.00043019 |
| 574 | Q96RU3 | FNBP1 | Formin-binding protein 1 | semkvLATDFDDEFDDeeplp | 535 | 545 | 132.7 | 0.000430285 |
| 575 | P12830 | CDH1 | Cadherin-1 | dpmeiLITVTDQNDNKpeftq | 249 | 259 | 132.7 | 0.000430379 |
| 576 | P98082 | DAB2 | Disabled homolog 2 | kykakLIGIDDVPDARgdkms | 54 | 64 | 132.7 | 0.000431794 |
| 577 | P0C2Y1 | NBPF7 | Putative neuroblastoma breakpoint family member 7 | hlvhkLSPENDTDEDEndktk | 182 | 192 | 132.7 | 0.000433115 |
| 578 | Q7Z6G3 | NECAB2 | N-terminal EF-hand calcium-binding protein 2 | keledLFHTIDSDNTNhvdtk | 102 | 112 | 132.7 | 0.000434342 |
| 579 | Q9Y5G5 | PCDHGA8 | Protocadherin gamma-A8 | ekndsLLTSVDFHEYKneadh | 791 | 801 | 132.7 | 0.000434814 |
| 580 | Q0VD83 | APOBR | Apolipoprotein B receptor | alegvLGQGWDSKEKEeaaag | 786 | 796 | 132.7 | 0.000435003 |
| 581 | Q9UGR2 | ZC3H7B | Zinc finger CCCH domain-containing protein 7B | nkildMQQTYDMWLKKhnpgk | 798 | 808 | 132.7 | 0.000435758 |
| 582 | Q9BY84 | DUSP16 | Dual specificity protein phosphatase 16 | qalsgLHLSADRLEDSnklkr | 383 | 393 | 132.7 | 0.000436229 |
| 583 | Q6P444 | MTFR2 | Mitochondrial fission regulator 2 | dipnmLDVLKDMNKVKlraie | 274 | 284 | 132.7 | 0.000437079 |
| 584 | Q92754 | TFAP2C | Transcription factor AP-2 gamma | mLWKITDNVKYEedced | 2 | 12 | 132.7 | 0.000437456 |
| 585 | Q01484 | ANK2 | Ankyrin-2 | tksaaLLLQNDHNADVqskmm | 210 | 220 | 132.7 | 0.0004384 |
| 586 | P78362 | SRPK2 | SRSF protein kinase 2 | ngpfsLEQQLDDEDDDeedcp | 394 | 404 | 132.7 | 0.000438871 |
| 588 | P40425 | PBX2 | Pre-B-cell leukemia transcription factor 2 | ggsfnLSGSGDMFLGMpglng | 349 | 359 | 132.7 | 0.000440476 |
| 587 | Q3SY00 | TSGA10IP | Testis-specific protein 10-interacting protein | klhrqLQRDLDCGPQKlpwkt | 328 | 338 | 132.7 | 0.000440476 |
| 590 | Q6NUM6 | TYW1B | S-adenosyl-L-methionine-dependent tRNA 4-demethylwyosine synthase TY | edhqsLNSIVDVEDLGkimdh | 262 | 272 | 132.6 | 0.000440853 |
| 589 | Q9NV66 | TYW1 | S-adenosyl-L-methionine-dependent tRNA 4-demethylwyosine synthase TY | edhqsLNSIVDVEDLGkimdh | 308 | 318 | 132.6 | 0.000440853 |
| 591 | O14654 | IRS4 | Insulin receptor substrate 4 | ggrfrLYFCVDRGATKeckea | 655 | 665 | 132.6 | 0.000441608 |
| 592 | Q9H0K1 | SIK2 | Serine/threonine-protein kinase SIK2 | lalseLPGLFDCEMLDavdpq | 903 | 913 | 132.6 | 0.000442457 |
| 593 | O43683 | BUB1 | Mitotic checkpoint serine/threonine-protein kinase BUB1 | feeqlLKQKMDELHKKlhqvv | 289 | 299 | 132.6 | 0.00044274 |
| 594 | P39880 | CUX1 | Homeobox protein cut-like 1 | qleekLKGQADYEEVKkelni | 345 | 355 | 132.6 | 0.000443401 |
| 595 | Q13948 | CUX1 | Protein CASP | qleekLKGQADYEEVKkelni | 356 | 366 | 132.6 | 0.000443401 |
| 596 | P01100 | FOS | Proto-oncogene c-Fos | eltdtLQAETDQLEDEksalq | 165 | 175 | 132.6 | 0.000443684 |
| 597 | Q8IY92 | SLX4 | Structure-specific endonuclease subunit SLX4 | eeaetLLKSKDHEEDQenvne | 833 | 843 | 132.6 | 0.000444722 |
| 598 | Q9HCE3 | ZNF532 | Zinc finger protein 532 | pdfddLLAAFDIPDMVdpkaa | 14 | 24 | 132.6 | 0.000445382 |
| 599 | Q5T848 | GPR158 | Probable G-protein coupled receptor 158 | pgttqMKDNFDIGEVCpwevy | 997 | 1007 | 132.6 | 0.000445665 |
| 601 | Q86X83 | COMMD2 | COMM domain-containing protein 2 | tiklhLNQNGDHNTKVlqtdp | 154 | 164 | 132.6 | 0.000445854 |
| 600 | Q92834 | RPGR | X-linked retinitis pigmentosa GTPase regulator | tqdtaLTENDDSDEYEemsem | 538 | 548 | 132.6 | 0.000445854 |
| 602 | Q6ZP82 | CCDC141 | Coiled-coil domain-containing protein 141 | kispvLSNAMDVGSTRsesek | 481 | 491 | 132.6 | 0.000447364 |
| 603 | Q8TDJ6 | DMXL2 | DmX-like protein 2 | nweqiLQEKMDQFEGPppnyi | 2557 | 2567 | 132.6 | 0.000448119 |
| 604 | Q9NZM1 | MYOF | Myoferlin | dpersLLTEADAGHTEftdev | 919 | 929 | 132.6 | 0.000449629 |
| 606 | Q5W111 | SPRYD7 | SPRY domain-containing protein 7 | rdmhsLVMRNDGALYHnneek | 100 | 110 | 132.6 | 0.000451044 |
| 605 | Q9Y496 | KIF3A | Kinesin-like protein KIF3A | eerkaLETKLDMEEEErnkar | 433 | 443 | 132.6 | 0.000451044 |
| 607 | Q5JRA6 | MIA3 | Transport and Golgi organization protein 1 homolog | ddkniLTTWGDTIFSlvtgge | 380 | 390 | 132.6 | 0.000451988 |
| 608 | Q8IYI6 | EXOC8 | Exocyst complex component 8 | lyvkqLSQQSDGDRDLqehrq | 31 | 41 | 132.6 | 0.000452742 |
| 609 | Q9BXW6 | OSBPL1A | Oxysterol-binding protein-related protein 1 | aevnvLNDMGDTPLHRaaftg | 77 | 87 | 132.6 | 0.000452742 |
| 610 | Q86TC9 | MYPN | Myopalladin | vnlarLAINYDPLEKAdetqa | 81 | 91 | 132.6 | 0.000452931 |
| 611 | Q9Y2Y8 | PRG3 | Proteoglycan 3 | phlesLETQADLGQDLdsske | 30 | 40 | 132.5 | 0.000453497 |

| | | | | | | | | |
|---|---|---|---|---|---|---|---|---|
| 612 | Q14919 | DRAP1 | Dr1-associated corepressor | kqcieLEQQFDFLKDLvasvp | 76 | 86 | 132.5 | 0.000453592 |
| 613 | Q6PJW8 | CNST | Consortin | glvsiLKKRNDTVGDHpaqmq | 623 | 633 | 132.5 | 0.000454441 |
| 614 | Q7KZ85 | SUPT6H | Transcription elongation factor SPT6 | kqaekMMETMDQGDVlirpss | 1346 | 1356 | 132.5 | 0.000456045 |
| 615 | Q86YR7 | MCF2L2 | Probable guanine nucleotide exchange factor MCF2L2 | lslagLFQSDDSHETCssksa | 1039 | 1049 | 132.5 | 0.000456989 |
| 616 | Q6ZMQ8 | AATK | Serine/threonine-protein kinase LMTK1 | gqpraLDSGYDTENYEspefv | 935 | 945 | 132.5 | 0.000457272 |
| 617 | Q9UJJ9 | GNPTG | N-acetylglucosamine-1-phosphotransferase subunit gamma | tlpeaLQRQWDQVEQDladel | 184 | 194 | 132.5 | 0.000458215 |
| 619 | P30511 | HLA-F | HLA class I histocompatibility antigen, alpha chain F | paeitLTWQRDGEEQTqdtel | 236 | 246 | 132.5 | 0.000458782 |
| 618 | Q53GG5 | PDLIM3 | PDZ and LIM domain protein 3 | gsfrvLQGMVDDGSDDrpagt | 256 | 266 | 132.5 | 0.000458782 |
| 620 | Q14004 | CDK13 | Cyclin-dependent kinase 13 | sgqddLIQHQDMRILEltpep | 1234 | 1244 | 132.5 | 0.000459725 |
| 621 | Q68DN1 | C2orf16 | Uncharacterized protein C2orf16 | rtaseLGGLWDSGIQEvsral | 750 | 760 | 132.5 | 0.000460386 |
| 622 | P41220 | RGS2 | Regulator of G-protein signaling 2 | qsamfLAVQHDCRPMDksags | 7 | 17 | 132.5 | 0.00046048 |
| 623 | O15056 | SYNJ2 | Synaptojanin-2 | eaaplLGDYQDPFWNLlhhpk | 1408 | 1418 | 132.5 | 0.000461895 |
| 624 | Q9NYQ6 | CELSR1 | Cadherin EGF LAG seven-pass G-type receptor 1 | htahvLINVTDANTHRpvfqs | 776 | 786 | 132.5 | 0.000463971 |
| 625 | O75145 | PPFIA3 | Liprin-alpha-3 | dlkeaLAQREDMEERIttlek | 295 | 305 | 132.5 | 0.00046416 |
| 626 | Q12873 | CHD3 | Chromodomain-helicase-DNA-binding protein 3 | aiarlLDRNQDATEDTdvqnm | 1264 | 1274 | 132.5 | 0.00046416 |
| 628 | P62191 | PSMC1 | 26S proteasome regulatory subunit 4 | msvgtLEEIIDDNHAIvstsv | 112 | 122 | 132.5 | 0.000466142 |
| 627 | Q96JN2 | CCDC136 | Coiled-coil domain-containing protein 136 | srlctLQKKYDTSQDEqnell | 374 | 384 | 132.5 | 0.000466142 |
| 629 | Q9H160 | ING2 | Inhibitor of growth protein 2 | kyqetLKEIDDVYEKYkkedd | 60 | 70 | 132.5 | 0.00046633 |
| 630 | Q9NRE2 | TSHZ2 | Teashirt homolog 2 | vsrryLFENSDQPIDLtksks | 757 | 767 | 132.5 | 0.00046633 |
| 631 | Q92538 | GBF1 | Golgi-specific brefeldin A-resistance guanine nucleotide exchange factor 1 | sgcsdLEEAVDSGADKkfark | 664 | 674 | 132.4 | 0.000467935 |
| 632 | P19087 | GNAT2 | Guanine nucleotide-binding protein G(t) subunit alpha-2 | ddgrqLNNLADSIEEGtmppe | 107 | 117 | 132.4 | 0.000468218 |
| 633 | Q8TBZ6 | TRMT10A | tRNA methyltransferase 10 homolog A | shggqLKKNMDENDKGwvnwk | 150 | 160 | 132.4 | 0.000469633 |
| 634 | Q7M4L6 | SHF | SH2 domain-containing adapter protein F | rplaiLEDYADPFDVQetgeg | 105 | 115 | 132.4 | 0.00047001 |
| 635 | Q9BRK0 | REEP2 | Receptor expression-enhancing protein 2 | spgslLDTIEDLGDDPalslr | 180 | 190 | 132.4 | 0.00047001 |
| 636 | Q9BUU2 | METTL22 | Methyltransferase-like protein 22 | lsqfkLLWSQDSWTDSgakgg | 52 | 62 | 132.4 | 0.00047001 |
| 637 | P0C1Z6 | TFPT | TCF3 fusion partner | qerrfLMRVLDSYGDDyrasq | 120 | 130 | 132.4 | 0.000471143 |
| 638 | Q9P2N5 | RBM27 | RNA-binding protein 27 | tetsdLFLPDDDDEDEdeyes | 1040 | 1050 | 132.4 | 0.00047152 |
| 639 | O95359 | TACC2 | Transforming acidic coiled-coil-containing protein 2 | pmkapLCGEGDQPGGFesqek | 213 | 223 | 132.4 | 0.000471898 |
| 640 | Q12955 | ANK3 | Ankyrin-3 | tkaaaLLLQNDNNADVesksg | 220 | 230 | 132.4 | 0.000472181 |
| 641 | Q9NS87 | KIF15 | Kinesin-like protein KIF15 | hknlkLQQHVDKLEHHstqmq | 743 | 753 | 132.4 | 0.000472653 |
| 642 | Q15029 | EFTUD2 | 116 kDa U5 small nuclear ribonucleoprotein component | retkdLDEMDDDDDDDdvgdh | 33 | 43 | 132.4 | 0.000474917 |
| 643 | Q9H3S7 | PTPN23 | Tyrosine-protein phosphatase non-receptor type 23 | dknevLDQFMDSMQLDpetvd | 394 | 404 | 132.4 | 0.00047605 |
| 644 | Q9H0K6 | PUS7L | Pseudouridylate synthase 7 homolog-like protein | qevhtLIKYTDGDQNHqsgse | 91 | 101 | 132.4 | 0.000476804 |
| 645 | Q9P2P6 | STARD9 | StAR-related lipid transfer protein 9 | givgsLCPSPDMQEFHsckge | 1357 | 1367 | 132.4 | 0.000476993 |
| 646 | Q86Y26 | NUTM1 | NUT family member 1 | afpssLLVTGDGGPCLsgaga | 70 | 80 | 132.4 | 0.000477087 |
| 647 | Q6V0I7 | FAT4 | Protocadherin Fat 4 | rdgntLEMHGDTCQPGifnya | 4731 | 4741 | 132.4 | 0.000477182 |
| 648 | Q8TE04 | PANK1 | Pantothenate kinase 1 | akkcrLRRRMDSGRKNrppfp | 222 | 232 | 132.4 | 0.000477937 |
| 649 | Q8IW19 | APLF | Aprataxin and PNK-like factor | nsvsfLGENRDCNKQQpilae | 166 | 176 | 132.3 | 0.000480768 |
| 650 | Q7Z7J9 | CAMK2N1 | Calcium/calmodulin-dependent protein kinase II inhibitor 1 | ygdekLSPYGDGGDVGqifsc | 12 | 22 | 132.3 | 0.000480862 |
| 651 | O94806 | PRKD3 | Serine/threonine-protein kinase D3 | gepssLGTDTDIPMDIdnndi | 334 | 344 | 132.3 | 0.000481617 |
| 652 | P48378 | RFX2 | DNA-binding protein RFX2 | slsltLLDKDDMGDEQrgsea | 682 | 692 | 132.3 | 0.000481617 |
| 653 | Q8IX12 | CCAR1 | Cell division cycle and apoptosis regulator protein 1 | ktirnLSTVMDEIHTVlkkdn | 1118 | 1128 | 132.3 | 0.000481806 |
| 654 | P14209 | CD99 | CD99 antigen | gllgvLVAAPDGGFDLsdalp | 18 | 28 | 132.3 | 0.000482277 |
| 655 | Q9UQ90 | SPG7 | Paraplegin | pfsqgLQQMMDHEARLlvaka | 695 | 705 | 132.3 | 0.000482277 |
| 656 | A6NKF2 | ARID3C | AT-rich interactive domain-containing protein 3C | ekravLMGPMDPPRPCmppsf | 325 | 335 | 132.3 | 0.000483315 |
| 657 | Q16720 | ATP2B3 | Plasma membrane calcium-transporting ATPase 3 | thnipLIDDTDVDENEerlra | 1158 | 1168 | 132.3 | 0.000484542 |
| 658 | Q9NUD7 | C20orf96 | Uncharacterized protein C20orf96 | tlmrqLQQVKDSQQDElddlg | 233 | 243 | 132.3 | 0.000484542 |
| 659 | P54257 | HAP1 | Huntingtin-associated protein 1 | eaeegLMLAADIMRGEdftpa | 542 | 552 | 132.3 | 0.00048558 |
| 660 | Q7L2J0 | MEPCE | 7SK snRNA methylphosphate capping enzyme | tdplsLNTCTDEGHVVlaspl | 241 | 251 | 132.3 | 0.00048558 |
| 661 | Q9BV73 | CEP250 | Centrosome-associated protein CEP250 | eavnlLQQQHDQWEEEgkalr | 419 | 429 | 132.3 | 0.000487562 |
| 662 | Q86TB3 | ALPK2 | Alpha-protein kinase 2 | dtaltLENVCDEPRDReavca | 788 | 798 | 132.3 | 0.000487656 |
| 663 | Q8IUN9 | CLEC10A | C-type lectin domain family 10 member A | wqghgLGGGEDCAHFHpdgrw | 276 | 286 | 132.3 | 0.000488694 |
| 664 | Q02446 | SP4 | Transcription factor Sp4 | tssdtLVSSADTGQYAstsas | 338 | 348 | 132.3 | 0.000489449 |
| 665 | P78362 | SRPK2 | SRSF protein kinase 2 | sleqqLDDEDDDEEDCpnpee | 398 | 408 | 132.3 | 0.000489637 |
| 666 | Q5VZ89 | DENND4C | DENN domain-containing protein 4C | geatnLKKNGDRGEKRqkhfp | 1026 | 1036 | 132.3 | 0.000489921 |
| 667 | Q9ULE0 | WWC3 | Protein WWC3 | srdspLAQLADSCEGPglgal | 481 | 491 | 132.3 | 0.000489921 |
| 668 | Q12955 | ANK3 | Ankyrin-3 | edlteLLNGMDEELDSpeelg | 1103 | 1113 | 132.3 | 0.000490581 |
| 669 | Q8WWM7 | ATXN2L | Ataxin-2-like protein | dlesdMSNGWDPNEMFkfnee | 247 | 257 | 132.3 | 0.000491053 |
| 670 | Q5VU43 | PDE4DIP | Myomegalin | flsdeLEACSDMDIVSeythy | 1617 | 1627 | 132.3 | 0.000491619 |
| 671 | Q8IYB7 | DIS3L2 | DIS3-like exonuclease 2 | lpqqsLKSYNDSPDVIveaqf | 155 | 165 | 132.3 | 0.000492846 |
| 672 | Q8N987 | NECAB1 | N-terminal EF-hand calcium-binding protein 1 | eelheLFHTIDTHNTNnldte | 68 | 78 | 132.2 | 0.000493034 |
| 673 | Q96IZ0 | PAWR | PRKC apoptosis WT1 regulator protein | eeidlLNRDLDDIEDEneqlk | 310 | 320 | 132.2 | 0.000493978 |
| 674 | Q2KJY2 | KIF26B | Kinesin-like protein KIF26B | ntfaeLQERLDCIDGSeepss | 929 | 939 | 132.2 | 0.00049511 |
| 675 | Q6KC79 | NIPBL | Nipped-B-like protein | ldfklLMEHLDPDEEEeegev | 2624 | 2634 | 132.2 | 0.000495582 |
| 676 | P53420 | COL4A4 | Collagen alpha-4(IV) chain | glkgeLGLVGDPGLFGligpk | 328 | 338 | 132.2 | 0.000495865 |
| 677 | Q9UPW6 | SATB2 | DNA-binding protein SATB2 | klkehLGSAVDVAEYKdeell | 680 | 690 | 132.2 | 0.00049662 |
| 678 | P55201 | BRPF1 | Peregrin | vkksfLVYRNDCSLPRsssds | 986 | 996 | 132.2 | 0.000496998 |
| 680 | P25686 | DNAJB2 | DnaJ homolog subfamily B member 2 | yrrkaLQWHPDKNPDNkefae | 28 | 38 | 132.2 | 0.000497092 |
| 679 | Q9Y2G0 | EFR3B | Protein EFR3 homolog B | vprpsLHQAVDTGRTGenrnr | 410 | 420 | 132.2 | 0.000497092 |
| 681 | Q5JRC9 | FAM47A | Protein FAM47A | dsgrtLKDVWDRCEARvkktk | 438 | 448 | 132.2 | 0.000498413 |
| 682 | P30622 | CLIP1 | CAP-Gly domain-containing linker protein 1 | cdcfdLHDTEDCPTQAqmsed | 1387 | 1397 | 132.2 | 0.000500678 |
| 683 | O75976 | CPD | Carboxypeptidase D | aglgsLIPEGDAGPDAagpda | 106 | 116 | 132.2 | 0.000500772 |
| 684 | Q13615 | MTMR3 | Myotubularin-related protein 3 | lgdaaLRSHLDMSWPLfsqgi | 750 | 760 | 132.2 | 0.000501338 |
| 685 | Q9H2D6 | TRIOBP | TRIO and F-actin-binding protein | astesLVPSMDSLHECphipt | 1147 | 1157 | 132.2 | 0.000501338 |
| 686 | Q8WWA1 | TMEM40 | Transmembrane protein 40 | vlkdeLQLYGDAPGEVvpsge | 114 | 124 | 132.2 | 0.000501433 |
| 687 | O15018 | PDZD2 | PDZ domain-containing protein 2 | wlqnsLQEGGDGPEQRlcqaa | 23 | 33 | 132.2 | 0.000501621 |
| 688 | P22413 | ENPP1 | Ectonucleotide pyrophosphatase/phosphodiesterase family member 1 | qaaasLLAPMDVGEEPlekaa | 49 | 59 | 132.2 | 0.000502187 |
| 689 | Q6V0I7 | FAT4 | Protocadherin Fat 4 | eveitLQDINDNPPVFptdml | 1730 | 1740 | 132.2 | 0.000502659 |
| 690 | Q9NQC8 | IFT46 | Intraflagellar transport protein 46 homolog | ehgapLEGAYDPADYEhlpvs | 60 | 70 | 132.2 | 0.00050332 |
| 691 | Q13813 | SPTAN1 | Spectrin alpha chain, non-erythrocytic 1 | hstvgLAQQWDQLDQLgmrmq | 2290 | 2300 | 132.2 | 0.000504263 |
| 692 | Q16584 | MAP3K11 | Mitogen-activated protein kinase kinase kinase 11 | rapwtLFPDSDPFWDSppanp | 805 | 815 | 132.2 | 0.000504924 |
| 693 | Q9BRD0 | BUD13 | BUD13 homolog | emqkpLARYIDDEDLDrmlre | 522 | 532 | 132.2 | 0.000505867 |
| 694 | Q9H2K2 | TNKS2 | Poly [ADP-ribose] polymerase tankyrase-2 | dgntpLDLVKDGDTDIqdllr | 629 | 639 | 132.2 | 0.000505962 |
| 695 | Q9Y6Z7 | COLEC10 | Collectin-10 | gikgeLGDMGDQGNIGktgpi | 79 | 89 | 132.2 | 0.000505962 |
| 696 | P08235 | NR3C2 | Mineralocorticoid receptor | tvaesMGLYMDSVRDAdysye | 105 | 115 | 132.2 | 0.000506717 |
| 697 | O60216 | RAD21 | Double-strand-break repair protein rad21 homolog | ilddkLISNNDGGIFDdppal | 233 | 243 | 132.1 | 0.00050851 |
| 698 | P48634 | PRRC2A | Protein PRRC2A | kefdqLDQENDDGWAGaheev | 318 | 328 | 132.1 | 0.000509925 |
| 699 | Q9UBB5 | MBD2 | Methyl-CpG-binding domain protein 2 | lmadiLSRAADTEEMDiemds | 392 | 402 | 132.1 | 0.000510302 |

| | | | | | | | | |
|---|---|---|---|---|---|---|---|---|
| 702 | P15848 | ARSB | Arylsulfatase B | sprieLLHNIDPNFVDsspcp | 391 | 401 | 132.1 | 0.000511435 |
| 700 | P35398 | RORA | Nuclear receptor ROR-alpha | elhddLSNYIDGHTPEgskad | 200 | 210 | 132.1 | 0.000511435 |
| 701 | Q9BYG4 | PARD6G | Partitioning defective 6 homolog gamma | vagktLDQVTDMMIANshnli | 229 | 239 | 132.1 | 0.000511435 |
| 703 | Q8TDI0 | CHD5 | Chromodomain-helicase-DNA-binding protein 5 | rraayLNMTQDPNHPAmalna | 1812 | 1822 | 132.1 | 0.000511623 |
| 704 | P40189 | IL6ST | Interleukin-6 receptor subunit beta | pedlqLVDHVDGGDGIlprqq | 798 | 808 | 132.1 | 0.000512567 |
| 705 | Q8N6H7 | ARFGAP2 | ADP-ribosylation factor GTPase-activating protein 2 | isssdLFGDMDGAHGAgsvsl | 465 | 475 | 132.1 | 0.00051285 |
| 706 | Q6NUM6 | TYW1B | S-adenosyl-L-methionine-dependent tRNA 4-demethylwyosine synthase TY | vdvedLGKIMDHVKKEkreke | 271 | 281 | 132.1 | 0.000512945 |
| 707 | Q9NV66 | TYW1 | S-adenosyl-L-methionine-dependent tRNA 4-demethylwyosine synthase TY | vdvedLGKIMDHVKKEkreke | 317 | 327 | 132.1 | 0.000512945 |
| 708 | Q9Y462 | ZNF711 | Zinc finger protein 711 | leshkLINKVDKTHEFteytr | 438 | 448 | 132.1 | 0.000513133 |
| 709 | Q9BUN5 | CCDC28B | Coiled-coil domain-containing protein 28B | lqhsfLTEVTDVYEMEeglln | 82 | 92 | 132.1 | 0.000513888 |
| 710 | O95817 | BAG3 | BAG family molecular chaperone regulator 3 | ekvqgLEQAVDNFEGKktdkk | 435 | 445 | 132.1 | 0.000514171 |
| 711 | O15063 | GARRE1 | Granule associated Rac and RHOG effector protein 1 | psppLTTVEDVNQDNktktw | 960 | 970 | 132.1 | 0.00051436 |
| 713 | Q8WXI7 | MUC16 | Mucin-16 | lltsgLAKTTDMLHKSsepvt | 8009 | 8019 | 132.1 | 0.000515398 |
| 712 | Q9ULL5 | PRR12 | Proline-rich protein 12 | hhpagLSGLFDTGLHHagsag | 68 | 78 | 132.1 | 0.000515398 |
| 714 | Q9Y2X9 | ZNF281 | Zinc finger protein 281 | ipsqeLASQIDPQKDIeprtt | 810 | 820 | 132.1 | 0.000517096 |
| 715 | Q8N5Y2 | MSL3 | Male-specific lethal 3 homolog | ndensLSSSSDCSENKdeeis | 129 | 139 | 132.1 | 0.000517285 |
| 716 | Q14524 | SCN5A | Sodium channel protein type 5 subunit alpha | ntaelLEQIPDLGQDVkdped | 1151 | 1161 | 132.1 | 0.000518134 |
| 717 | O95196 | CSPG5 | Chondroitin sulfate proteoglycan 5 | tpswsLLDLYDDFTPFdesdf | 250 | 260 | 132.1 | 0.000519361 |
| 718 | Q5VYK3 | ECPAS | Proteasome adapter and scaffold protein ECM29 | seqqdLERNADTLPDQeeliq | 777 | 787 | 132.1 | 0.000519927 |
| 720 | Q8TEQ8 | PIGO | GPI ethanolamine phosphate transferase 3 | endtlLVVAGDHGMTTngdhg | 272 | 282 | 132.1 | 0.000520776 |
| 719 | Q92481 | TFAP2B | Transcription factor AP-2-beta | ksvtsLMMNKDGFLGGmsvnt | 211 | 221 | 132.1 | 0.000520776 |
| 721 | O14578 | CIT | Citron Rho-interacting kinase | eeqakLQQMDLQKNHifrlt | 1159 | 1169 | 132.1 | 0.000522475 |
| 722 | Q96R06 | SPAG5 | Sperm-associated antigen 5 | lvpspLGQQQDMIFEArldtm | 137 | 147 | 132.1 | 0.000523041 |
| 723 | Q2M2I8 | AAK1 | AP2-associated protein kinase 1 | aedsnLISGFDVPEGSdkvae | 900 | 910 | 132.0 | 0.000523419 |
| 724 | Q9UJ78 | ZMYM5 | Zinc finger MYM-type protein 5 | amatsLMDIGDSFGHPacplv | 28 | 38 | 132.0 | 0.000523607 |
| 725 | Q9H7P9 | PLEKHG2 | Pleckstrin homology domain-containing family G member 2 | lqvpaLTTFSDQGHPEiqvpa | 969 | 979 | 132.0 | 0.000524457 |
| 726 | Q08AD1 | CAMSAP2 | Calmodulin-regulated spectrin-associated protein 2 | mrrkqLKLMEDMDTVlkprpq | 1242 | 1252 | 132.0 | 0.000524928 |
| 727 | Q8NHS0 | DNAJB8 | DnaJ homolog subfamily B member 8 | yrklaLRWHPDKNPDNkeeae | 28 | 38 | 132.0 | 0.000525117 |
| 728 | Q8NBF2 | NHLRC2 | NHL repeat-containing protein 2 | kncttLAGTGDTNNVTsssft | 512 | 522 | 132.0 | 0.000525211 |
| 729 | Q9H4L7 | SMARCAD1 | SWI/SNF-related matrix-associated actin-dependent regulator of chromatin s | aiaaaLLMFGDAGGGPrkrkl | 195 | 205 | 132.0 | 0.000525211 |
| 730 | P26599 | PTBP1 | Polypyrimidine tract-binding protein 1 | kkenaLVQMADGNQAQlamsh | 378 | 388 | 132.0 | 0.000525495 |
| 732 | O60524 | NEMF | Nuclear export mediator factor NEMF | pveveLMTQVDQEDITlqsgr | 720 | 730 | 132.0 | 0.00052691 |
| 731 | P11047 | LAMC1 | Laminin subunit gamma-1 | lidqkLKDYEDLREDMrgkel | 1304 | 1314 | 132.0 | 0.00052691 |
| 733 | O95249 | GOSR1 | Golgi SNAP receptor complex member 1 | almhtLQRHRDILQDYthefh | 108 | 118 | 132.0 | 0.000527193 |
| 734 | O95613 | PCNT | Pericentrin | kedsaLCGGGDICKSTscddt | 63 | 73 | 132.0 | 0.000528514 |
| 735 | Q9Y3S1 | WNK2 | Serine/threonine-protein kinase WNK2 | tfieqMKDVMDKAEDMlsedt | 1250 | 1260 | 132.0 | 0.000529175 |
| 736 | Q14457 | BECN1 | Beclin-1 | nlsrrLKVTGDLFDIMsgqtd | 116 | 126 | 132.0 | 0.000529552 |
| 738 | P20929 | NEB | Nebulin | pqlrqLKAAGDALSDKlyken | 375 | 385 | 132.0 | 0.000530024 |
| 737 | Q8NAN2 | MIGA1 | Mitoguardin 1 | sdpnsLADDIDKDTDItmkgn | 294 | 304 | 132.0 | 0.000530024 |
| 739 | Q9NS87 | KIF15 | Kinesin-like protein KIF15 | iveemLKMKADLEEVQsalyn | 1264 | 1274 | 132.0 | 0.00053059 |
| 740 | Q00653 | NFKB2 | Nuclear factor NF-kappa-B p100 subunit | dlaglLEALSDMGLEEgvrll | 832 | 842 | 132.0 | 0.000530873 |
| 741 | Q8TCN5 | ZNF507 | Zinc finger protein 507 | neeemLEVISDAEENLipdsl | 362 | 372 | 132.0 | 0.000530873 |
| 742 | Q6UWI2 | PARM1 | Prostate androgen-regulated mucin-like protein 1 | vsgkvMCELIDMETTTtfprv | 230 | 240 | 132.0 | 0.000531156 |
| 743 | Q9Y2L6 | FRMD4B | FERM domain-containing protein 4B | esqshLLSEMDSDKPFfslsk | 701 | 711 | 132.0 | 0.000531439 |
| 745 | P15336 | ATF2 | Cyclic AMP-dependent transcription factor ATF-2 | qlkqlLLAHKDCPVTAmqkks | 412 | 422 | 132.0 | 0.000531628 |
| 744 | P17544 | ATF7 | Cyclic AMP-dependent transcription factor ATF-7 | qlkqlLLAHKDCPVTAlqkkt | 392 | 402 | 132.0 | 0.000531628 |
| 746 | O43660 | PLRG1 | Pleiotropic regulator 1 | lvfrsLKRTHDMFVADngkpv | 18 | 28 | 132.0 | 0.000532005 |
| 747 | P07307 | ASGR2 | Asialoglycoprotein receptor 2 | whgheLGGSEDCVEVQpdgrw | 272 | 282 | 132.0 | 0.000532005 |
| 748 | Q7Z5N4 | SDK1 | Protein sidekick-1 | syklrLKATNDIGDSDfsset | 1553 | 1563 | 132.0 | 0.000533893 |
| 749 | P51587 | BRCA2 | Breast cancer type 2 susceptibility protein | fknstMVLYGDTGDKQatqvs | 927 | 937 | 132.0 | 0.000534836 |
| 750 | O60477 | BRINP1 | BMP/retinoic acid-inducible neural-specific protein 1 | ggeeaLTMYMDKSRLDrksgn | 141 | 151 | 132.0 | 0.000535308 |
| 751 | P51587 | BRCA2 | Breast cancer type 2 susceptibility protein | deeqhLESHTDCILAVkqais | 474 | 484 | 132.0 | 0.000537195 |
| 752 | Q9H1M0 | NUP62CL | Nucleoporin-62 C-terminal-like protein | rdqsgLHYLQDADEEHveist | 164 | 174 | 131.9 | 0.000539082 |
| 753 | Q5T9S5 | CCDC18 | Coiled-coil domain-containing protein 18 | qlsqsLITCNDSQESSklssl | 471 | 481 | 131.9 | 0.000539366 |
| 754 | Q12888 | TP53BP1 | TP53-binding protein 1 | eavtpLTKAADISLDNlvegk | 1611 | 1621 | 131.9 | 0.000541819 |
| 755 | Q9NQW6 | ANLN | Anillin | qasqaLNCCVDEEHGKgslee | 741 | 751 | 131.9 | 0.000542385 |
| 756 | A6NC57 | ANKRD62 | Ankyrin repeat domain-containing protein 62 | ryskeLNVLMDENTMLnselq | 625 | 635 | 131.9 | 0.000542574 |
| 757 | P0C264 | SBK3 | Uncharacterized serine/threonine-protein kinase SBK3 | eggssLEEWTDEGDDSksggr | 336 | 346 | 131.9 | 0.00054314 |
| 758 | Q6H8Q1 | ABLIM2 | Actin-binding LIM protein 2 | kksswLMLKGDADTRTnspdl | 464 | 474 | 131.9 | 0.000543234 |
| 759 | Q9H7P9 | PLEKHG2 | Pleckstrin homology domain-containing family G member 2 | srlpsLSDISDVFEMPclpai | 659 | 669 | 131.9 | 0.000543517 |
| 760 | P51693 | APLP1 | Amyloid-like protein 1 | ehegfLRAKMDLEERRmrqin | 311 | 321 | 131.9 | 0.00054465 |
| 761 | Q8N1G0 | ZNF687 | Zinc finger protein 687 | pdfddLLAAFDIPDIDaneai | 12 | 22 | 131.9 | 0.000544838 |
| 762 | Q03989 | ARID5A | AT-rich interactive domain-containing protein 5A | vkepqLVWGGDANRPSafhkg | 387 | 397 | 131.9 | 0.000545216 |
| 763 | Q92945 | KHSRP | Far upstream element-binding protein 2 | nvdkpLRIIGDPYKVQqacem | 283 | 293 | 131.9 | 0.000546159 |
| 764 | O43491 | EPB41L2 | Band 4.1-like protein 2 | qksytLVVAKDGGDKKeptqa | 90 | 100 | 131.9 | 0.000546726 |
| 765 | Q99538 | LGMN | Legumain | imkrkLMNTNDLEESRqltee | 319 | 329 | 131.9 | 0.000547481 |
| 766 | Q8TAU0 | NKX2-3 | Homeobox protein Nkx-2.3 | qlkksLETAGDCKAAEeserp | 128 | 138 | 131.9 | 0.000547669 |
| 767 | P07766 | CD3E | T-cell surface glycoprotein CD3 epsilon chain | pgseiLWQHNDKNIGGdeddk | 58 | 68 | 131.9 | 0.000548047 |
| 768 | P78312 | FAM193A | Protein FAM193A | kqhepLSFFFDIMQHHkegng | 1030 | 1040 | 131.9 | 0.000550028 |
| 769 | Q53GL0 | PLEKHO1 | Pleckstrin homology domain-containing family O member 1 | relrdLYRQMDLQTPDshlrq | 382 | 392 | 131.8 | 0.000552482 |
| 770 | Q96D09 | GPRASP2 | G-protein coupled receptor-associated sorting protein 2 | elakqLQAQIDNQNDPevgqq | 822 | 832 | 131.8 | 0.000553708 |
| 771 | Q9Y6R1 | SLC4A4 | Electrogenic sodium bicarbonate cotransporter 1 | rgasfLKHVCDEEEVEghhti | 14 | 24 | 131.8 | 0.000553991 |
| 774 | A2AJT9 | BCLAF3 | BCLAF1 and THRAP3 family member 3 | fqkgpLNRELDCFNTGrgret | 318 | 328 | 131.8 | 0.000554463 |
| 773 | O00311 | CDC7 | Cell division cycle 7-related protein kinase | measLGIQMDEPMAFspqrd | 5 | 15 | 131.8 | 0.000554463 |
| 772 | O75335 | PPFIA4 | Liprin-alpha-4 | dlreaLAQKEDMEERIttlek | 288 | 298 | 131.8 | 0.000554463 |
| 775 | Q8ND83 | SLAIN1 | SLAIN motif-containing protein 1 | kplssLSTLRDGNWRDgcy | 555 | 565 | 131.8 | 0.000554746 |
| 776 | Q6VY07 | PACS1 | Phosphofurin acidic cluster sorting protein 1 | eveedLDELYDSLEMYnpsds | 366 | 376 | 131.8 | 0.000554935 |
| 777 | O75096 | LRP4 | Low-density lipoprotein receptor-related protein 4 | iirnkLHFPMDIHTLHpqrqp | 681 | 691 | 131.8 | 0.000556539 |
| 778 | Q15813 | TBCE | Tubulin-specific chaperone E | vagpwLGVEWDNPERGkhdgs | 37 | 47 | 131.8 | 0.000556728 |
| 779 | Q9C0B7 | TANGO6 | Transport and Golgi organization protein 6 homolog | kilpdLLAQYDSSKDKhtpet | 917 | 927 | 131.8 | 0.000557388 |
| 780 | Q15011 | HERPUD1 | Homocysteine-responsive endoplasmic reticulum-resident ubiquitin-like dom | qrhrdLELSGDRGWSVghlka | 24 | 34 | 131.8 | 0.000558049 |
| 781 | Q562F6 | SGO2 | Shugoshin 2 | farvpLTSNDDEDEDKekmqc | 161 | 171 | 131.8 | 0.000558238 |
| 782 | Q99549 | MPHOSPH8 | M-phase phosphoprotein 8 | kdiqrLSLNNDIFEANsdsdq | 125 | 135 | 131.8 | 0.000558709 |
| 783 | A5D8V7 | ODAD3 | Outer dynein arm-docking complex subunit 3 | rialpLATSKDKFFDEeseee | 550 | 560 | 131.8 | 0.000559087 |
| 784 | P49821 | NDUFV1 | NADH dehydrogenase [ubiquinone] flavoprotein 1, mitochondrial | grpkyLVVNADEGEPGtckdr | 113 | 123 | 131.8 | 0.000559936 |
| 785 | Q16891 | IMMT | MICOS complex subunit MIC60 | ahsniLKAAMDNSEIAgekks | 256 | 266 | 131.8 | 0.000560691 |
| 786 | Q8NDM7 | CFAP43 | Cilia- and flagella-associated protein 43 | dafaqLMKAMDELDNIsnmpe | 1348 | 1358 | 131.8 | 0.000560974 |
| 787 | P50479 | PDLIM4 | PDZ and LIM domain protein 4 | dpargLPRSRDCRVDLgsevy | 176 | 186 | 131.8 | 0.000562012 |

| | | | | | | | | |
|---|---|---|---|---|---|---|---|---|
| 791 | O75616 | ERAL1 | GTPase Era, mitochondrial | eqdvlLVHHPDMPENSrvlrv | 101 | 111 | 131.8 | 0.000562578 |
| 789 | Q8WUN7 | UBTD2 | Ubiquitin domain-containing protein 2 | mtdgqLRSKRDEFWDTapafe | 50 | 60 | 131.8 | 0.000562578 |
| 790 | Q9HAC8 | UBTD1 | Ubiquitin domain-containing protein 1 | mtdgqLRSKRDEFWDTapafe | 47 | 57 | 131.8 | 0.000562578 |
| 788 | Q9ULJ8 | PPP1R9A | Neurabin-1 | ssdesLDMIDDEILDDgqspk | 961 | 971 | 131.8 | 0.000562578 |
| 792 | Q9HCR9 | PDE11A | Dual 3',5'-cyclic-AMP and -GMP phosphodiesterase 11A | ahsqpLPGGGDCGGVPlspsw | 87 | 97 | 131.8 | 0.000562767 |
| 793 | P61328 | FGF12 | Fibroblast growth factor 12 | qqgyfLQMHPDGTIDGtkden | 85 | 95 | 131.8 | 0.000562861 |
| 794 | O75379 | VAMP4 | Vesicle-associated membrane protein 4 | errnlLEDDSDEEEDFflrgp | 26 | 36 | 131.8 | 0.000563711 |
| 795 | P51798 | CLCN7 | H(+)/Cl(-) exchange transporter 7 | lddelLDPDMDPPHPFpkeip | 69 | 79 | 131.8 | 0.000563805 |
| 796 | Q9UIF9 | BAZ2A | Bromodomain adjacent to zinc finger domain protein 2A | pvsggLYGIDDTELMGaedkl | 314 | 324 | 131.8 | 0.000563994 |
| 797 | Q8N3U4 | STAG2 | Cohesin subunit SA-2 | griedLNEGMDFDTMDidlpp | 1185 | 1195 | 131.8 | 0.00056456 |
| 798 | Q9Y4F3 | MARF1 | Meiosis regulator and mRNA stability factor 1 | lknpkLCLIKDASEQSsaka | 613 | 623 | 131.8 | 0.000564843 |
| 799 | Q5VWQ0 | RSBN1 | Lysine-specific demethylase 9 | saraaLARCADGGAVGpfkcv | 29 | 39 | 131.8 | 0.000565032 |
| 800 | Q8NFZ8 | CADM4 | Cell adhesion molecule 4 | rpaatLRWYRDRKELKgvsss | 156 | 166 | 131.8 | 0.000565126 |
| 801 | Q8N7J2 | AMER2 | APC membrane recruitment protein 2 | vddtyLQEFWDMLSQTeeqgp | 436 | 446 | 131.8 | 0.000565692 |
| 802 | Q5FBB7 | SGO1 | Shugoshin 1 | ipqdtLGVDFDSGEAKstdnv | 169 | 179 | 131.7 | 0.000566824 |
| 803 | Q9ULJ1 | ODF2L | Protein BCAP | htireLQGQVDGNHNLltkls | 503 | 513 | 131.7 | 0.000567296 |
| 804 | Q96JM7 | L3MBTL3 | Lethal(3)malignant brain tumor-like protein 3 | nefgaLEVITDENEMEnvkka | 43 | 53 | 131.7 | 0.000567391 |
| 805 | Q92625 | ANKS1A | Ankyrin repeat and SAM domain-containing protein 1A | vvqvlLDAGMDSNYQTemgsa | 235 | 245 | 131.7 | 0.00056824 |
| 807 | O95210 | STBD1 | Starch-binding domain-containing protein 1 | apvlnLNQGMDNGRSTlvear | 228 | 238 | 131.7 | 0.000568334 |
| 806 | Q9BQN1 | FAM83C | Protein FAM83C | ssylaLPGGGDCSDTGvvsss | 365 | 375 | 131.7 | 0.000568334 |
| 808 | Q86XS8 | RNF130 | E3 ubiquitin-protein ligase RNF130 | grgapLTFRIDRGRYGldspk | 52 | 62 | 131.7 | 0.000569278 |
| 809 | Q5HYI7 | MTX3 | Metaxin-3 | nkesnLIEKMDDNLRQspqlp | 273 | 283 | 131.7 | 0.000571071 |
| 810 | P0CG31 | ZNF286B | Putative zinc finger protein 286B | thmnsLSEETDHEHDVywksf | 184 | 194 | 131.7 | 0.000571259 |
| 811 | Q9UJW9 | SERTAD3 | SERTA domain-containing protein 3 | silreLDTSMDGTEPPqnpvt | 102 | 112 | 131.7 | 0.000571731 |
| 812 | Q96SU4 | OSBPL9 | Oxysterol-binding protein-related protein 9 | tsdadLFDSHDDRDDDaeags | 352 | 362 | 131.7 | 0.00057192 |
| 813 | Q9UK76 | JPT1 | Jupiter microtubule associated homolog 1 | gdfldLKGEGDIHENVdtdlp | 98 | 108 | 131.7 | 0.000572769 |
| 815 | Q14191 | WRN | Werner syndrome ATP-dependent helicase | rehevLIHVEDETWDPtldhl | 339 | 349 | 131.7 | 0.000573147 |
| 814 | Q8IVL6 | P3H3 | Prolyl 3-hydroxylase 3 | anpmhLQMREDMAKYRrmsgv | 188 | 198 | 131.7 | 0.000573147 |
| 816 | Q9HBL6 | LRTM1 | Leucine-rich repeat and transmembrane domain-containing protein 1 | yhggpLAQTNDPGKVEekerf | 325 | 335 | 131.7 | 0.000573618 |
| 817 | O43747 | AP1G1 | AP-1 complex subunit gamma-1 | aggelLDLLGDINLTGapaaa | 657 | 667 | 131.7 | 0.000573807 |
| 818 | Q8NDA2 | HMCN2 | Hemicentin-2 | apsptLMWLKDGNPVSpagtp | 2017 | 2027 | 131.7 | 0.000574468 |
| 819 | P19532 | TFE3 | Transcription factor E3 | fpsdhLGDLGDPFHLGledil | 510 | 520 | 131.7 | 0.000574751 |
| 820 | A6QL64 | ANKRD36 | Ankyrin repeat domain-containing protein 36A | pkqpaLKAICDKEDSVpnmat | 1131 | 1141 | 131.7 | 0.000574845 |
| 822 | A8MW99 | MEI4 | Meiosis-specific protein MEI4 | ssestLTSMEDSGCDLsneqr | 96 | 106 | 131.7 | 0.000574845 |
| 821 | Q5T1M5 | FKBP15 | FK506-binding protein 15 | ppptpLFGDDDDDDDIdwlg | 1205 | 1215 | 131.7 | 0.000574845 |
| 823 | Q14137 | BOP1 | Ribosome biogenesis protein BOP1 | rtrdeLDQFLDKMDDPdywrt | 169 | 179 | 131.7 | 0.000575034 |
| 824 | Q9BZV3 | IMPG2 | Interphotoreceptor matrix proteoglycan 2 | dkklvLVDKMDSTDQIskhsk | 651 | 661 | 131.7 | 0.000575223 |
| 826 | Q92994 | BRF1 | Transcription factor IIIB 90 kDa subunit | vsalqMMGSNDYGCDGdeddg | 661 | 671 | 131.7 | 0.0005756 |
| 825 | Q9Y566 | SHANK1 | SH3 and multiple ankyrin repeat domains protein 1 | rrrstLFLSTDAGDEDggdgg | 1256 | 1266 | 131.7 | 0.0005756 |
| 827 | Q9H0E2 | TOLLIP | Toll-interacting protein | cseedLKAIQDMFPNMdqevi | 234 | 244 | 131.7 | 0.000576072 |
| 828 | Q9UPN7 | PPP6R1 | Serine/threonine-protein phosphatase 6 regulatory subunit 1 | vnthhLHSSSDDEDDRlkefn | 527 | 537 | 131.7 | 0.000576072 |
| 830 | Q9BT81 | SOX7 | Transcription factor SOX-7 | aerlrLQHMQDYPNYKyrprr | 103 | 113 | 131.7 | 0.000576261 |
| 829 | Q9NRC6 | SPTBN5 | Spectrin beta chain, non-erythrocytic 5 | eekrpLVSSRDYGRDEaatlr | 1648 | 1658 | 131.7 | 0.000576261 |
| 831 | Q12802 | AKAP13 | A-kinase anchor protein 13 | dqdscLQSLPDCGVKGtegls | 401 | 411 | 131.7 | 0.000576638 |
| 832 | Q9NZT1 | CALML5 | Calmodulin-like protein 5 | aqlrkLISEVDSDGDGeisfq | 52 | 62 | 131.7 | 0.000576921 |
| 833 | Q5TF21 | SOGA3 | Protein SOGA3 | neideLRTEMDEMRDTffeed | 373 | 383 | 131.7 | 0.000577299 |
| 834 | Q8NEP3 | DNAAF1 | Dynein assembly factor 1, axonemal | rdaapLTSSGDRDSDFlaass | 672 | 682 | 131.7 | 0.000578148 |
| 835 | Q6U949 | IGF2-AS | Putative insulin-like growth factor 2 antisense gene protein | ersnaLWQAVDAAEALalssp | 93 | 103 | 131.7 | 0.000578431 |
| 836 | Q8N3X6 | LCORL | Ligand-dependent nuclear receptor corepressor-like protein | lpdtgLYNMTDSGTGScknss | 584 | 594 | 131.7 | 0.00057928 |
| 837 | Q9UPW8 | UNC13A | Protein unc-13 homolog A | kkleqLNAMRDQDEYSfqdeq | 148 | 158 | 131.7 | 0.000579374 |
| 838 | Q15276 | RABEP1 | Rab GTPase-binding effector protein 1 | nagnkLGRRCDMCSNYekqlq | 529 | 539 | 131.7 | 0.000579469 |
| 839 | Q7Z2Z1 | TICRR | Treslin | vvdsiLNQTHDSLADTasaas | 457 | 467 | 131.7 | 0.000579846 |
| 840 | O95180 | CACNA1H | Voltage-dependent T-type calcium channel subunit alpha-1H | vhtdsLEGKIDSPRDTldpae | 2024 | 2034 | 131.6 | 0.000581356 |
| 841 | Q9BY07 | SLC4A5 | Electrogenic sodium bicarbonate cotransporter 4 | apnptLFTEMDTLQHDgdqme | 113 | 123 | 131.6 | 0.000583715 |
| 842 | Q8NDD1 | C1orf131 | Uncharacterized protein C1orf131 | lldalLQNLYDFGGTEgeteq | 33 | 43 | 131.6 | 0.000584847 |
| 843 | O43379 | WDR62 | WD repeat-containing protein 62 | qgdsyLRVSSDSPKDQspped | 976 | 986 | 131.6 | 0.000584942 |
| 844 | Q9HBH5 | RDH14 | Retinol dehydrogenase 14 | prvqrLRRGGDPGLMHgktvl | 32 | 42 | 131.6 | 0.000585414 |
| 845 | Q6UUV9 | CRTC1 | CREB-regulated transcription coactivator 1 | ltldgLHMLNDPDMVLadpat | 610 | 620 | 131.6 | 0.000585602 |
| 847 | B5ME19 | EIF3CL | Eukaryotic translation initiation factor 3 subunit C-like protein | esrkfLKKMDDEDEDSedsed | 212 | 222 | 131.6 | 0.000586263 |
| 846 | Q99613 | EIF3C | Eukaryotic translation initiation factor 3 subunit C | esrkfLKKMDDEDEDSedsed | 212 | 222 | 131.6 | 0.000586263 |
| 848 | P28290 | ITPRID2 | Protein ITPRID2 | ktlskLNLCVDKTEKGesssp | 302 | 312 | 131.6 | 0.000587395 |
| 849 | Q8TDM0 | BCAS4 | Breast carcinoma-amplified sequence 4 | dpvalLMLLVDADQPEpmrsg | 30 | 40 | 131.6 | 0.000587395 |
| 850 | Q5T7V8 | GORAB | RAB6-interacting golgin | eelrrLKQTKDPFEPQrrlpa | 41 | 51 | 131.6 | 0.000587961 |
| 851 | P40426 | PBX3 | Pre-B-cell leukemia transcription factor 3 | sgsfnLPNSGDMFMNMqslng | 344 | 354 | 131.6 | 0.000588999 |
| 853 | A6NIR3 | AGAP5 | Arf-GAP with GTPase, ANK repeat and PH domain-containing protein 5 | sksngLSKDMDTGLGDsicfs | 363 | 373 | 131.6 | 0.000589848 |
| 854 | Q5VW22 | AGAP6 | Arf-GAP with GTPase, ANK repeat and PH domain-containing protein 6 | sksngLSKDMDTGLGDsicfs | 340 | 350 | 131.6 | 0.000589848 |
| 852 | Q96P64 | AGAP4 | Arf-GAP with GTPase, ANK repeat and PH domain-containing protein 4 | sksngLSKDMDTGLGDsicfs | 340 | 350 | 131.6 | 0.000589848 |
| 855 | Q9NU22 | MDN1 | Midasin | klderLWGDDDEEEDEeeedn | 4778 | 4788 | 131.6 | 0.000590037 |
| 856 | Q5T4S7 | UBR4 | E3 ubiquitin-protein ligase UBR4 | efiesLDSTTDEEEDEeevyk | 4459 | 4469 | 131.6 | 0.000590698 |
| 857 | Q8IYT3 | CCDC170 | Coiled-coil domain-containing protein 170 | efltqLRDCLDPDERNdkasd | 175 | 185 | 131.6 | 0.000590698 |
| 858 | Q6ZN84 | CCDC81 | Coiled-coil domain-containing protein 81 | krkaiLHQLVDQRRDLqmlqr | 545 | 555 | 131.6 | 0.000590886 |
| 859 | Q9P2M4 | TBC1D14 | TBC1 domain family member 14 | kqsarLDKHNDLGWKLfgkap | 252 | 262 | 131.6 | 0.000590886 |
| 860 | P43243 | MATR3 | Matrin-3 | gdetdLANLGDVASDGkkeps | 681 | 691 | 131.6 | 0.000591075 |
| 861 | O43719 | HTATSF1 | HIV Tat-specific factor 1 | dadekLFEESDDKEDEdadgk | 672 | 682 | 131.6 | 0.000591264 |
| 863 | Q53GL7 | PARP10 | Protein mono-ADP-ribosyltransferase PARP10 | gselsLVPHYDILEPEelaen | 243 | 253 | 131.6 | 0.000592207 |
| 862 | Q9H582 | ZNF644 | Zinc finger protein 644 | gsnnfLHEIHDPQHLEtadas | 930 | 940 | 131.6 | 0.000592207 |
| 864 | P49427 | CDC34 | Ubiquitin-conjugating enzyme E2 R1 | irkqvLGTKVDAERDGvkvpt | 170 | 180 | 131.6 | 0.000593057 |
| 865 | Q9H582 | ZNF644 | Zinc finger protein 644 | edgedLLVKDDCVNTVtgiss | 231 | 241 | 131.6 | 0.000593434 |
| 866 | Q96SB8 | SMC6 | Structural maintenance of chromosomes protein 6 | fkinqLSELADPLKDElnlad | 790 | 800 | 131.5 | 0.000598058 |
| 867 | Q8WU20 | FRS2 | Fibroblast growth factor receptor substrate 2 | trrteLYAVIDIERTAamsnl | 470 | 480 | 131.5 | 0.000598435 |
| 868 | Q9NWQ8 | PAG1 | Phosphoprotein associated with glycosphingolipid-enriched microdomains 1 | nvesiLGNSCDPEEEApppvp | 246 | 256 | 131.5 | 0.000598435 |
| 869 | Q5H9L4 | TAF7L | Transcription initiation factor TFIID subunit 7-like | disqmLVCTADGDIHLspeep | 181 | 191 | 131.5 | 0.000598907 |
| 870 | P15407 | FOSL1 | Fos-related antigen 1 | eltdfLQAETDKLEDEksglq | 133 | 143 | 131.5 | 0.000599568 |
| 871 | Q9NPB6 | PARD6A | Partitioning defective 6 homolog alpha | vagktLDQVTDMMVANshnli | 228 | 238 | 131.5 | 0.000601455 |
| 872 | O15090 | ZNF536 | Zinc finger protein 536 | qnlgiMSQMSDIEDDArknrk | 114 | 124 | 131.5 | 0.000601549 |
| 873 | O94875 | SORBS2 | Sorbin and SH3 domain-containing protein 2 | scddlLNDDCDSFPDPkvkse | 388 | 398 | 131.5 | 0.000602115 |
| 874 | Q969S2 | NEIL2 | Endonuclease 8-like 2 | vhgkkLFLRFDLDEEMgppgs | 52 | 62 | 131.5 | 0.00060221 |
| 875 | Q9Y485 | DMXL1 | DmX-like protein 1 | ddsleLKWDSDNDEENedvpi | 1966 | 1976 | 131.5 | 0.000602776 |

| | | | | | | | | |
|---|---|---|---|---|---|---|---|---|
| 877 | Q96DR7 | ARHGEF26 | Rho guanine nucleotide exchange factor 26 | lrdqlLVESCDNEELNsspgk | 713 | 723 | 131.5 | 0.000604191 |
| 876 | Q9ULC0 | EMCN | Endomucin | vglyrMCWKADPGTPEngndq | 215 | 225 | 131.5 | 0.000604191 |
| 878 | Q13439 | GOLGA4 | Golgin subfamily A member 4 | telesLKHQQDALWTEklqvl | 619 | 629 | 131.5 | 0.000606267 |
| 879 | Q6ZMN7 | PDZRN4 | PDZ domain-containing RING finger protein 4 | eyissLPADADRTEDFeyeev | 388 | 398 | 131.5 | 0.000606833 |
| 880 | O94913 | PCF11 | Pre-mRNA cleavage complex 2 protein Pcf11 | rlkkhLQDKTDGKDDDvkekr | 408 | 418 | 131.5 | 0.000607494 |
| 882 | Q15075 | EEA1 | Early endosome antigen 1 | dleqvLRQIGDKDQKIqnlea | 515 | 525 | 131.5 | 0.00060806 |
| 881 | Q8TE73 | DNAH5 | Dynein heavy chain 5, axonemal | cihtlMRAMTDCGKPHremrm | 2276 | 2286 | 131.5 | 0.00060806 |
| 883 | Q92932 | PTPRN2 | Receptor-type tyrosine-protein phosphatase N2 | elmagLMQGVDHGVARgspgr | 339 | 349 | 131.5 | 0.000608249 |
| 884 | P09958 | FURIN | Furin | knhpdLAGNYDPGASFdvndq | 163 | 173 | 131.5 | 0.000608909 |
| 885 | Q9H4B4 | PLK3 | Serine/threonine-protein kinase PLK3 | sprgtLASSGDGFEEGltvat | 414 | 424 | 131.5 | 0.000609098 |
| 886 | P52655 | GTF2A1 | Transcription initiation factor IIA subunit 1 | qtqapLVLQVDGTGDTsseed | 269 | 279 | 131.5 | 0.000609759 |
| 889 | Q9NQL2 | RRAGD | Ras-related GTP-binding protein D | delvgLADYGDGPDSSdadpd | 28 | 38 | 131.5 | 0.000610419 |
| 888 | Q9UEW8 | STK39 | STE20/SPS1-related proline-alanine-rich protein kinase | gssghLHKTEDGDWEWsddem | 374 | 384 | 131.5 | 0.000610419 |
| 887 | Q9UPT8 | ZC3H4 | Zinc finger CCCH domain-containing protein 4 | mepglLGDAEDYGHYEelpge | 712 | 722 | 131.5 | 0.000610419 |
| 890 | Q9BW19 | KIFC1 | Kinesin-like protein KIFC1 | lsgsrLKRRPDQMEDGlepek | 35 | 45 | 131.5 | 0.000612023 |
| 892 | A6NJL1 | ZSCAN5B | Zinc finger and SCAN domain-containing protein 5B | gedflLHKSIDVTGDPnsprp | 202 | 212 | 131.5 | 0.000612212 |
| 893 | Q9BRR9 | ARHGAP9 | Rho GTPase-activating protein 9 | gpacpLLQRLDAWEQHldpns | 212 | 222 | 131.5 | 0.000612212 |
| 891 | Q9BUG6 | ZSCAN5A | Zinc finger and SCAN domain-containing protein 5A | gedflLHKSIDVTGDPkslrp | 202 | 212 | 131.5 | 0.000612212 |
| 894 | Q2NL68 | PROSER3 | Proline and serine-rich protein 3 | eadarLSFLLDQAEDLgswsp | 440 | 450 | 131.4 | 0.000613061 |
| 896 | Q12873 | CHD3 | Chromodomain-helicase-DNA-binding protein 3 | ekprfMFNIADGGFTElhtlw | 1708 | 1718 | 131.4 | 0.000613344 |
| 895 | Q9BYV9 | BACH2 | Transcription regulator protein BACH2 | eesitLCLSGDEPDAKdragd | 291 | 301 | 131.4 | 0.000613344 |
| 897 | A6NGG8 | PCARE | Photoreceptor cilium actin regulator | sptesLRMLGDSKDAGaspcl | 744 | 754 | 131.4 | 0.000613627 |
| 898 | Q5VST9 | OBSCN | Obscurin | ckaelLVLGGDNEPDSekqsh | 6445 | 6455 | 131.4 | 0.000613627 |
| 899 | Q99715 | COL12A1 | Collagen alpha-1(XII) chain | pststLNVRWDHAEGNprqyk | 1860 | 1870 | 131.4 | 0.00061542 |
| 900 | P23508 | MCC | Colorectal mutant cancer protein | nitqmLKRAHDCRKTAenaak | 499 | 509 | 131.4 | 0.000615703 |
| 901 | Q8N3U1 | | Putative uncharacterized protein LOC400692 | fglarLLGSQDHGDDPaergr | 32 | 42 | 131.4 | 0.000615798 |
| 902 | O60841 | EIF5B | Eukaryotic translation initiation factor 5B | kpkveMYSGSDDDDDFnklpk | 133 | 143 | 131.4 | 0.000615986 |
| 903 | Q9UQL6 | HDAC5 | Histone deacetylase 5 | syklpLPGPYDSRDDFplrkt | 242 | 252 | 131.4 | 0.00061759 |
| 905 | Q53TS8 | C2CD6 | C2 calcium-dependent domain-containing protein 6 | mvssgLVHINDKTSDYemhkm | 599 | 609 | 131.4 | 0.000617685 |
| 904 | Q9BXX0 | EMILIN2 | EMILIN-2 | dslhlLKSLNDTMHRKfqete | 583 | 593 | 131.4 | 0.000617685 |
| 907 | Q6ZSR9 | | Uncharacterized protein FLJ45252 | aqtglLSQTGDVPLPAgrera | 196 | 206 | 131.4 | 0.000620893 |
| 906 | Q9H5I5 | PIEZO2 | Piezo-type mechanosensitive ion channel component 2 | lkchgLWDEDDMTESGmaree | 2038 | 2048 | 131.4 | 0.000620893 |
| 908 | Q9HAP2 | MLXIP | MLX-interacting protein | hfvtpLDGSVDVDEHRrpeai | 189 | 199 | 131.4 | 0.000621365 |
| 909 | P01880 | IGHD | Immunoglobulin heavy constant delta | ppnilLMWLEDQREVNtsgfa | 306 | 316 | 131.4 | 0.000621837 |
| 910 | P0DOX3 | | Immunoglobulin delta heavy chain | ppnilLMWLEDQREVNtsgfa | 435 | 445 | 131.4 | 0.000621837 |
| 911 | P13805 | TNNT1 | Troponin T, slow skeletal muscle | kdlleLQTLIDVHFEQrkkee | 78 | 88 | 131.4 | 0.000621837 |
| 912 | Q8N443 | RIBC1 | RIB43A-like with coiled-coils protein 1 | alsnqLRLAMDAQATHlarle | 188 | 198 | 131.4 | 0.000622025 |
| 913 | Q9UKV0 | HDAC9 | Histone deacetylase 9 | sykytLPGAQDAKDDFplrkt | 203 | 213 | 131.4 | 0.00062212 |
| 914 | P23443 | RPS6KB1 | Ribosomal protein S6 kinase beta-1 | eeggqLNESMDHGGVGpyelg | 50 | 60 | 131.4 | 0.000624573 |
| 915 | Q70EL1 | USP54 | Inactive ubiquitin carboxyl-terminal hydrolase 54 | sqeaqLESGMDTEFGAssffh | 909 | 919 | 131.4 | 0.000624573 |
| 916 | P04626 | ERBB2 | Receptor tyrosine-protein kinase erbB-2 | tfyrsLLEDDDMGDLVdaeey | 1008 | 1018 | 131.3 | 0.000629102 |
| 917 | Q8IU99 | CALHM1 | Calcium homeostasis modulator protein 1 | eerekLRGITDQGTMNrllts | 290 | 300 | 131.3 | 0.000629197 |
| 918 | Q5VZ18 | SHE | SH2 domain-containing adapter protein E | etviiLEDYADPYDAKrtkgq | 201 | 211 | 131.3 | 0.00062948 |
| 919 | Q9UBU7 | DBF4 | Protein DBF4 homolog A | qtqvkLRIQTDGDKYGgtsiq | 269 | 279 | 131.3 | 0.000630046 |
| 920 | O43852 | CALU | Calumenin | aearhLVYESDQNKDGkltke | 273 | 283 | 131.3 | 0.000630707 |
| 921 | Q9P2Y5 | UVRAG | UV radiation resistance-associated gene protein | fmehgLMVRCDRHHTSsaipv | 462 | 472 | 131.3 | 0.000631178 |
| 922 | O60284 | ST18 | Suppression of tumorigenicity 18 protein | lnesnLKIEADMMKLQtqits | 941 | 951 | 131.3 | 0.000631462 |
| 924 | P51531 | SMARCA2 | Probable global transcription activator SNF2L2 | ylnsiLQHAKDFKEYHrsvag | 456 | 466 | 131.3 | 0.000631933 |
| 923 | P51532 | SMARCA4 | Transcription activator BRG1 | ylnsiLQHAKDFKEYHrsvtg | 480 | 490 | 131.3 | 0.000631933 |
| 935 | A0A087WUL8 | NBPF19 | Neuroblastoma breakpoint family member 19 | qqcvgLAVDMDEIEKYqevee | 643 | 653 | 131.3 | 0.000632216 |
| 933 | A0A087WUL8 | NBPF19 | Neuroblastoma breakpoint family member 19 | qqcvgLAVDMDEIEKYqevee | 1131 | 1141 | 131.3 | 0.000632216 |
| 958 | A0A087WUL8 | NBPF19 | Neuroblastoma breakpoint family member 19 | qqcvgLAVDMDEIEKYqevee | 1375 | 1385 | 131.3 | 0.000632216 |
| 963 | A0A087WUL8 | NBPF19 | Neuroblastoma breakpoint family member 19 | qqcvgLAVDMDEIEKYqevee | 1619 | 1629 | 131.3 | 0.000632216 |
| 931 | A0A087WUL8 | NBPF19 | Neuroblastoma breakpoint family member 19 | qqcvgLAVDMDEIEKYqevee | 1863 | 1873 | 131.3 | 0.000632216 |
| 943 | A0A087WUL8 | NBPF19 | Neuroblastoma breakpoint family member 19 | qqcvgLAVDMDEIEKYqevee | 2107 | 2117 | 131.3 | 0.000632216 |
| 940 | A0A087WUL8 | NBPF19 | Neuroblastoma breakpoint family member 19 | qqcvgLAVDMDEIEKYqevee | 2351 | 2361 | 131.3 | 0.000632216 |
| 941 | A0A087WUL8 | NBPF19 | Neuroblastoma breakpoint family member 19 | qqcvgLAVDMDEIEKYqevee | 2595 | 2605 | 131.3 | 0.000632216 |
| 939 | A0A087WUL8 | NBPF19 | Neuroblastoma breakpoint family member 19 | qqcvgLAVDMDEIEKYqevee | 2839 | 2849 | 131.3 | 0.000632216 |
| 928 | A0A087WUL8 | NBPF19 | Neuroblastoma breakpoint family member 19 | qqcvgLAVDMDEIEKYqevee | 3083 | 3093 | 131.3 | 0.000632216 |
| 965 | A0A087WUL8 | NBPF19 | Neuroblastoma breakpoint family member 19 | qqcvgLAVDMDEIEKYqevee | 3327 | 3337 | 131.3 | 0.000632216 |
| 955 | A0A087WUL8 | NBPF19 | Neuroblastoma breakpoint family member 19 | qqcvgLAVDMDEIEKYqevee | 3571 | 3581 | 131.3 | 0.000632216 |
| 949 | B4DH59 | NBPF26 | Neuroblastoma breakpoint family member 26 | qqrvgLAVDMDEIEKYqevee | 330 | 340 | 131.3 | 0.000632216 |
| 954 | B4DH59 | NBPF26 | Neuroblastoma breakpoint family member 26 | qqrvgLAVDMDEIEKYqevee | 405 | 415 | 131.3 | 0.000632216 |
| 929 | B4DH59 | NBPF26 | Neuroblastoma breakpoint family member 26 | qqrvgLAVDMDEIEKYqevee | 480 | 490 | 131.3 | 0.000632216 |
| 961 | B4DH59 | NBPF26 | Neuroblastoma breakpoint family member 26 | qqrvgLAVDMDEIEKYqevee | 555 | 565 | 131.3 | 0.000632216 |
| 968 | B4DH59 | NBPF26 | Neuroblastoma breakpoint family member 26 | qqrvgLAVDMDEIEKYqevee | 630 | 640 | 131.3 | 0.000632216 |
| 953 | P0DPF2 | NBPF20 | Neuroblastoma breakpoint family member 20 | qqrigLAVDMDEIEKYqevee | 55 | 65 | 131.3 | 0.000632216 |
| 957 | P0DPF2 | NBPF20 | Neuroblastoma breakpoint family member 20 | qqrvgLAVDMDEIEKYqevee | 299 | 309 | 131.3 | 0.000632216 |
| 964 | P0DPF2 | NBPF20 | Neuroblastoma breakpoint family member 20 | qqrvgLAVDMDEIEKYqevee | 543 | 553 | 131.3 | 0.000632216 |
| 934 | P0DPF2 | NBPF20 | Neuroblastoma breakpoint family member 20 | qqrvgLAVDMDEIEKYqevee | 787 | 797 | 131.3 | 0.000632216 |
| 962 | P0DPF2 | NBPF20 | Neuroblastoma breakpoint family member 20 | qqrvgLAVDMDEIEKYqevee | 1031 | 1041 | 131.3 | 0.000632216 |
| 946 | P0DPF2 | NBPF20 | Neuroblastoma breakpoint family member 20 | qqrvgLAVDMDEIEKYqevee | 1275 | 1285 | 131.3 | 0.000632216 |
| 947 | P0DPF2 | NBPF20 | Neuroblastoma breakpoint family member 20 | qqrvgLAVDMDEIEKYqevee | 1519 | 1529 | 131.3 | 0.000632216 |
| 937 | P0DPF2 | NBPF20 | Neuroblastoma breakpoint family member 20 | qqrvgLAVDMDEIEKYqevee | 1763 | 1773 | 131.3 | 0.000632216 |
| 944 | P0DPF2 | NBPF20 | Neuroblastoma breakpoint family member 20 | qqrvgLAVDMDEIEKYqevee | 2007 | 2017 | 131.3 | 0.000632216 |
| 952 | P0DPF2 | NBPF20 | Neuroblastoma breakpoint family member 20 | qqrvgLAVDMDEIEKYqevee | 2251 | 2261 | 131.3 | 0.000632216 |
| 932 | P0DPF2 | NBPF20 | Neuroblastoma breakpoint family member 20 | qqrvgLAVDMDEIEKYqevee | 2495 | 2505 | 131.3 | 0.000632216 |
| 951 | P0DPF2 | NBPF20 | Neuroblastoma breakpoint family member 20 | qqrvgLAVDMDEIEKYqevee | 2739 | 2749 | 131.3 | 0.000632216 |
| 936 | P0DPF2 | NBPF20 | Neuroblastoma breakpoint family member 20 | qqrvgLAVDMDEIEKYqevee | 2983 | 2993 | 131.3 | 0.000632216 |
| 938 | P0DPF2 | NBPF20 | Neuroblastoma breakpoint family member 20 | qqrvgLAVDMDEIEKYqevee | 3227 | 3237 | 131.3 | 0.000632216 |
| 948 | P0DPF2 | NBPF20 | Neuroblastoma breakpoint family member 20 | qqrvgLAVDMDEIEKYqevee | 3471 | 3481 | 131.3 | 0.000632216 |
| 959 | P0DPF2 | NBPF20 | Neuroblastoma breakpoint family member 20 | qqrvgLAVDMDEIEKYqevee | 3715 | 3725 | 131.3 | 0.000632216 |
| 966 | P0DPF2 | NBPF20 | Neuroblastoma breakpoint family member 20 | qqrvgLAVDMDEIEKYqevee | 3959 | 3969 | 131.3 | 0.000632216 |
| 956 | P0DPF2 | NBPF20 | Neuroblastoma breakpoint family member 20 | qqrvgLAVDMDEIEKYqevee | 4203 | 4213 | 131.3 | 0.000632216 |
| 967 | P0DPF2 | NBPF20 | Neuroblastoma breakpoint family member 20 | qqrvgLAVDMDEIEKYqevee | 4691 | 4701 | 131.3 | 0.000632216 |
| 945 | P0DPF2 | NBPF20 | Neuroblastoma breakpoint family member 20 | qqrvgLAVDMDEIEKYqevee | 4935 | 4945 | 131.3 | 0.000632216 |
| 930 | P0DPF3 | NBPF9 | Neuroblastoma breakpoint family member 9 | qqrvgLAVDMDEIEKYqevee | 670 | 680 | 131.3 | 0.000632216 |
| 925 | Q3BBV2 | NBPF8 | Putative neuroblastoma breakpoint family member 8 | qqrvgLAVDMDEIEKYqevee | 635 | 645 | 131.3 | 0.000632216 |

| | | | | | | | | |
|---|---|---|---|---|---|---|---|---|
| 926 | Q5TAG4 | NBPF12 | Neuroblastoma breakpoint family member 12 | qqrvgLAVDMDEIEKYqevee | 941 | 951 | 131.3 | 0.000632216 |
| 969 | Q5TAG4 | NBPF12 | Neuroblastoma breakpoint family member 12 | qqrvgLAVDMDEIEKYqevee | 1185 | 1195 | 131.3 | 0.000632216 |
| 950 | Q5TI25 | NBPF14 | Neuroblastoma breakpoint family member 14 | qqrvgLAVDMDEIEKYqevee | 574 | 584 | 131.3 | 0.000632216 |
| 942 | Q5TI25 | NBPF14 | Neuroblastoma breakpoint family member 14 | qqrvgLAVDMDEIEKYqevee | 649 | 659 | 131.3 | 0.000632216 |
| 960 | Q6P3W6 | NBPF10 | Neuroblastoma breakpoint family member 10 | qqrvgLAVDMDEIEKYqevee | 670 | 680 | 131.3 | 0.000632216 |
| 927 | Q86T75 | NBPF11 | Neuroblastoma breakpoint family member 11 | qqrigLAVDMDEIEKYqevee | 670 | 680 | 131.3 | 0.000632216 |
| 970 | Q8IYI0 | SHLD1 | Shieldin complex subunit 1 | ssaldLPSACDIRDYVlqgps | 20 | 30 | 131.3 | 0.000632216 |
| 971 | P51814 | ZNF41 | Zinc finger protein 41 | sileeLWQDNDQLEQRqenqn | 179 | 189 | 131.3 | 0.000633915 |
| 972 | Q96NX9 | DACH2 | Dachshund homolog 2 | tsdsgLRMLKDTGIPDieien | 557 | 567 | 131.3 | 0.000634481 |
| 973 | Q5THK1 | PRR14L | Protein PRR14L | shhppLEGRADVIADIqtipi | 694 | 704 | 131.3 | 0.000634575 |
| 975 | Q5SVQ8 | ZBTB41 | Zinc finger and BTB domain-containing protein 41 | krkdkLKYHIDHVHElkspdd | 739 | 749 | 131.3 | 0.000634764 |
| 974 | Q9BXX2 | ANKRD30B | Ankyrin repeat domain-containing protein 30B | nygnhLKERIDQYEKEkaere | 1358 | 1368 | 131.3 | 0.000634764 |
| 976 | P13611 | VCAN | Versican core protein | vyediLGMQTDIDTEVpseph | 2591 | 2601 | 131.3 | 0.000636274 |
| 977 | Q5THK1 | PRR14L | Protein PRR14L | ksvetLDQKADEVLDCqsnqn | 962 | 972 | 131.3 | 0.000638161 |
| 978 | O94875 | SORBS2 | Sorbin and SH3 domain-containing protein 2 | aldsaLKDICDQIKAEkkrgs | 735 | 745 | 131.3 | 0.00063901 |
| 980 | Q96NE9 | FRMD6 | FERM domain-containing protein 6 | yisdnLDLDMDQLEKRsrasg | 355 | 365 | 131.3 | 0.000639482 |
| 979 | Q9ULN7 | PNMA8B | Paraneoplastic antigen-like protein 8B | vdavvLRKAGDDGDLRecist | 402 | 412 | 131.3 | 0.000639482 |
| 981 | O60663 | LMX1B | LIM homeobox transcription factor 1-beta | dsdtsLTSLSDCFLGSsdvgs | 365 | 375 | 131.3 | 0.000640048 |
| 982 | Q52LD8 | RFTN2 | Raftlin-2 | ssdnvLRELDDGQFDQedgvt | 481 | 491 | 131.3 | 0.000640237 |
| 983 | Q13368 | MPP3 | MAGUK p55 subfamily member 3 | pvlppLPDNIDEDFDEesvki | 122 | 132 | 131.3 | 0.000641086 |
| 984 | Q96D15 | RCN3 | Reticulocalbin-3 | veanhLLHESDTDKDGrlska | 285 | 295 | 131.3 | 0.000641747 |
| 986 | P78344 | EIF4G2 | Eukaryotic translation initiation factor 4 gamma 2 | mdrdpLGGLADMFGQMpgsgi | 368 | 378 | 131.3 | 0.000641936 |
| 985 | Q9H0H0 | INTS2 | Integrator complex subunit 2 | nntigLVGQTDAPEVTreelk | 912 | 922 | 131.3 | 0.000641936 |
| 987 | Q9H8G2 | CAAP1 | Caspase activity and apoptosis inhibitor 1 | kmgsdLVSQQDICIDSassvr | 219 | 229 | 131.3 | 0.000642219 |
| 988 | P46939 | UTRN | Utrophin | ayletLKTLKDVLNDSenkaq | 916 | 926 | 131.3 | 0.000642785 |
| 989 | Q86UP3 | ZFHX4 | Zinc finger homeobox protein 4 | dqlkiLRAYFDINNSPseeqi | 2099 | 2109 | 131.3 | 0.000644483 |
| 990 | Q9H2F9 | CCDC68 | Coiled-coil domain-containing protein 68 | khcgnLQQGSDSEMDPsccsl | 73 | 83 | 131.2 | 0.000645238 |
| 991 | Q03721 | KCNC4 | Potassium voltage-gated channel subfamily C member 4 | lpgtrLAWLADPDGGGrpetd | 62 | 72 | 131.2 | 0.000648446 |
| 992 | O95294 | RASAL1 | RasGAP-activating-like protein 1 | rdqlrLKLLEDSNMDTtlead | 742 | 752 | 131.2 | 0.000648918 |
| 993 | Q13516 | OLIG2 | Oligodendrocyte transcription factor 2 | krmhdLNIAMDGLREVmpyah | 124 | 134 | 131.2 | 0.000648918 |
| 994 | O94880 | PHF14 | PHD finger protein 14 | ldfvsMEELNDMDDYDseddn | 197 | 207 | 131.2 | 0.000649296 |
| 995 | Q9Y5G9 | PCDHGA4 | Protocadherin gamma-A4 | rdlqlLMTASDSGDPPlssnv | 562 | 572 | 131.2 | 0.000650145 |
| 996 | O15083 | ERC2 | ERC protein 2 | qtqnrMKLMADNYDDDhhhyh | 909 | 919 | 131.2 | 0.0006509 |
| 997 | Q5JTC6 | AMER1 | APC membrane recruitment protein 1 | ksfdsLTGCGDIIAEQdmdsm | 325 | 335 | 131.2 | 0.0006509 |
| 998 | Q96QE4 | LRRC37B | Leucine-rich repeat-containing protein 37B | qeakaLNVEWDTDQQKtnyin | 862 | 872 | 131.2 | 0.000652221 |
| 999 | P17544 | ATF7 | Cyclic AMP-dependent transcription factor ATF-7 | pgslpLHLGYDPLHPTlpspt | 158 | 168 | 131.2 | 0.000652504 |
| 1000 | Q9Y4E8 | USP15 | Ubiquitin carboxyl-terminal hydrolase 15 | etegsLHCCKDQNINGngpng | 631 | 641 | 131.2 | 0.00065307 |
| 1001 | Q13395 | TARBP1 | Probable methyltransferase TARBP1 | avalaLGGGGDGDEAGpaeda | 172 | 182 | 131.2 | 0.000657694 |
| 1002 | Q9UHV2 | SERTAD1 | SERTA domain-containing protein 1 | ldeaeLDYLMDVLVGTqaler | 215 | 225 | 131.2 | 0.000657694 |
| 1003 | Q8TC71 | SPATA18 | Mitochondria-eating protein | eqwnsLKQNADQQDTEamsdy | 214 | 224 | 131.2 | 0.000659015 |
| 1004 | O94915 | FRYL | Protein furry homolog-like | snslrLSLIGDRRGDRrrsnt | 1944 | 1954 | 131.2 | 0.000660147 |
| 1005 | Q92800 | EZH1 | Histone-lysine N-methyltransferase EZH1 | tfieeLINNYDGKVHGeeemi | 150 | 160 | 131.2 | 0.000660619 |
| 1006 | P49588 | AARS1 | Alanine--tRNA ligase, cytoplasmic | etlksLKKVMDDLDRAskadv | 822 | 832 | 131.2 | 0.000660713 |
| 1007 | Q8N5D0 | WDTC1 | WD and tetratricopeptide repeats protein 1 | ditaaLFSKNDGEEKKgpggg | 483 | 493 | 131.2 | 0.000660713 |
| 1008 | O00178 | GTPBP1 | GTP-binding protein 1 | qtatiLSMDKDCLRTGdkatv | 525 | 535 | 131.2 | 0.000660902 |
| 1009 | Q7Z6I8 | C5orf24 | UPF0461 protein C5orf24 | gsikaLSRLADLGYGCgtaaf | 148 | 158 | 131.2 | 0.000660902 |
| 1010 | Q8IWJ2 | GCC2 | GRIP and coiled-coil domain-containing protein 2 | eqilyLQKQLDATTDEkketv | 201 | 211 | 131.2 | 0.000661185 |
| 1011 | Q9NVU0 | POLR3E | DNA-directed RNA polymerase III subunit RPC5 | qqkveLEMAIDTLNPNycrsk | 58 | 68 | 131.2 | 0.000661374 |
| 1012 | Q9Y4F3 | MARF1 | Meiosis regulator and mRNA stability factor 1 | teqelLRLTDDSPVDLlcapv | 1614 | 1624 | 131.2 | 0.000661657 |
| 1013 | Q9P227 | ARHGAP23 | Rho GTPase-activating protein 23 | flvghLKTIADHSEKNkmepr | 1039 | 1049 | 131.1 | 0.000661846 |
| 1014 | Q6UB99 | ANKRD11 | Ankyrin repeat domain-containing protein 11 | drpskLEKKNDLKEDKiskek | 776 | 786 | 131.1 | 0.000662695 |
| 1015 | A1L4H1 | SSC5D | Soluble scavenger receptor cysteine-rich domain-containing protein SSC5D | kcsgrLEVWHDQRWGTvcdds | 480 | 490 | 131.1 | 0.000663167 |
| 1016 | Q9NYQ8 | FAT2 | Protocadherin Fat 2 | dvttvMVNITDVNEHRpqfpq | 3308 | 3318 | 131.1 | 0.000663355 |
| 1017 | O60237 | PPP1R12B | Protein phosphatase 1 regulatory subunit 12B | laprkLNSTSDIEEKEnresa | 502 | 512 | 131.1 | 0.000663544 |
| 1018 | P55884 | EIF3B | Eukaryotic translation initiation factor 3 subunit B | nerleLRGGVDTDELDsnvdd | 778 | 788 | 131.1 | 0.000663544 |
| 1019 | Q9Y294 | ASF1A | Histone chaperone ASF1A | slnvmLESHMDCM--- | 197 | 204 | 131.1 | 0.000663733 |
| 1020 | Q9BSA9 | TMEM175 | Endosomal/lysosomal potassium channel TMEM175 | tpeqaLDTPGDCPPGRrdeda | 11 | 21 | 131.1 | 0.000663827 |
| 1023 | A0A087WUL | NBPF19 | Neuroblastoma breakpoint family member 19 | qqrvgLAIDMDEIEKYqevee | 399 | 409 | 131.1 | 0.00066411 |
| 1022 | A0A087WUL | NBPF19 | Neuroblastoma breakpoint family member 19 | qqcvgLAIDMDEIEKYqevee | 887 | 897 | 131.1 | 0.00066411 |
| 1021 | Q8N660 | NBPF15 | Neuroblastoma breakpoint family member 15 | qqrvgLAIDMDEIEKYqevee | 399 | 409 | 131.1 | 0.00066411 |
| 1024 | Q96LZ2 | MAGEB10 | Melanoma-associated antigen B10 | ggledLIDALDILEEEespp | 25 | 35 | 131.1 | 0.000664205 |
| 1026 | O75051 | PLXNA2 | Plexin-A2 | vkvwhLVKNHDHGDQKegdrg | 1654 | 1664 | 131.1 | 0.000664771 |
| 1025 | Q5TDP6 | LGSN | Lengsin | vgetdMSNSNDCMRDSsqilt | 50 | 60 | 131.1 | 0.000664771 |
| 1027 | Q9C0H6 | KLHL4 | Kelch-like protein 4 | kraqdLEMMADDNIEDstarl | 127 | 137 | 131.1 | 0.000664771 |
| 1028 | Q969G3 | SMARCE1 | SWI/SNF-related matrix-associated actin-dependent regulator of chromatin s | krlcgLKVEVDMEKIAaeiaq | 276 | 286 | 131.1 | 0.000665809 |
| 1029 | Q460N5 | PARP14 | Protein mono-ADP-ribosyltransferase PARP14 | pldggLDKMEDIPEECeniss | 134 | 144 | 131.1 | 0.000667224 |
| 1030 | Q8N8Z6 | DCBLD1 | Discoidin, CUB and LCCL domain-containing protein 1 | dgcghLVTYQDSGTMTsknyp | 44 | 54 | 131.1 | 0.000668073 |
| 1031 | Q6P9F5 | TRIM40 | E3 ubiquitin ligase TRIM40 | kklqaLQFQVDHGNHRleagp | 146 | 156 | 131.1 | 0.000669206 |
| 1032 | Q8IXY8 | PPIL6 | Probable inactive peptidyl-prolyl cis-trans isomerase-like 6 | erpihMCRITDSGDPYa | 300 | 310 | 131.1 | 0.000670716 |
| 1033 | Q8IY63 | AMOTL1 | Angiomotin-like protein 1 | vlhqeLQGYYDNADKLhkfek | 459 | 469 | 131.1 | 0.000670716 |
| 1034 | Q9C0J9 | BHLHE41 | Class E basic helix-loop-helix protein 41 | qpsaeLAAENDTDTDSgygge | 230 | 240 | 131.1 | 0.000670904 |
| 1035 | Q05DH4 | FAM160A1 | Protein FAM160A1 | peseeLIAQYDQIIKEldsga | 751 | 761 | 131.1 | 0.000672697 |
| 1036 | Q8WYN3 | CSRNP3 | Cysteine/serine-rich nuclear protein 3 | ansstLYYQIDSHIPGtpnqi | 435 | 445 | 131.1 | 0.000672697 |
| 1037 | Q9P0X4 | CACNA1I | Voltage-dependent T-type calcium channel subunit alpha-1I | dvftkMGDRGDRGEDEeeidy | 1109 | 1119 | 131.1 | 0.00067449 |
| 1038 | Q9NQC3 | RTN4 | Reticulon-4 | tsnpfLVAAQDSETDYvttdn | 519 | 529 | 131.0 | 0.00067666 |
| 1039 | Q9Y247 | FAM50B | Protein FAM50B | rkiscLSFALDDLDDQadaae | 120 | 130 | 131.0 | 0.000676849 |
| 1040 | Q2M243 | CCDC27 | Coiled-coil domain-containing protein 27 | gviasLQQQVDFQETQlrkin | 446 | 456 | 131.0 | 0.000677793 |
| 1041 | Q9HC44 | GPBP1L1 | Vasculin-like protein 1 | fnngpLRTAGDSWHQPslfrh | 59 | 69 | 131.0 | 0.000678547 |
| 1042 | Q9UIF8 | BAZ2B | Bromodomain adjacent to zinc finger domain protein 2B | nnqvlLHGISDPKADGqkate | 303 | 313 | 131.0 | 0.000678925 |
| 1043 | Q15643 | TRIP11 | Thyroid receptor-interacting protein 11 | ehlikLNQKKDMEIAElkkni | 767 | 777 | 131.0 | 0.000679208 |
| 1045 | Q9NRY4 | ARHGAP35 | Rho GTPase-activating protein 35 | semesLQRQFDQDHNLdlaek | 1295 | 1305 | 131.0 | 0.000679302 |
| 1044 | Q9Y2I6 | NINL | Ninein-like protein | eeledLFNKLDQDGDGkvsle | 241 | 251 | 131.0 | 0.000679302 |
| 1047 | Q00536 | CDK16 | Cyclin-dependent kinase 16 | ivhedLKMGSDGESDQasats | 74 | 84 | 131.0 | 0.000679869 |
| 1046 | Q00537 | CDK17 | Cyclin-dependent kinase 17 | ivhenLKMGSDGESDQasgts | 101 | 111 | 131.0 | 0.000679869 |
| 1049 | P05549 | TFAP2A | Transcription factor AP-2-alpha | mLWKLTDNIKYEdcedr | 2 | 12 | 131.0 | 0.000680152 |
| 1048 | Q05D60 | DEUP1 | Deuterosome assembly protein 1 | aelqeLMEQIDIMVSNkkmdw | 21 | 31 | 131.0 | 0.000680152 |
| 1050 | Q9C0A1 | ZFHX2 | Zinc finger homeobox protein 2 | elvrhLKKCYDDQTLEeeeee | 1689 | 1699 | 131.0 | 0.000680246 |
| 1051 | Q68DX3 | FRMPD2 | FERM and PDZ domain-containing protein 2 | aagglLSTSMDNFNVDgskea | 699 | 709 | 131.0 | 0.00068119 |

| | | | | | | | | |
|---|---|---|---|---|---|---|---|---|
| 1052 | Q8WW24 | TEKT4 | Tektin-4 | hlhktLREITDQEHNVaalkq | 314 | 324 | 131.0 | 0.000681378 |
| 1053 | Q96JJ3 | ELMO2 | Engulfment and cell motility protein 2 | lleerMMTKMDPNDQAqrdii | 301 | 311 | 131.0 | 0.000682039 |
| 1054 | Q4G0P3 | HYDIN | Hydrocephalus-inducing protein homolog | vkrppLTMTDDLEHFVfvipp | 2627 | 2637 | 131.0 | 0.000682888 |
| 1055 | P53814 | SMTN | Smoothelin | rlagqLESMNDVEELTallrs | 86 | 96 | 131.0 | 0.000682982 |
| 1056 | Q4AC94 | C2CD3 | C2 domain-containing protein 3 | lalepLSETYDSYHPLpttdm | 169 | 179 | 131.0 | 0.000682982 |
| 1057 | O15197 | EPHB6 | Ephrin type-B receptor 6 | rkpdtLQAGGDPGERPsqall | 925 | 935 | 131.0 | 0.000683266 |
| 1058 | P20929 | NEB | Nebulin | apdhhLSTYSDGGVFAvstay | 6563 | 6573 | 131.0 | 0.000683737 |
| 1059 | Q2LD37 | KIAA1109 | Transmembrane protein KIAA1109 | eqpipLWNEHDGTADGdkpki | 3137 | 3147 | 131.0 | 0.000684303 |
| 1060 | A7E2V4 | ZSWIM8 | Zinc finger SWIM domain-containing protein 8 | ddqakLKKILDKLLDResqth | 1004 | 1014 | 131.0 | 0.000684681 |
| 1061 | Q12968 | NFATC3 | Nuclear factor of activated T-cells, cytoplasmic 3 | sscesLSHIYDDVDSElneaa | 185 | 195 | 131.0 | 0.000684964 |
| 1062 | O15079 | SNPH | Syntaphilin | keikqLKQVIDTVKNNlidkd | 145 | 155 | 131.0 | 0.000685436 |
| 1063 | Q96I24 | FUBP3 | Far upstream element-binding protein 3 | gadkpLRITGDAFKVQqarem | 212 | 222 | 131.0 | 0.00068553 |
| 1065 | Q8IWN7 | RP1L1 | Retinitis pigmentosa 1-like 1 protein | prrilLIKNMDPRLQQtvvls | 156 | 166 | 131.0 | 0.000685719 |
| 1064 | Q9UNK9 | ANGEL1 | Protein angel homolog 1 | laqssLALLMDNPGEEnaase | 91 | 101 | 131.0 | 0.000685719 |
| 1066 | Q5BKU9 | OXLD1 | Oxidoreductase-like domain-containing protein 1 | yadrlLQHFQDGGERAlaale | 108 | 118 | 131.0 | 0.000686474 |
| 1067 | Q01970 | PLCB3 | 1-phosphatidylinositol 4,5-bisphosphate phosphodiesterase beta-3 | lrpgaLGGAADVEDTKegede | 1012 | 1022 | 131.0 | 0.000687984 |
| 1068 | P25054 | APC | Adenomatous polyposis coli protein | ameeqLGTCQDMEKRAqrria | 204 | 214 | 131.0 | 0.000690248 |
| 1069 | Q96KR7 | PHACTR3 | Phosphatase and actin regulator 3 | errniLKQRNDQTEQEerrei | 460 | 470 | 131.0 | 0.000690248 |
| 1070 | Q08999 | RBL2 | Retinoblastoma-like protein 2 | ttrrrLFVENDSPSDGgtpgr | 682 | 692 | 131.0 | 0.000690437 |
| 1071 | Q9H8M5 | CNNM2 | Metal transporter CNNM2 | kieltLTELHDGLPDEtanll | 839 | 849 | 131.0 | 0.00069072 |
| 1072 | Q13772 | NCOA4 | Nuclear receptor coactivator 4 | enlgnLKCLNDHLEAKkplst | 366 | 376 | 131.0 | 0.000691097 |
| 1073 | Q9UQC2 | GAB2 | GRB2-associated-binding protein 2 | efgdlLVDNMDVPATPlsayq | 310 | 320 | 131.0 | 0.000691569 |
| 1075 | P46939 | UTRN | Utrophin | eklrdLQGAMDDLDADmkeae | 2696 | 2706 | 130.9 | 0.000692607 |
| 1074 | Q14CN4 | KRT72 | Keratin, type II cytoskeletal 72 | sllapLNVEMDPEIQRvraqe | 110 | 120 | 130.9 | 0.000692607 |
| 1076 | Q9NQL2 | RRAGD | Ras-related GTP-binding protein D | eeedeLVGLADYGDGPdssda | 25 | 35 | 130.9 | 0.000692607 |
| 1077 | Q96CA5 | BIRC7 | Baculoviral IAP repeat-containing protein 7 | ethsqLLGSWDPWEEPedaap | 173 | 183 | 130.9 | 0.00069289 |
| 1079 | P29377 | S100G | Protein S100-G | ntlddLFQELDKNGDGevsfe | 53 | 63 | 130.9 | 0.000693362 |
| 1078 | Q7Z333 | SETX | Probable helicase senataxin | yklieLIHGKDTVEVEedsvs | 1522 | 1532 | 130.9 | 0.000693362 |
| 1080 | Q96GU1 | PAGE5 | P antigen family member 5 | qqelaLLKIEDAPGDGpdvre | 96 | 106 | 130.9 | 0.000693551 |
| 1081 | Q68DQ2 | CRYBG3 | Very large A-kinase anchor protein | svserLKMNFDEDDREaadee | 2134 | 2144 | 130.9 | 0.000694023 |
| 1083 | O75157 | TSC22D2 | TSC22 domain family protein 2 | sseetLNNVGDAETPGtvspn | 72 | 82 | 130.9 | 0.000694117 |
| 1082 | O94972 | TRIM37 | E3 ubiquitin-protein ligase TRIM37 | agslsLRRAVDPGENSrskgd | 774 | 784 | 130.9 | 0.000694117 |
| 1084 | Q71F56 | MED13L | Mediator of RNA polymerase II transcription subunit 13-like | dmfvdLPFPDDMDNDIgilmt | 2029 | 2039 | 130.9 | 0.000695627 |
| 1085 | O75177 | SS18L1 | Calcium-responsive transactivator | rnlvyLATIADSNQNMqsllp | 61 | 71 | 130.9 | 0.000696099 |
| 1086 | Q15532 | SS18 | Protein SSXT | tnlvyLATIADSNQNMqsllp | 61 | 71 | 130.9 | 0.000696099 |
| 1087 | Q5UIP0 | RIF1 | Telomere-associated protein RIF1 | nfhlgLKEDNDTINDSlivse | 1800 | 1810 | 130.9 | 0.000697042 |
| 1088 | Q8WXI7 | MUC16 | Mucin-16 | ptqerLTTYKDTAHTEamhas | 5698 | 5708 | 130.9 | 0.000697325 |
| 1089 | Q8WXI7 | MUC16 | Mucin-16 | tsherLTTYKDTAHTEavhps | 7754 | 7764 | 130.9 | 0.000697325 |
| 1090 | O94911 | ABCA8 | ABC-type organic anion transporter ABCA8 | tdhvaLEDEMDADPSFhdsfe | 453 | 463 | 130.9 | 0.00069808 |
| 1091 | Q8N7Z5 | ANKRD31 | Ankyrin repeat domain-containing protein 31 | raptfLMNQTDTHIVEkmakn | 1045 | 1055 | 130.9 | 0.000698835 |
| 1092 | Q6ZMV9 | KIF6 | Kinesin-like protein KIF6 | dsdsrLEVGADMRKVHhcfhh | 412 | 422 | 130.9 | 0.000699118 |
| 1093 | Q5T0N5 | FNBP1L | Formin-binding protein 1-like | dqkdaLNKMKDVYEKNpqmgd | 428 | 438 | 130.9 | 0.000699307 |
| 1094 | Q96QT4 | TRPM7 | Transient receptor potential cation channel subfamily M member 7 | krvssLAGFTDCHRTSipvhs | 1478 | 1488 | 130.9 | 0.000701288 |
| 1095 | Q9ULJ7 | ANKRD50 | Ankyrin repeat domain-containing protein 50 | vvdllLEGGADVDHTDnngrt | 768 | 778 | 130.9 | 0.000701477 |
| 1098 | O94910 | ADGRL1 | Adhesion G protein-coupled receptor L1 | lrpeaLESWKDMNATEqvhta | 633 | 643 | 130.9 | 0.000702798 |
| 1097 | P61011 | SRP54 | Signal recognition particle 54 kDa protein | ggikgLFKGGDMSKNVsqsqm | 440 | 450 | 130.9 | 0.000702798 |
| 1096 | Q9H6N6 | MYH16 | Putative uncharacterized protein MYH16 | kaesdLKMTIDNLNEMerskl | 251 | 261 | 130.9 | 0.000702798 |
| 1099 | Q92994 | BRF1 | Transcription factor IIIB 90 kDa subunit | dgeldLSGIDDLEIDRyilne | 449 | 459 | 130.9 | 0.000703364 |
| 1100 | Q6V0I7 | FAT4 | Protocadherin Fat 4 | sstsvLVTVTDVNDNPprfqh | 2453 | 2463 | 130.9 | 0.000704025 |
| 1101 | Q9NTJ3 | SMC4 | Structural maintenance of chromosomes protein 4 | keekkLKEVMDSLKQEtqglq | 460 | 470 | 130.9 | 0.000704119 |
| 1102 | Q8N5U6 | RNF10 | RING finger protein 10 | vsketLEMFSDDIEKRkrqrq | 590 | 600 | 130.9 | 0.000704497 |
| 1103 | Q9NTX7 | RNF146 | E3 ubiquitin-protein ligase RNF146 | kgvagLRLDCDANTVNlares | 181 | 191 | 130.9 | 0.000704968 |
| 1104 | P86452 | ZBED6 | Zinc finger BED domain-containing protein 6 | adapaLLASNDPEQDEeslfe | 101 | 111 | 130.9 | 0.000705063 |
| 1107 | A0A096LP55 | UQCRHL | Cytochrome b-c1 complex subunit 6-like, mitochondrial | deqkmLTESGDPEEEEeeeee | 10 | 20 | 130.9 | 0.000705252 |
| 1106 | P07919 | UQCRH | Cytochrome b-c1 complex subunit 6, mitochondrial | deqkmLTESGDPEEEEeeeee | 10 | 20 | 130.9 | 0.000705252 |
| 1105 | Q9H081 | MIS12 | Protein MIS12 homolog | qnsrkLQNIRDNVEKEskrlk | 188 | 198 | 130.9 | 0.000705252 |
| 1109 | Q96A49 | SYAP1 | Synapse-associated protein 1 | saeeeLQQAGDQELLHqakdf | 42 | 52 | 130.9 | 0.000705629 |
| 1108 | Q9Y2D1 | ATF5 | Cyclic AMP-dependent transcription factor ATF-5 | llkkeLEQMEDFFLDApplpp | 109 | 119 | 130.9 | 0.000705629 |
| 1110 | Q96KM6 | ZNF512B | Zinc finger protein 512B | lrdipLSLMNDWKDEFkahsr | 89 | 99 | 130.8 | 0.000707894 |
| 1113 | B7ZAP0 | RABGAP1L | Rab GTPase-activating protein 1-like, isoform 10 | tskiaLRNDLDQAEDKadvln | 44 | 54 | 130.8 | 0.000707988 |
| 1111 | Q92636 | NSMAF | Protein FAN | eggvdLNSIQDPDEKVamltq | 541 | 551 | 130.8 | 0.000707988 |
| 1112 | Q9BWU0 | SLC4A1AP | Kanadaptin | elpptLMRMKDEPEVEeeeee | 638 | 648 | 130.8 | 0.000707988 |
| 1114 | Q08AD1 | CAMSAP2 | Calmodulin-regulated spectrin-associated protein 2 | rkqlkLMEDMDTVIKPrpqvv | 1244 | 1254 | 130.8 | 0.000708365 |
| 1115 | Q9H2J1 | ARRDC1-AS1 | Uncharacterized protein ARRDC1-AS1 | stepcLSGLGDVGRGEaaslr | 113 | 123 | 130.8 | 0.000708365 |
| 1116 | Q5THK1 | PRR14L | Protein PRR14L | lckpnLVCEADDNHQQlhghh | 304 | 314 | 130.8 | 0.00070912 |
| 1118 | A2CJ06 | DYTN | Dystrotelin | trihkLKTNQDSLWTKlqqir | 361 | 371 | 130.8 | 0.000709875 |
| 1119 | Q5U651 | RASIP1 | Ras-interacting protein 1 | lgehfLFMYKDPRTGGsgpar | 484 | 494 | 130.8 | 0.000709875 |
| 1117 | Q9Y6R1 | SLC4A4 | Electrogenic sodium bicarbonate cotransporter 1 | tkksnLRSLADIGKTVssasr | 221 | 231 | 130.8 | 0.000709875 |
| 1120 | Q8N7Z5 | ANKRD31 | Ankyrin repeat domain-containing protein 31 | qkendLTEATDKDHEFyvssp | 1565 | 1575 | 130.8 | 0.00071063 |
| 1121 | Q00325 | SLC25A3 | Phosphate carrier protein, mitochondrial | fntphLQLVHDGLGDLrsssp | 19 | 29 | 130.8 | 0.000711196 |
| 1123 | Q15691 | MAPRE1 | Microtubule-associated protein RP/EB family member 1 | endpvLQRIVDILYATdegfv | 239 | 249 | 130.8 | 0.000711196 |
| 1122 | Q68DQ2 | CRYBG3 | Very large A-kinase anchor protein | ktekdLQNPSDHHEDGikrer | 156 | 166 | 130.8 | 0.000711196 |
| 1124 | Q8NBJ4 | GOLM1 | Golgi membrane protein 1 | rnqqkLRGEDDYNMDEneaes | 345 | 355 | 130.8 | 0.000711951 |
| 1125 | Q9P2K8 | EIF2AK4 | eIF-2-alpha kinase GCN2 | latdhLAFSADSKQDDqtgdl | 874 | 884 | 130.8 | 0.000711951 |
| 1126 | O14974 | PPP1R12A | Protein phosphatase 1 regulatory subunit 12A | tiprrLASTSDIEEKEnrdss | 505 | 515 | 130.8 | 0.000712612 |
| 1127 | Q08174 | PCDH1 | Protocadherin-1 | etavaLVQVSDRDEGEnaavt | 416 | 426 | 130.8 | 0.000712612 |
| 1130 | P18206 | VCL | Vinculin | gtckmLGQMTDQVADLrargq | 328 | 338 | 130.8 | 0.000713083 |
| 1129 | Q13342 | SP140 | Nuclear body protein SP140 | gvsckLAIQIDEGESEempkl | 226 | 236 | 130.8 | 0.000713083 |
| 1128 | Q14687 | GSE1 | Genetic suppressor element 1 | frqqvLEQHLDMGRPPvpaea | 523 | 533 | 130.8 | 0.000713083 |
| 1132 | Q15916 | ZBTB6 | Zinc finger and BTB domain-containing protein 6 | nqhtdLCQSSDPDVKNedens | 142 | 152 | 130.8 | 0.000713178 |
| 1131 | Q68G74 | LHX8 | LIM/homeobox protein Lhx8 | tmltaLHSYMDAHSPTtlglq | 326 | 336 | 130.8 | 0.000713178 |
| 1133 | Q14126 | DSG2 | Desmoglein-2 | laevcLGQKIDINKEleqrqk | 841 | 851 | 130.8 | 0.000713272 |
| 1134 | O15021 | MAST4 | Microtubule-associated serine/threonine-protein kinase 4 | asalsLMIPGDMFAVSplgsp | 1094 | 1104 | 130.8 | 0.000713461 |
| 1135 | Q3B7T3 | BEAN1 | Protein BEAN1 | dwpppLDISSDGDVDAtvlre | 124 | 134 | 130.8 | 0.000714405 |
| 1136 | P82094 | TMF1 | TATA element modulatory factor | kekalLEEAFDNLKDEmfrvk | 475 | 485 | 130.8 | 0.000715065 |
| 1137 | Q9NRJ5 | PAPOLB | Poly(A) polymerase beta | tegrrLTDLNDSSFDLsagce | 515 | 525 | 130.8 | 0.000715631 |
| 1138 | Q5CZC0 | FSIP2 | Fibrous sheath-interacting protein 2 | kskrnLGTTTDTLEIRirtss | 5638 | 5648 | 130.8 | 0.000716575 |
| 1139 | Q5FWF4 | ZRANB3 | DNA annealing helicase and endonuclease ZRANB3 | mqidsLNHIQDKNEKDdsqkd | 658 | 668 | 130.8 | 0.000716575 |

| | | | | | | | | |
|---|---|---|---|---|---|---|---|---|
| 1140 | O94988 | FAM13A | Protein FAM13A | rrsssLGSYDDEQEDLtpaql | 653 | 663 | 130.8 | 0.00071733 |
| 1141 | O95159 | ZFPL1 | Zinc finger protein-like 1 | drtpgLHGDCDDDKYRrrpal | 226 | 236 | 130.8 | 0.000717613 |
| 1142 | P21333 | FLNA | Filamin-A | rlspfMADIRDAPQDFhpdrv | 660 | 670 | 130.8 | 0.000718273 |
| 1143 | Q9Y4D8 | HECTD4 | Probable E3 ubiquitin-protein ligase HECTD4 | kdpqgLGVTSDAIADAcqalv | 2953 | 2963 | 130.8 | 0.000718556 |
| 1144 | O15085 | ARHGEF11 | Rho guanine nucleotide exchange factor 11 | qltlkLNRLKDMELAHrellk | 1436 | 1446 | 130.8 | 0.000718839 |
| 1146 | O43513 | MED7 | Mediator of RNA polymerase II transcription subunit 7 | ekdaaLCVLIDEMNERp | 222 | 232 | 130.8 | 0.000719406 |
| 1145 | Q14865 | ARID5B | AT-rich interactive domain-containing protein 5B | fsakpLASRVDPEKDNetdqg | 508 | 518 | 130.8 | 0.000719406 |
| 1147 | Q92610 | ZNF592 | Zinc finger protein 592 | pdfddLLAAFDIPDPTsldak | 12 | 22 | 130.8 | 0.000719594 |
| 1148 | Q96AP0 | ACD | Adrenocortical dysplasia protein homolog | lmawaLHFLMDAQPGSeptpm | 443 | 453 | 130.8 | 0.000720066 |
| 1149 | Q8N4X5 | AFAP1L2 | Actin filament-associated protein 1-like 2 | agpvtLGTTVDTTHLEnvspr | 754 | 764 | 130.8 | 0.000720349 |
| 1150 | Q5JRA6 | MIA3 | Transport and Golgi organization protein 1 homolog | fdelpLLTFTDGEDMKtpaks | 330 | 340 | 130.8 | 0.000721482 |
| 1151 | Q9H9B1 | EHMT1 | Histone-lysine N-methyltransferase EHMT1 | kvllmLVDGIDPNFKMehqnk | 758 | 768 | 130.8 | 0.000723558 |
| 1155 | P01889 | HLA-B | HLA class I histocompatibility antigen, B alpha chain | paeitLTWQRDGEDQTqdtel | 239 | 249 | 130.8 | 0.000724595 |
| 1153 | P01893 | HLA-H | Putative HLA class I histocompatibility antigen, alpha chain H | paeitLTWQRDGEDQTqdtel | 239 | 249 | 130.8 | 0.000724595 |
| 1156 | P04439 | HLA-A | HLA class I histocompatibility antigen, A alpha chain | paeitLTWQRDGEDQTqdtel | 239 | 249 | 130.8 | 0.000724595 |
| 1154 | P10321 | HLA-C | HLA class I histocompatibility antigen, C alpha chain | paeitLTWQRDGEDQTqdtel | 239 | 249 | 130.8 | 0.000724595 |
| 1152 | Q8N1D5 | C1orf158 | Uncharacterized protein C1orf158 | pphryLISTYDDHYNRhgynp | 94 | 104 | 130.8 | 0.000724595 |
| 1158 | Q5CZC0 | FSIP2 | Fibrous sheath-interacting protein 2 | kmayhLQKMQDTGFNGedigk | 303 | 313 | 130.7 | 0.000725256 |
| 1157 | Q96NE9 | FRMD6 | FERM domain-containing protein 6 | mneesLEVSPDMCIYItedml | 473 | 483 | 130.7 | 0.000725256 |
| 1159 | Q9BY89 | KIAA1671 | Uncharacterized protein KIAA1671 | esrspLEDETDNTWMFkdste | 1677 | 1687 | 130.7 | 0.000725256 |
| 1160 | Q15642 | TRIP10 | Cdc42-interacting protein 4 | dqreaLKKMKDVYEKTpqmgd | 424 | 434 | 130.7 | 0.000726294 |
| 1161 | Q9BV35 | SLC25A23 | Calcium-binding mitochondrial carrier protein SCaMC-3 | qrwgrLFEELDSNKDGrvdvh | 17 | 27 | 130.7 | 0.000726954 |
| 1162 | Q96T51 | RUFY1 | RUN and FYVE domain-containing protein 1 | ctvgdLQTKIDGLEKTnsklq | 325 | 335 | 130.7 | 0.000727521 |
| 1163 | P53814 | SMTN | Smoothelin | edegvLDKMLDQSTDFeerkl | 592 | 602 | 130.7 | 0.000727709 |
| 1164 | P55317 | FOXA1 | Hepatocyte nuclear factor 3-alpha | phesqLHLKGDPHYSFnhpfs | 386 | 396 | 130.7 | 0.000729125 |
| 1165 | Q7Z3G6 | PRICKLE2 | Prickle-like protein 2 | advdpLSLQMDMLSLSsqtps | 372 | 382 | 130.7 | 0.000731295 |
| 1166 | P07359 | GP1BA | Platelet glycoprotein Ib alpha chain | kgcptLGDEGDTDLYDyypee | 283 | 293 | 130.7 | 0.000731578 |
| 1167 | Q6UXY8 | TMC5 | Transmembrane channel-like protein 5 | dpvgsLWGENDYPEGIemasm | 286 | 296 | 130.7 | 0.000731956 |
| 1168 | Q8NFC6 | BOD1L1 | Biorientation of chromosomes in cell division protein 1-like 1 | pvettLKMKDDSKTDTgivtv | 2887 | 2897 | 130.7 | 0.000733088 |
| 1169 | Q01433 | AMPD2 | AMP deaminase 2 | hfpldLRTSMDGKCKEiaeel | 115 | 125 | 130.7 | 0.000734315 |
| 1170 | Q8IWP9 | CCDC28A | Coiled-coil domain-containing protein 28A | iqhsfLTDVSDVQEMErglls | 169 | 179 | 130.7 | 0.000734315 |
| 1171 | Q8N163 | CCAR2 | Cell cycle and apoptosis regulator protein 2 | ldpelLLLRDDGEEEFagakl | 653 | 663 | 130.7 | 0.000735164 |
| 1172 | P42858 | HTT | Huntingtin | iathhLYQAWDPVPSLspatt | 2573 | 2583 | 130.7 | 0.000735353 |
| 1174 | Q7Z2Z1 | TICRR | Treslin | aeelhLVADVDPGEGRppitg | 392 | 402 | 130.7 | 0.000735353 |
| 1173 | Q8TDI0 | CHD5 | Chromodomain-helicase-DNA-binding protein 5 | aisklLDRNQDATDDTelqnm | 1266 | 1276 | 130.7 | 0.000735353 |
| 1176 | O94986 | CEP152 | Centrosomal protein of 152 kDa | klivpLSSQQDSGFDSpfvnl | 1694 | 1704 | 130.7 | 0.000735919 |
| 1175 | P02462 | COL4A1 | Collagen alpha-1(IV) chain | efyfdLRLKGDKGDPGfpgqp | 537 | 547 | 130.7 | 0.000735919 |
| 1177 | P25963 | NFKBIA | NF-kappa-B inhibitor alpha | kkerlLDDRHDSGLDSmkdee | 26 | 36 | 130.7 | 0.000736485 |
| 1178 | Q02505 | MUC3A | Mucin-3A | siqttLTTYMDTSSMMpeses | 2825 | 2835 | 130.7 | 0.000736957 |
| 1179 | Q96AJ1 | CLUAP1 | Clusterin-associated protein 1 | kteeeLQKQYDTYLEKfqnlt | 252 | 262 | 130.7 | 0.000738183 |
| 1180 | O15047 | SETD1A | Histone-lysine N-methyltransferase SETD1A | sskcsLYADSDGENDStsdse | 1019 | 1029 | 130.7 | 0.000739316 |
| 1181 | Q7RTU3 | OLIG3 | Oligodendrocyte transcription factor 3 | krmhdLNLAMDGLREVmpyah | 99 | 109 | 130.7 | 0.000739316 |
| 1182 | Q92552 | MRPS27 | 28S ribosomal protein S27, mitochondrial | qgsekLVEQLDIEETEqsklp | 313 | 323 | 130.7 | 0.000740354 |
| 1183 | Q96G01 | BICD1 | Protein bicaudal D homolog 1 | pqcsqLAGRQDCPTVSpdtal | 937 | 947 | 130.7 | 0.000741203 |
| 1185 | Q641Q2 | WASHC2A | WASH complex subunit 2A | lfshkLQKDNDPDVDLfagtk | 851 | 861 | 130.7 | 0.000741392 |
| 1184 | Q9Y4E1 | WASHC2C | WASH complex subunit 2C | lfshkLQKDNDPDVDLfagtk | 851 | 861 | 130.7 | 0.000741392 |
| 1186 | Q6PIW4 | FIGNL1 | Fidgetin-like protein 1 | eidslLSQRGDGEHESsrrik | 506 | 516 | 130.7 | 0.000742147 |
| 1187 | Q9C0D2 | CEP295 | Centrosomal protein of 295 kDa | ieqqkLRLETDCFRAQleeek | 534 | 544 | 130.7 | 0.000742335 |
| 1188 | Q8N1N4 | KRT78 | Keratin, type II cytoskeletal 78 | aldaeLKACRDQEEEYkskye | 187 | 197 | 130.6 | 0.000743468 |
| 1189 | Q15293 | RCN1 | Reticulocalbin-1 | aearhLVYESDKNKDEkltke | 288 | 298 | 130.6 | 0.000743656 |
| 1190 | Q8TAP6 | CEP76 | Centrosomal protein of 76 kDa | dvmkeLNFVTDSVEQElpssp | 69 | 79 | 130.6 | 0.000744506 |
| 1191 | Q8NET4 | RTL9 | Retrotransposon Gag-like protein 9 | krcyyLKEHGDPQEGLhdhlg | 1360 | 1370 | 130.6 | 0.000745449 |
| 1192 | P82094 | TMF1 | TATA element modulatory factor | kdlgrLQVDMDELEEKnrsiq | 642 | 652 | 130.6 | 0.000745638 |
| 1193 | O60422 | ONECUT3 | One cut domain family member 3 | qhlppLAAVADKFHQHaaaaa | 117 | 127 | 130.6 | 0.000745827 |
| 1194 | Q14966 | ZNF638 | Zinc finger protein 638 | nslftLDELIDQDDCIshsep | 1645 | 1655 | 130.6 | 0.000747053 |
| 1197 | Q8N2C7 | UNC80 | Protein unc-80 homolog | letssLLQHGDTVLHIseeng | 3214 | 3224 | 130.6 | 0.000747053 |
| 1195 | Q9BZA7 | PCDH11X | Protocadherin-11 X-linked | hgtvgLITVTDPDYGDnsavt | 591 | 601 | 130.6 | 0.000747053 |
| 1196 | Q9BZA8 | PCDH11Y | Protocadherin-11 Y-linked | hgtvgLITVTDPDYGDnsavt | 623 | 633 | 130.6 | 0.000747053 |
| 1198 | O14776 | TCERG1 | Transcription elongation regulator 1 | drpddLIGRADVDKIIqepph | 562 | 572 | 130.6 | 0.000747525 |
| 1199 | Q86UP3 | ZFHX4 | Zinc finger homeobox protein 4 | lppqlLQYQCDQCTVAfptle | 2446 | 2456 | 130.6 | 0.000747525 |
| 1200 | Q96K83 | ZNF521 | Zinc finger protein 521 | htveeLYSHMDSHQQPescnh | 325 | 335 | 130.6 | 0.000747714 |
| 1201 | Q8TDM6 | DLG5 | Disks large homolog 5 | eemeaLRQIKDTVTMDagran | 477 | 487 | 130.6 | 0.000747808 |
| 1202 | A6NL46 | | Putative UPF0607 protein ENSP00000332738 | qqtppLQLRWDRDEGPppakl | 264 | 274 | 130.6 | 0.000748374 |
| 1203 | A8MX80 | | Putative UPF0607 protein ENSP00000383144 | qptppLQLRWDRDEGPppakl | 265 | 275 | 130.6 | 0.000748374 |
| 1204 | Q92882 | OSTF1 | Osteoclast-stimulating factor 1 | davrtLSNAEDYLDDEdsd | 201 | 211 | 130.6 | 0.000748846 |
| 1205 | Q70EL2 | USP45 | Ubiquitin carboxyl-terminal hydrolase 45 | qknenLEMNGDSLMFAslmns | 463 | 473 | 130.6 | 0.000750262 |
| 1206 | Q8NEV8 | EXPH5 | Exophilin-5 | grkkpLTSGMDASELTprawe | 1143 | 1153 | 130.6 | 0.000750262 |
| 1207 | O43396 | TXNL1 | Thioredoxin-like protein 1 | kifinLPRSMDFEEAErsept | 200 | 210 | 130.6 | 0.000750922 |
| 1208 | O95670 | ATP6V1G2 | V-type proton ATPase subunit G 2 | vlaqlLGMVCDVRPQVhpnyr | 100 | 110 | 130.6 | 0.000751771 |
| 1209 | P35579 | MYH9 | Myosin-9 | evkvlLQGKGDSEHKRkkvea | 1237 | 1247 | 130.6 | 0.000751771 |
| 1210 | O76041 | NEBL | Nebulette | ikgkgMQVSMDIPDILrakrt | 536 | 546 | 130.6 | 0.00075196 |
| 1211 | Q5D1E8 | ZC3H12A | Endoribonuclease ZC3H12A | svlqkLGVQADTNTVLgelvk | 72 | 82 | 130.6 | 0.000753092 |
| 1213 | Q9C0D5 | TANC1 | Protein TANC1 | pnqggLATCSDVRHPAsltss | 1648 | 1658 | 130.6 | 0.000753942 |
| 1212 | Q9NYF5 | FAM13B | Protein FAM13B | iphldLKNVSDGDKWEepfpa | 470 | 480 | 130.6 | 0.000753942 |
| 1214 | Q9H3E2 | SNX25 | Sorting nexin-25 | eedsdLSDYGDDVDGRkdala | 667 | 677 | 130.6 | 0.00075781 |
| 1215 | O60721 | SLC24A1 | Sodium/potassium/calcium exchanger 1 | ttpttLKGMFDSTPTFlthev | 249 | 259 | 130.6 | 0.000758377 |
| 1216 | Q2NKX8 | ERCC6L | DNA excision repair protein ERCC-6-like | linmvLDHVEDMEERLddsse | 1104 | 1114 | 130.6 | 0.000758377 |
| 1217 | O15015 | ZNF646 | Zinc finger protein 646 | gaegnLESDGDCLQAEsegdk | 723 | 733 | 130.6 | 0.000759131 |
| 1218 | P51692 | STAT5B | Signal transducer and activator of transcription 5B | rveelLGRPMDSQWIPhaqs | 773 | 783 | 130.6 | 0.000761019 |
| 1219 | Q9HC29 | NOD2 | Nucleotide-binding oligomerization domain-containing protein 2 | sqspkLHGCWDPHSLHpardl | 119 | 129 | 130.6 | 0.000762528 |
| 1220 | Q9Y2W2 | WBP11 | WW domain-binding protein 11 | qkraqLSQYFDAVKNAqhvev | 121 | 131 | 130.6 | 0.000762906 |
| 1221 | Q7Z2Z1 | TICRR | Treslin | hasspLLITSDTEHVTllsea | 1455 | 1465 | 130.6 | 0.000763095 |
| 1222 | Q9BRD0 | BUD13 | BUD13 homolog | gdkkqLDSKGDCQKATdsdls | 342 | 352 | 130.6 | 0.000763566 |
| 1223 | Q9BXX3 | ANKRD30A | Ankyrin repeat domain-containing protein 30A | edstsLSKILDTVHSCerare | 961 | 971 | 130.6 | 0.000763566 |
| 1224 | O60393 | NOBOX | Homeobox protein NOBOX | ltspdLEGTWDTRDKDgfkaq | 12 | 22 | 130.5 | 0.000764793 |
| 1225 | P07814 | EPRS1 | Bifunctional glutamate/proline--tRNA ligase | nfkgkLIMRFDDTNPEkeked | 232 | 242 | 130.5 | 0.000764793 |
| 1226 | Q9NQS1 | AVEN | Cell death regulator Aven | kptspLQSAGDHLEEEldlll | 275 | 285 | 130.5 | 0.000764793 |
| 1227 | Q7Z5N4 | SDK1 | Protein sidekick-1 | ltasqLEVTWDPPPPEsqngn | 1718 | 1728 | 130.5 | 0.000765265 |

| | | | | | | | | |
|---|---|---|---|---|---|---|---|---|
| 1228 | Q9Y2Y8 | PRG3 | Proteoglycan 3 | gtvsaLHLENDAPHLEsletq | 18 | 28 | 130.5 | 0.000765265 |
| 1229 | Q9P2G1 | ANKIB1 | Ankyrin repeat and IBR domain-containing protein 1 | lrpqdLRRLKDMLIVEtadml | 233 | 243 | 130.5 | 0.000766114 |
| 1230 | Q7Z2Z1 | TICRR | Treslin | ltaeeLHLVADVDPGEgrppi | 390 | 400 | 130.5 | 0.00076668 |
| 1231 | Q2NKX9 | C2orf68 | UPF0561 protein C2orf68 | hqlfcLEYEADSGEVTsvivy | 110 | 120 | 130.5 | 0.000767718 |
| 1232 | Q96AQ6 | PBXIP1 | Pre-B-cell leukemia transcription factor-interacting protein 1 | tdgkeLAGTMDGEGTLfqtes | 52 | 62 | 130.5 | 0.000768096 |
| 1233 | Q96HL8 | SH3YL1 | SH3 domain-containing YSC84-like protein 1 | aqaedLYEILDSFTEKyeneg | 203 | 213 | 130.5 | 0.000769134 |
| 1234 | Q9H9R9 | DBNDD1 | Dysbindin domain-containing protein 1 | fdlldLTELTDMSDQElaevf | 77 | 87 | 130.5 | 0.000771587 |
| 1235 | Q0VF96 | CGNL1 | Cingulin-like protein 1 | rkvkeLVMQVDDEHLSltdqk | 1186 | 1196 | 130.5 | 0.000772719 |
| 1237 | P28370 | SMARCA1 | Probable global transcription activator SNF2L1 | dekqsLISAGDYRHRRteqee | 138 | 148 | 130.5 | 0.000773757 |
| 1236 | Q15915 | ZIC1 | Zinc finger protein ZIC 1 | ngqmrLGFSGDMYPRPeqygq | 165 | 175 | 130.5 | 0.000773757 |
| 1238 | Q8TCZ2 | CD99L2 | CD99 antigen-like protein 2 | gsgldLADALDDQDDGrrkpg | 73 | 83 | 130.5 | 0.000773757 |
| 1239 | Q13029 | PRDM2 | PR domain zinc finger protein 2 | nisenLNYYIDGKIQTnnnts | 566 | 576 | 130.5 | 0.000775078 |
| 1240 | Q8NA77 | TEX19 | Testis-expressed protein 19 | ldfkdLLESEDWEEDNwdpel | 48 | 58 | 130.5 | 0.000775078 |
| 1241 | O94941 | UBOX5 | RING finger protein 37 | lqapaLPMESDCDPGDqpesq | 228 | 238 | 130.5 | 0.00077706 |
| 1242 | Q9P281 | BAHCC1 | BAH and coiled-coil domain-containing protein 1 | relarLQRKHDHERDEssrsp | 1463 | 1473 | 130.5 | 0.000777909 |
| 1243 | Q6ZMK1 | CYHR1 | Cysteine and histidine-rich protein 1 | ktgseLMEILDGMDQShrkem | 199 | 209 | 130.5 | 0.000778192 |
| 1244 | Q9ULT8 | HECTD1 | E3 ubiquitin-protein ligase HECTD1 | crastLLAELDDDEDLpepde | 1671 | 1681 | 130.5 | 0.000779042 |
| 1245 | O43424 | GRID2 | Glutamate receptor ionotropic, delta-2 | rrvnsLCTDDDSPHKQfstss | 884 | 894 | 130.5 | 0.000779513 |
| 1246 | O95153 | TSPOAP1 | Peripheral-type benzodiazepine receptor-associated protein 1 | gpldlLTSALDCGSLGdcppp | 535 | 545 | 130.5 | 0.00078074 |
| 1247 | Q9NP59 | SLC40A1 | Solute carrier family 40 member 1 | legthLMGVKDSNIHEleheq | 265 | 275 | 130.5 | 0.000782439 |
| 1248 | Q9BY43 | CHMP4A | Charged multivesicular body protein 4a | aisrpMGFGDDVDEDEllee l | 150 | 160 | 130.5 | 0.000782533 |
| 1249 | A0A1B0GWK | PVALEF | Parvalbumin-like EF-hand-containing protein | eeaeaMIQAADTHGDGrinye | 104 | 114 | 130.4 | 0.000783665 |
| 1250 | Q96PV0 | SYNGAP1 | Ras/Rap GTPase-activating protein SynGAP | fmargLNSSMDMARLPsptke | 777 | 787 | 130.4 | 0.000783665 |
| 1251 | Q15911 | ZFHX3 | Zinc finger homeobox protein 3 | ihpqfLDRSLDMPFMLfdpsn | 2566 | 2576 | 130.4 | 0.000784137 |
| 1252 | Q9UH77 | KLHL3 | Kelch-like protein 3 | lssqtLIQAGDDEKNQrtitv | 13 | 23 | 130.4 | 0.000784326 |
| 1253 | Q02952 | AKAP12 | A-kinase anchor protein 12 | pffegLEGSIDTGITVsrekv | 1284 | 1294 | 130.4 | 0.000784798 |
| 1254 | O00750 | PIK3C2B | Phosphatidylinositol 4-phosphate 3-kinase C2 domain-containing subunit bet | itrlnLKSTYDAEMLRdatrg | 240 | 250 | 130.4 | 0.000784892 |
| 1255 | O60299 | LZTS3 | Leucine zipper putative tumor suppressor 3 | akletLPVRADPGRDPllafa | 7 | 17 | 130.4 | 0.000785836 |
| 1257 | O60716 | CTNND1 | Catenin delta-1 | dhnrtLDRSGDLGDMEplkgt | 917 | 927 | 130.4 | 0.000786213 |
| 1256 | Q96JN0 | LCOR | Ligand-dependent corepressor | vlsklLMADQDSPLDLtvrks | 57 | 67 | 130.4 | 0.000786213 |
| 1258 | Q13563 | PKD2 | Polycystin-2 | geeggMVVEMDVEWRPgsrrs | 110 | 120 | 130.4 | 0.000786968 |
| 1259 | Q02241 | KIF23 | Kinesin-like protein KIF23 | pegyrLNRNGDYKETQysfkq | 62 | 72 | 130.4 | 0.000788195 |
| 1261 | Q09472 | EP300 | Histone acetyltransferase p300 | stelgLTNGGDINQLQtslgm | 55 | 65 | 130.4 | 0.000788289 |
| 1260 | Q12923 | PTPN13 | Tyrosine-protein phosphatase non-receptor type 13 | kpgdrLIKVNDTDVTNmthtd | 1834 | 1844 | 130.4 | 0.000788289 |
| 1262 | Q9P2P6 | STARD9 | StAR-related lipid transfer protein 9 | hrsfsLDSLIDAEEELgedqq | 1204 | 1214 | 130.4 | 0.000788761 |
| 1263 | Q8IUM7 | NPAS4 | Neuronal PAS domain-containing protein 4 | qqisqLAQGMDRPFSAeagtg | 642 | 652 | 130.4 | 0.000789421 |
| 1264 | Q96SN8 | CDK5RAP2 | CDK5 regulatory subunit-associated protein 2 | sqgqrLLAEMDIQTQEapsst | 1747 | 1757 | 130.4 | 0.000791025 |
| 1265 | O94842 | TOX4 | TOX high mobility group box family member 4 | eylkaLAAYKDNQECQatvet | 287 | 297 | 130.4 | 0.00079112 |
| 1266 | Q2M3G4 | SHROOM1 | Protein Shroom1 | ererlLQDQLDAIRDDlghha | 813 | 823 | 130.4 | 0.000792535 |
| 1267 | P37108 | SRP14 | Signal recognition particle 14 kDa protein | aysnlLRANMDGLKKRdkknk | 87 | 97 | 130.4 | 0.000792724 |
| 1269 | P43034 | PAFAH1B1 | Platelet-activating factor acetylhydrolase IB subunit beta | dfertLKGHTDSVQDIsfdhs | 146 | 156 | 130.4 | 0.000792913 |
| 1268 | Q92800 | EZH1 | Histone-lysine N-methyltransferase EZH1 | elvdaLNQYSDEEEEGhndts | 183 | 193 | 130.4 | 0.000792913 |
| 1271 | Q8NDZ2 | SIMC1 | SUMO-interacting motif-containing protein 1 | lpgdvLHSPGDMPHSSgdvth | 324 | 334 | 130.4 | 0.000793667 |
| 1270 | Q92576 | PHF3 | PHD finger protein 3 | pvgspLFKFSDKEEHEqndsi | 285 | 295 | 130.4 | 0.000793667 |
| 1272 | Q2UY09 | COL28A1 | Collagen alpha-1(XXVIII) chain | pgpmgLRGVGDTGAKGepgvr | 657 | 667 | 130.4 | 0.000794045 |
| 1273 | Q8NEN0 | ARMC2 | Armadillo repeat-containing protein 2 | slpshLKNGGDQGKRHarass | 220 | 230 | 130.4 | 0.000794045 |
| 1274 | Q15699 | ALX1 | ALX homeobox protein 1 | plhteLNRAMDNCNSLrmspv | 92 | 102 | 130.4 | 0.000794989 |
| 1275 | Q8IVF2 | AHNAK2 | Protein AHNAK2 | pastdLKVQADQVDVKlpegh | 1055 | 1065 | 130.4 | 0.000794989 |
| 1276 | Q6ZU80 | CEP128 | Centrosomal protein of 128 kDa | pptspLKDYGDPQGIKrmrsr | 131 | 141 | 130.4 | 0.000795272 |
| 1277 | Q63HK5 | TSHZ3 | Teashirt homolog 3 | lrenaLSDISDMLKNLtesht | 844 | 854 | 130.4 | 0.00079546 |
| 1278 | Q86UP3 | ZFHX4 | Zinc finger homeobox protein 4 | tvaasLKRKLDDKEDNncsek | 2533 | 2543 | 130.4 | 0.000795932 |
| 1279 | Q02779 | MAP3K10 | Mitogen-activated protein kinase kinase kinase 10 | atlvsLSSVSDCNSTRsllrs | 749 | 759 | 130.4 | 0.000796781 |
| 1282 | A1L4H1 | SSC5D | Soluble scavenger receptor cysteine-rich domain-containing protein SSC5D | kdgykLPWTWDTPSGRglaeg | 936 | 946 | 130.4 | 0.000797159 |
| 1281 | Q6ZN55 | ZNF574 | Zinc finger protein 574 | qqhfeLVGVADPGVTVatdta | 46 | 56 | 130.4 | 0.000797159 |
| 1280 | Q9UKS7 | IKZF2 | Zinc finger protein Helios | tnsvkLEMQSDEECDRkplsr | 52 | 62 | 130.4 | 0.000797159 |
| 1283 | Q3YEC7 | RABL6 | Rab-like protein 6 | krgdgLKMENDPQEAEsemal | 114 | 124 | 130.4 | 0.00079914 |
| 1284 | Q92752 | TNR | Tenascin-R | grqqsLESTVDAFTGFrpish | 764 | 774 | 130.4 | 0.000799235 |
| 1285 | O15061 | SYNM | Synemin | srtfvLAGSADSPELGklads | 1429 | 1439 | 130.4 | 0.000800178 |
| 1286 | Q8ND04 | SMG8 | Protein SMG8 | rqvfpLFRHQDPGDPGpgirt | 79 | 89 | 130.3 | 0.00080216 |
| 1287 | Q8TDR2 | STK35 | Serine/threonine-protein kinase 35 | pdafeLETRMDQVTCAa | 523 | 533 | 130.3 | 0.000803292 |
| 1288 | Q16543 | CDC37 | Hsp90 co-chaperone Cdc37 | dakyhMQRCIDSGLWVpnska | 333 | 343 | 130.3 | 0.000803387 |
| 1289 | Q9H582 | ZNF644 | Zinc finger protein 644 | nvlngLANNMDDLKINtditg | 22 | 32 | 130.3 | 0.000803953 |
| 1290 | O15056 | SYNJ2 | Synaptojanin-2 | ldspqLSGATDSQDDSspadi | 565 | 575 | 130.3 | 0.000804142 |
| 1291 | P0CE72 | OCM | Oncomodulin-1 | diaaaLQECRDPDTFEpqkff | 16 | 26 | 130.3 | 0.000804142 |
| 1292 | P19838 | NFKB1 | Nuclear factor NF-kappa-B p105 subunit | enfepLYDLDDSWENAgedeg | 753 | 763 | 130.3 | 0.000804142 |
| 1293 | P23443 | RPS6KB1 | Ribosomal protein S6 kinase beta-1 | reaedMAGVFDIDLDQpedag | 24 | 34 | 130.3 | 0.000805179 |
| 1294 | Q13489 | BIRC3 | Baculoviral IAP repeat-containing protein 3 | hlleqLLSTSDSPGDEnaess | 345 | 355 | 130.3 | 0.00080584 |
| 1295 | Q9NYI0 | PSD3 | PH and SEC7 domain-containing protein 3 | qgveiLWTGGDKRETQhpidf | 307 | 317 | 130.3 | 0.00080584 |
| 1296 | Q9ULM3 | YEATS2 | YEATS domain-containing protein 2 | sesdsLSQHNDFLSDKdnnsn | 143 | 153 | 130.3 | 0.00080584 |
| 1299 | Q460N5 | PARP14 | Protein mono-ADP-ribosyltransferase PARP14 | dtklpLDGGLDKMEDIpeece | 130 | 140 | 130.3 | 0.000806689 |
| 1297 | Q5T5U3 | ARHGAP21 | Rho GTPase-activating protein 21 | lrghsLYLYKDKREQTtpsee | 970 | 980 | 130.3 | 0.000806689 |
| 1298 | Q9Y6I3 | EPN1 | Epsin-1 | keessLMDLADVFTAPapapt | 257 | 267 | 130.3 | 0.000806689 |
| 1300 | Q9NVP4 | DZANK1 | Double zinc ribbon and ankyrin repeat-containing protein 1 | dnntlLEMKSDTPDVNiyytl | 32 | 42 | 130.3 | 0.000807255 |
| 1302 | P13984 | GTF2F2 | General transcription factor IIF subunit 2 | tlnedLANIHDIGGKPasvsa | 62 | 72 | 130.3 | 0.000807916 |
| 1301 | Q9C0C7 | AMBRA1 | Activating molecule in BECN1-regulated autophagy protein 1 | achnlLTFNNDTLRWErttpn | 575 | 585 | 130.3 | 0.000807916 |
| 1303 | Q9P232 | CNTN3 | Contactin-3 | gsrseLVITWDPVPEElqnge | 717 | 727 | 130.3 | 0.000808388 |
| 1304 | O15068 | MCF2L | Guanine nucleotide exchange factor DBS | gdvveLVQEGDEGLWYvrdpt | 1084 | 1094 | 130.3 | 0.00080886 |
| 1305 | Q8TBY9 | CFAP251 | Cilia- and flagella-associated protein 251 | teilgLTISEDSGQDGq | 1138 | 1148 | 130.3 | 0.000809426 |
| 1306 | Q9Y6X3 | MAU2 | MAU2 chromatid cohesion factor homolog | lpehnLITWTDGPPPVqfqaq | 587 | 597 | 130.3 | 0.000810747 |
| 1307 | Q8NCE0 | TSEN2 | tRNA-splicing endonuclease subunit Sen2 | sqhigLLHPGDRGPDHeyvlv | 245 | 255 | 130.3 | 0.000811973 |
| 1308 | Q3V6T2 | CCDC88A | Girdin | ktveeLRTTVDSVEGNaskil | 479 | 489 | 130.3 | 0.00081254 |
| 1309 | Q9UL36 | ZNF236 | Zinc finger protein 236 | apqdpLRGHVDQFEEQspaqq | 858 | 868 | 130.3 | 0.000812728 |
| 1311 | P10071 | GLI3 | Transcriptional activator GLI3 | vtlesLTMDADANLNDedflp | 1078 | 1088 | 130.3 | 0.000813106 |
| 1312 | P30304 | CDC25A | M-phase inducer phosphatase 1 | spvtnLTVTMDQLQGLgsdye | 45 | 55 | 130.3 | 0.000813106 |
| 1310 | Q15276 | RABEP1 | Rab GTPase-binding effector protein 1 | klqlmLRQANDQLEKTmkdkq | 564 | 574 | 130.3 | 0.000813106 |
| 1313 | Q08378 | GOLGA3 | Golgin subfamily A member 3 | qemenLKWEVDQKEREiqslk | 1289 | 1299 | 130.3 | 0.0008132 |
| 1314 | Q9H981 | ACTR8 | Actin-related protein 8 | aqgdgLMAGNDSEEALtalms | 481 | 491 | 130.3 | 0.000813672 |
| 1315 | Q8IV76 | PASD1 | Circadian clock protein PASD1 | nqqnaLELMMDHLQKQpntlr | 402 | 412 | 130.3 | 0.000815276 |

| | | | | | | | | |
|---|---|---|---|---|---|---|---|---|
| 1316 | Q9Y2H0 | DLGAP4 | Disks large-associated protein 4 | atcpsLGVGTDTNYVKrgsws | 280 | 290 | 130.3 | 0.000816503 |
| 1317 | Q01094 | E2F1 | Transcription factor E2F1 | vtcqdLRSIADPAEQMvmvik | 251 | 261 | 130.3 | 0.000817069 |
| 1318 | Q04864 | REL | Proto-oncogene c-Rel | mpsadLYGISDPNMLSncsvn | 482 | 492 | 130.3 | 0.000817918 |
| 1319 | Q5VZ46 | KIAA1614 | Uncharacterized protein KIAA1614 | aalagLLRLGDQTEPVgiprp | 830 | 840 | 130.3 | 0.000818296 |
| 1320 | Q9NQT8 | KIF13B | Kinesin-like protein KIF13B | qqlqkLVSKRDKTEDDadrea | 1113 | 1123 | 130.3 | 0.000818767 |
| 1321 | Q9UN79 | SOX13 | Transcription factor SOX-13 | leeamLSCDMDGSRHFpesrn | 405 | 415 | 130.3 | 0.00081905 |
| 1322 | Q9Y6R1 | SLC4A4 | Electrogenic sodium bicarbonate cotransporter 1 | ltsssLNDISDKPEKDqlknk | 258 | 268 | 130.2 | 0.000819711 |
| 1323 | Q14562 | DHX8 | ATP-dependent RNA helicase DHX8 | lerkrLTRISDPEKWEikqmi | 391 | 401 | 130.2 | 0.000820183 |
| 1324 | O94967 | WDR47 | WD repeat-containing protein 47 | qklgeLNIGMDGLGNEvsaln | 486 | 496 | 130.2 | 0.000820749 |
| 1325 | Q8NEV8 | EXPH5 | Exophilin-5 | spsepLNIYEDDPVDSncdtd | 1956 | 1966 | 130.2 | 0.000821409 |
| 1326 | P15924 | DSP | Desmoplakin | rseddLRQQRDVLDGHlrekq | 1633 | 1643 | 130.2 | 0.000821693 |
| 1327 | Q9Y4J8 | DTNA | Dystrobrevin alpha | ekplnLAHIVDTWPPRpvtsm | 330 | 340 | 130.2 | 0.000821693 |
| 1328 | A6NCF5 | KLHL33 | Kelch-like protein 33 | dllhqLMVEADVPGQErrrep | 192 | 202 | 130.2 | 0.000822164 |
| 1329 | Q6ZS94 | C1orf229 | Putative uncharacterized protein C1orf229 | hlprrLRLAFDFLEPRargqr | 37 | 47 | 130.2 | 0.000822164 |
| 1330 | Q9Y5H0 | PCDHGA3 | Protocadherin gamma-A3 | itqdlLEMKGDSNLLQqappn | 798 | 808 | 130.2 | 0.000822353 |
| 1331 | Q9NRS6 | SNX15 | Sorting nexin-15 | ayaaaLQGYRDGVHVLlqgvp | 292 | 302 | 130.2 | 0.000822636 |
| 1332 | Q9Y2Z0 | SUGT1 | Protein SGT1 homolog | vrwekLEGQGDVPTPKqfvad | 257 | 267 | 130.2 | 0.000823863 |
| 1333 | Q86W92 | PPFIBP1 | Liprin-beta-1 | nnlthMLKEDDMFKDFaarsp | 985 | 995 | 130.2 | 0.000824146 |
| 1334 | O60240 | PLIN1 | Perilipin-1 | slsdaLKGVTDNVVDTvvhyv | 387 | 397 | 130.2 | 0.00082424 |
| 1335 | Q6KC79 | NIPBL | Nipped-B-like protein | sieniLDNLEDMDFTAfgddd | 1210 | 1220 | 130.2 | 0.000824429 |
| 1336 | A0A1B0GTU | ZC3H11B | Zinc finger CCCH domain-containing protein 11B | slkerLGMSADPNNEDatdkv | 335 | 345 | 130.2 | 0.000824523 |
| 1337 | Q9H5V7 | IKZF5 | Zinc finger protein Pegasus | kkpepLDFVKDFQEYLtqqth | 9 | 19 | 130.2 | 0.000825939 |
| 1338 | P27482 | CALML3 | Calmodulin-like protein 3 | eevdeMIRAADTDGDGqvnye | 125 | 135 | 130.2 | 0.000826128 |
| 1339 | Q00994 | BEX3 | Protein BEX3 | ilmgeLSNHHDHHDEFclmp | 97 | 107 | 130.2 | 0.000826694 |
| 1340 | Q9H6J7 | C11orf49 | UPF0705 protein C11orf49 | qalgaLPDKGDLMHDPamdee | 255 | 265 | 130.2 | 0.000827732 |
| 1341 | Q9Y210 | TRPC6 | Short transient receptor potential channel 6 | ikryvLQAQIDKESDEvnege | 868 | 878 | 130.2 | 0.000829053 |
| 1343 | Q07075 | ENPEP | Glutamyl aminopeptidase | glavgLTRSCDSSGDGgpgta | 39 | 49 | 130.2 | 0.000829996 |
| 1342 | Q8NAP3 | ZBTB38 | Zinc finger and BTB domain-containing protein 38 | wgeeaLKMDLDNNFYStevsv | 660 | 670 | 130.2 | 0.000829996 |
| 1344 | Q9H7C4 | SYNC | Syncoilin | edilyLEDTGDLDETLyvqet | 61 | 71 | 130.2 | 0.000830468 |
| 1345 | Q9BSK4 | FEM1A | Protein fem-1 homolog A | ngftpLHMAVDKDTTNvgryp | 539 | 549 | 130.2 | 0.000830846 |
| 1346 | Q8N393 | ZNF786 | Zinc finger protein 786 | prnldLPGLWDVPAWEstqhp | 173 | 183 | 130.2 | 0.000833676 |
| 1347 | O00555 | CACNA1A | Voltage-dependent P/Q-type calcium channel subunit alpha-1A | asersLGRYTDVDTGLgtdls | 2167 | 2177 | 130.2 | 0.000834054 |
| 1348 | O15060 | ZBTB39 | Zinc finger and BTB domain-containing protein 39 | mdelqLEDLGDDDLQFedpae | 296 | 306 | 130.2 | 0.000834054 |
| 1349 | O95983 | MBD3 | Methyl-CpG-binding domain protein 3 | diaeeLVKTMDLPKGLqgvgp | 155 | 165 | 130.2 | 0.000834054 |
| 1350 | P14652 | HOXB2 | Homeobox protein Hox-B2 | acpgaLEDICDPAEEPaaspg | 218 | 228 | 130.2 | 0.000834431 |
| 1351 | Q5JSZ5 | PRRC2B | Protein PRRC2B | kglddLDADADDGWAGlheev | 364 | 374 | 130.2 | 0.000834714 |
| 1353 | A4QPH2 | PI4KAP2 | Putative phosphatidylinositol 4-kinase alpha-like protein P2 | lekegLRCRSDSEDECstqea | 289 | 299 | 130.2 | 0.000835469 |
| 1352 | P42356 | PI4KA | Phosphatidylinositol 4-kinase alpha | lekegLRCRSDSEDECstqea | 1822 | 1832 | 130.2 | 0.000835469 |
| 1354 | P19013 | KRT4 | Keratin, type II cytoskeletal 4 | tkvqqLQISVDQHGDNlkntk | 337 | 347 | 130.2 | 0.000835564 |
| 1355 | Q86YH2 | ZNF280B | Zinc finger protein 280B | elpspLITFTDSLHHPvstal | 145 | 155 | 130.2 | 0.000835658 |
| 1356 | Q9H3M7 | TXNIP | Thioredoxin-interacting protein | spttpLLDDMDGSQDSpifmy | 351 | 361 | 130.2 | 0.000836507 |
| 1357 | Q5VTJ3 | KLHDC7A | Kelch domain-containing protein 7A | hyvapLQGSSDMNQSWvftrv | 213 | 223 | 130.2 | 0.000838394 |
| 1358 | Q8N8E3 | CEP112 | Centrosomal protein of 112 kDa | qnmklLQTKYDADINLlkqeh | 480 | 490 | 130.1 | 0.000843207 |
| 1359 | Q04637 | EIF4G1 | Eukaryotic translation initiation factor 4 gamma 1 | pisrgLPLVDDGGWNTvpisk | 1044 | 1054 | 130.1 | 0.000844905 |
| 1361 | Q5JRA6 | MIA3 | Transport and Golgi organization protein 1 homolog | kgkdtLKSAYDDTENDlkgaa | 517 | 527 | 130.1 | 0.000845 |
| 1362 | Q6UVJ0 | SASS6 | Spindle assembly abnormal protein 6 homolog | glsqnLFSNSDHQRDGtlgal | 618 | 628 | 130.1 | 0.000845 |
| 1360 | Q92922 | SMARCC1 | SWI/SNF complex subunit SMARCC1 | rhfeeLETIMDREKEAleqqr | 917 | 927 | 130.1 | 0.000845 |
| 1363 | Q9Y6N7 | ROBO1 | Roundabout homolog 1 | yisgpLVSDMDTDAPEeeede | 1318 | 1328 | 130.1 | 0.000845 |
| 1364 | Q8TCG1 | CIP2A | Protein CIP2A | kknkdLQITCDSLNKQietvk | 748 | 758 | 130.1 | 0.000845849 |
| 1365 | Q99490 | AGAP2 | Arf-GAP with GTPase, ANK repeat and PH domain-containing protein 2 | krherLFHRQDALWIStssag | 69 | 79 | 130.1 | 0.000845849 |
| 1366 | O15350 | TP73 | Tumor protein p73 | vegnnLSQYVDDPVTGrqsvv | 222 | 232 | 130.1 | 0.000848208 |
| 1368 | P11277 | SPTB | Spectrin beta chain, erythrocytic | qdahrLLSGEDVGQDEgatra | 765 | 775 | 130.1 | 0.000849435 |
| 1367 | Q9NRA8 | EIF4ENIF1 | Eukaryotic translation initiation factor 4E transporter | tnnrqLKKDGDMTAFNklvst | 475 | 485 | 130.1 | 0.000849435 |
| 1369 | O75420 | GIGYF1 | GRB10-interacting GYF protein 1 | pspppLLGNMDQERLKkqqel | 542 | 552 | 130.1 | 0.000849529 |
| 1370 | Q5VST9 | OBSCN | Obscurin | stadeLARTGDADLSHtssdd | 4792 | 4802 | 130.1 | 0.000849623 |
| 1371 | Q71F56 | MED13L | Mediator of RNA polymerase II transcription subunit 13-like | svaeeLCMEQDTPGQKlglag | 497 | 507 | 130.1 | 0.000849812 |
| 1372 | P08048 | ZFY | Zinc finger Y-chromosomal protein | pgeddLGGTVDIVESEpendh | 261 | 271 | 130.1 | 0.000850095 |
| 1373 | P17010 | ZFX | Zinc finger X-chromosomal protein | pgeddLGGTVDIVESEpendh | 265 | 275 | 130.1 | 0.000850095 |
| 1374 | Q7Z2Y5 | NRK | Nik-related protein kinase | ngnddLDNQVDQANDVckdhd | 955 | 965 | 130.1 | 0.000850473 |
| 1375 | P30307 | CDC25C | M-phase inducer phosphatase 3 | itatqLTTSADLDETGhldss | 87 | 97 | 130.1 | 0.000851039 |
| 1377 | D6RIA3 | C4orf54 | Uncharacterized protein C4orf54 | dfdvgLASRWDFEDNNviysf | 388 | 398 | 130.1 | 0.000851133 |
| 1376 | Q8IYH5 | ZZZ3 | ZZ-type zinc finger-containing protein 3 | gfvekLQKKADIGLPYpqrvv | 545 | 555 | 130.1 | 0.000851133 |
| 1378 | Q13253 | NOG | Noggin | lplvdLIEHPDPIFDPkekdl | 46 | 56 | 130.1 | 0.000852832 |
| 1379 | Q5T036 | FAM120AOS | Uncharacterized protein FAM120AOS | MGKTKDIGDDDtvase | 1 | 11 | 130.1 | 0.000854436 |
| 1381 | Q92889 | ERCC4 | DNA repair endonuclease XPF | gkpeeLEEEGDVEEGYrreis | 504 | 514 | 130.1 | 0.000855662 |
| 1380 | Q9UN81 | L1RE1 | LINE-1 retrotransposable element ORF1 protein | eecrsLRSRCDQLEERvsame | 107 | 117 | 130.1 | 0.000855662 |
| 1382 | Q9NYF8 | BCLAF1 | Bcl-2-associated transcription factor 1 | skkyfLHDDRDDGVDYwakrg | 848 | 858 | 130.1 | 0.000856323 |
| 1384 | Q96MA1 | DMRTB1 | Doublesex- and mab-3-related transcription factor B1 | pssfsLTVLFDTDKENtddqd | 309 | 319 | 130.1 | 0.000857078 |
| 1383 | Q9Y608 | LRRFIP2 | Leucine-rich repeat flightless-interacting protein 2 | tslseLRDIYDLKDQlqdveg | 344 | 354 | 130.1 | 0.000857078 |
| 1385 | Q5T1B0 | AXDND1 | Axonemal dynein light chain domain-containing protein 1 | tpkgtLPRLVDHVWHHpvrrn | 93 | 103 | 130.1 | 0.00085821 |
| 1386 | P13611 | VCAN | Versican core protein | ptvapLPFSTDIGHPQnqtvr | 2349 | 2359 | 130.1 | 0.000858682 |
| 1387 | A5D8V7 | ODAD3 | Outer dynein arm-docking complex subunit 3 | hrehlLLQSDDTIQDSlhake | 328 | 338 | 130.0 | 0.00086123 |
| 1388 | Q8WWL2 | SPIRE2 | Protein spire homolog 2 | erlidLMANNDSEDSGcgaad | 131 | 141 | 130.0 | 0.000861324 |
| 1389 | A1L4H1 | SSC5D | Soluble scavenger receptor cysteine-rich domain-containing protein SSC5D | gspptLRVHGDTGSPRkpwpe | 976 | 986 | 130.0 | 0.000861513 |
| 1390 | Q5EBL4 | RILPL1 | RILP-like protein 1 | qvmkkLKEVVDKQRDEirakd | 166 | 176 | 130.0 | 0.000862362 |
| 1391 | Q9UIF8 | BAZ2B | Bromodomain adjacent to zinc finger domain protein 2B | ksftkLCKEHDGEFTGedess | 1843 | 1853 | 130.0 | 0.0008634 |
| 1392 | Q5VUA4 | ZNF318 | Zinc finger protein 318 | kesqeLRKSEDCRESEietnt | 1764 | 1774 | 130.0 | 0.000866986 |
| 1393 | P35712 | SOX6 | Transcription factor SOX-6 | ngedeMEMYDDYEDDPksdys | 801 | 811 | 130.0 | 0.000867552 |
| 1394 | Q9BXF3 | CECR2 | Cat eye syndrome critical region protein 2 | hqprtLGHVMDSRVMRppvpp | 825 | 835 | 130.0 | 0.000867835 |
| 1395 | Q14008 | CKAP5 | Cytoskeleton-associated protein 5 | dekpaLLSQIDAEFEKmqgqs | 801 | 811 | 130.0 | 0.000868212 |
| 1396 | Q49A88 | CCDC14 | Coiled-coil domain-containing protein 14 | dapekLSRASDMKDTQllkki | 845 | 855 | 130.0 | 0.000868779 |
| 1398 | P48960 | ADGRE5 | Adhesion G protein-coupled receptor E5 | vhsqtLSRFFDKVQDLgrdsk | 275 | 285 | 130.0 | 0.000869722 |
| 1397 | Q496M5 | PLK5 | Inactive serine/threonine-protein kinase PLK5 | aalrhLQLCLDVGPPAtqdpl | 180 | 190 | 130.0 | 0.000869722 |
| 1399 | Q9UHX3 | ADGRE2 | Adhesion G protein-coupled receptor E2 | vhsqtLSRFFDKVQDLgrdyk | 278 | 288 | 130.0 | 0.000869722 |
| 1400 | P0C860 | MSL3P1 | Putative male-specific lethal-3 protein-like 2 | ndensLSSSSDSSEDKdekis | 54 | 64 | 130.0 | 0.0008701 |
| 1401 | Q92918 | MAP4K1 | Mitogen-activated protein kinase kinase kinase kinase 1 | sprkqLSESSDDDYDDvdipt | 373 | 383 | 130.0 | 0.000870666 |
| 1402 | A8MUI8 | | Putative UPF0607 protein ENSP00000383783 | qptppLQLKWDRDEGPppakf | 265 | 275 | 130.0 | 0.000871798 |
| 1403 | Q02224 | CENPE | Centromere-associated protein E | emelkLKEKNDLDEFEalerk | 528 | 538 | 130.0 | 0.000871892 |

| | | | | | | | | |
|---|---|---|---|---|---|---|---|---|
| 1404 | Q3KR37 | GRAMD1B | Protein Aster-B | cygneLGLTSDDEDYVppddd | 226 | 236 | 130.0 | 0.000871987 |
| 1406 | P23743 | DGKA | Diacylglycerol kinase alpha | httktLPMQIDGEPWMqtpct | 693 | 703 | 130.0 | 0.00087227 |
| 1405 | Q9Y6T7 | DGKB | Diacylglycerol kinase beta | rtsksLPMQIDGEPWMqtpct | 754 | 764 | 130.0 | 0.00087227 |
| 1407 | Q07617 | SPAG1 | Sperm-associated antigen 1 | eatedLSKVLDVEPDNdlakk | 299 | 309 | 130.0 | 0.000872836 |
| 1408 | Q13351 | KLF1 | Krueppel-like factor 1 | stltaLGPFPDTQDDFlkwwr | 17 | 27 | 130.0 | 0.000873213 |
| 1409 | Q14693 | LPIN1 | Phosphatidate phosphatase LPIN1 | adgvyLDDLTDMDPEVaalyf | 399 | 409 | 130.0 | 0.000874535 |
| 1410 | Q8WYA6 | CTNNBL1 | Beta-catenin-like protein 1 | ddkkrLLQIIDRDGEEeeeee | 59 | 69 | 130.0 | 0.000874535 |
| 1411 | Q8TB72 | PUM2 | Pumilio homolog 2 | ingrgLPNGMDADCKDfnrtp | 161 | 171 | 130.0 | 0.00087595 |
| 1412 | Q6ULP2 | AFTPH | Aftiphilin | issemLATSIDGMERPgnlnk | 121 | 131 | 130.0 | 0.000876422 |
| 1413 | Q6UX27 | VSTM1 | V-set and transmembrane domain-containing protein 1 | nvtfvLRKVNDSGYKQeqssa | 60 | 70 | 130.0 | 0.000876988 |
| 1414 | P55291 | CDH15 | Cadherin-15 | tlssiLSSQGDEDQDYdylrd | 753 | 763 | 130.0 | 0.000877554 |
| 1415 | O14662 | STX16 | Syntaxin-16 | dtsvpLMDDGDDNTLYhrgft | 204 | 214 | 130.0 | 0.000878215 |
| 1416 | Q92544 | TM9SF4 | Transmembrane 9 superfamily member 4 | atrleLYSNRDSDDKKkekdv | 146 | 156 | 130.0 | 0.000878215 |
| 1417 | A6NJL1 | ZSCAN5B | Zinc finger and SCAN domain-containing protein 5B | lgkeyLMLNSDVEMAEapasv | 139 | 149 | 130.0 | 0.00087963 |
| 1418 | Q5PRF9 | SAMD4B | Protein Smaug homolog 2 | spqaiLMFPPDCPVPGpdlei | 653 | 663 | 129.9 | 0.000883027 |
| 1419 | Q8WXI7 | MUC16 | Mucin-16 | erlttLEDTVDTEDMQpstht | 8336 | 8346 | 129.9 | 0.00088331 |
| 1420 | Q8IVT2 | MISP | Mitotic interactor and substrate of PLK1 | reqrgLRQATDHQELVeiptr | 313 | 323 | 129.9 | 0.000883688 |
| 1421 | Q8WU90 | ZC3H15 | Zinc finger CCCH domain-containing protein 15 | didlsLYIPRDVDETGitvas | 336 | 346 | 129.9 | 0.00088548 |
| 1422 | Q08999 | RBL2 | Retinoblastoma-like protein 2 | nhsalLRRLQDVANDRgsh | 1126 | 1136 | 129.9 | 0.000885669 |
| 1423 | P25440 | BRD2 | Bromodomain-containing protein 2 | asalgLHDYHDIIKHPmdlst | 383 | 393 | 129.9 | 0.000887368 |
| 1424 | Q15059 | BRD3 | Bromodomain-containing protein 3 | aealeLHDYHDIIKHPmdlst | 345 | 355 | 129.9 | 0.000887368 |
| 1425 | Q53LP3 | SOWAHC | Ankyrin repeat domain-containing protein SOWAHC | klshaLEDGGDHHHHHhsaeg | 432 | 442 | 129.9 | 0.0008885 |
| 1427 | P15170 | GSPT1 | Eukaryotic peptide chain release factor GTP-binding subunit ERF3A | retwyLSWALDTNQEErdkgk | 123 | 133 | 129.9 | 0.000888783 |
| 1426 | Q8IYD1 | GSPT2 | Eukaryotic peptide chain release factor GTP-binding subunit ERF3B | retwyLSWALDTNQEErdkgk | 252 | 262 | 129.9 | 0.000888783 |
| 1428 | P13611 | VCAN | Versican core protein | kevgpLVSQTDIFEGSgsvts | 2314 | 2324 | 129.9 | 0.000890198 |
| 1429 | P46939 | UTRN | Utrophin | sdseeLTQRWDSLVQRledss | 617 | 627 | 129.9 | 0.000890198 |
| 1430 | Q6UXC1 | MAMDC4 | Apical endosomal glycoprotein | vepqqLCDGEDNCGDLsdenp | 247 | 257 | 129.9 | 0.000890953 |
| 1431 | Q7Z7A1 | CNTRL | Centriolin | keikdLQIAIDSLDSKdpkhs | 536 | 546 | 129.9 | 0.000890953 |
| 1432 | Q86T90 | KIAA1328 | Protein hinderin | ismesLKGTGDSVDEQnscrg | 69 | 79 | 129.9 | 0.000891614 |
| 1433 | D6RIA3 | C4orf54 | Uncharacterized protein C4orf54 | rksdpLPMMMDSHVLSliase | 1282 | 1292 | 129.9 | 0.000891803 |
| 1434 | Q9UBF1 | MAGEC2 | Melanoma-associated antigen C2 | ptsveLEDWVDAQHPTdeeee | 22 | 32 | 129.9 | 0.000892086 |
| 1435 | Q8IXR5 | FAM178B | Protein FAM178B | tvpilLYNLEDGLSDHpldqg | 59 | 69 | 129.9 | 0.000892652 |
| 1436 | Q9H0W5 | CCDC8 | Coiled-coil domain-containing protein 8 | spgdrLGNAGDVCVPQasprr | 249 | 259 | 129.9 | 0.000892935 |
| 1437 | O60282 | KIF5C | Kinesin heavy chain isoform 5C | kekehLTRLQDAEEMKkaleq | 690 | 700 | 129.9 | 0.000893124 |
| 1438 | Q6ZUT6 | CCDC9B | Coiled-coil domain-containing protein 9B | idlarLARHRDAQGDWrrpwd | 250 | 260 | 129.9 | 0.000893501 |
| 1439 | Q92620 | DHX38 | Pre-mRNA-splicing factor ATP-dependent RNA helicase PRP16 | gvvhrLEVDEDFEEDNaakvh | 391 | 401 | 129.9 | 0.000893501 |
| 1440 | Q5THJ4 | VPS13D | Vacuolar protein sorting-associated protein 13D | tlsdgLGKTMDNRHQSereyi | 4120 | 4130 | 129.9 | 0.000893878 |
| 1441 | Q8N4C6 | NIN | Ninein | qhearLKELFDSFDTTgtgsl | 13 | 23 | 129.9 | 0.000894067 |
| 1442 | P53667 | LIMK1 | LIM domain kinase 1 | hgkrgLSVSIDPPHGPpgcgt | 178 | 188 | 129.9 | 0.000895105 |
| 1443 | Q96G25 | MED8 | Mediator of RNA polymerase II transcription subunit 8 | qviipLVLSPDRDEDLmrqte | 79 | 89 | 129.9 | 0.0008952 |
| 1444 | Q8IWJ2 | GCC2 | GRIP and coiled-coil domain-containing protein 2 | qerehLEMLIDQLKIKlqdsq | 1354 | 1364 | 129.9 | 0.00089586 |
| 1445 | Q7Z5L2 | R3HCC1L | Coiled-coil domain-containing protein R3HCC1L | avgidLGSTGDTTEALhelrt | 509 | 519 | 129.9 | 0.000896804 |
| 1446 | Q9H094 | NBPF3 | Neuroblastoma breakpoint family member 3 | hlvqkLSPENDDDEDEdvkve | 218 | 228 | 129.9 | 0.000896898 |
| 1448 | Q8IWZ3 | ANKHD1 | Ankyrin repeat and KH domain-containing protein 1 | qvdtiLFKDNDVDDEQqspps | 903 | 913 | 129.9 | 0.000897087 |
| 1447 | Q8WXI7 | MUC16 | Mucin-16 | vltsgLVKTTDMLNTSmepvt | 4490 | 4500 | 129.9 | 0.000897087 |
| 1450 | P11532 | DMD | Dystrophin | arekeLQTIFDTLPPMryqet | 932 | 942 | 129.9 | 0.000897653 |
| 1449 | Q9BWV1 | BOC | Brother of CDO | sthqlLQPHHDCCQRQeqpaa | 1012 | 1022 | 129.9 | 0.000897653 |
| 1451 | Q53GI3 | ZNF394 | Zinc finger protein 394 | peaegLNSISDVNKNGsiege | 264 | 274 | 129.9 | 0.00089803 |
| 1452 | Q86VF7 | NRAP | Nebulin-related-anchoring protein | rsrgkLIGAKDVQGDSqmshs | 833 | 843 | 129.9 | 0.00089803 |
| 1453 | Q92485 | SMPDL3B | Acid sphingomyelinase-like phosphodiesterase 3b | epdfiLWTGDDTPHVPdeklg | 89 | 99 | 129.9 | 0.000898408 |
| 1454 | Q53QZ3 | ARHGAP15 | Rho GTPase-activating protein 15 | sssteLLSHYDSDIKEqkpeh | 215 | 225 | 129.9 | 0.000898502 |
| 1455 | Q8NI08 | NCOA7 | Nuclear receptor coactivator 7 | tlsssLSQAGDPITEGnkepd | 564 | 574 | 129.9 | 0.000899446 |
| 1456 | Q14934 | NFATC4 | Nuclear factor of activated T-cells, cytoplasmic 4 | sdeaaLYAACDEVESElneaa | 191 | 201 | 129.9 | 0.000899729 |
| 1457 | O75112 | LDB3 | LIM domain-binding protein 3 | lsqgdLVVAIDGVNTDtmthl | 48 | 58 | 129.9 | 0.000901144 |
| 1459 | P98082 | DAB2 | Disabled homolog 2 | dqtnkLKSGVDQMDLFgdmst | 206 | 216 | 129.9 | 0.000901144 |
| 1458 | Q99941 | ATF6B | Cyclic AMP-dependent transcription factor ATF-6 beta | slappLCLLGDDPTSSfetvq | 133 | 143 | 129.9 | 0.000901144 |
| 1460 | Q96JG9 | ZNF469 | Zinc finger protein 469 | vaghqLGLEADGHWGLlgqae | 1947 | 1957 | 129.8 | 0.000901616 |
| 1461 | Q99570 | PIK3R4 | Phosphoinositide 3-kinase regulatory subunit 4 | vqlknLNMENDPNNEEidevt | 505 | 515 | 129.8 | 0.000902277 |
| 1462 | Q5JSP0 | FGD3 | FYVE, RhoGEF and PH domain-containing protein 3 | gdpvsLAAAGDGSPDIgptge | 49 | 59 | 129.8 | 0.00090256 |
| 1463 | Q9H3P2 | NELFA | Negative elongation factor A | akgrgLLRKMDTTTPLkgipk | 204 | 214 | 129.8 | 0.000902937 |
| 1465 | Q6PJT7 | ZC3H14 | Zinc finger CCCH domain-containing protein 14 | vmaetLQMSQDYYDMEsmvha | 447 | 457 | 129.8 | 0.000904447 |
| 1464 | Q8IY92 | SLX4 | Structure-specific endonuclease subunit SLX4 | rkegsLPHSDDAGDYEqlfss | 974 | 984 | 129.8 | 0.000904447 |
| 1466 | Q14592 | ZNF460 | Zinc finger protein 460 | prysyLGQAMDQDGPSemqey | 90 | 100 | 129.8 | 0.000907938 |
| 1467 | O75923 | DYSF | Dysferlin | ptlpdLDVVADTGGEEdtedq | 148 | 158 | 129.8 | 0.000908316 |
| 1468 | Q9HCK8 | CHD8 | Chromodomain-helicase-DNA-binding protein 8 | aqftkLRRGMDEKEFTvqikd | 2242 | 2252 | 129.8 | 0.000909165 |
| 1470 | Q8IZ63 | PRR22 | Proline-rich protein 22 | pgpeaLGFVGDAGPAAfvelp | 157 | 167 | 129.8 | 0.000909637 |
| 1469 | Q9ULN7 | PNMA8B | Paraneoplastic antigen-like protein 8B | lelvaLLAAQDMAEVMkeeke | 454 | 464 | 129.8 | 0.000909637 |
| 1471 | Q9UDT6 | CLIP2 | CAP-Gly domain-containing linker protein 2 | rlqhqLMSTEDALRDAldqaq | 990 | 1000 | 129.8 | 0.000910203 |
| 1472 | Q9BWV1 | BOC | Brother of CDO | pdspvLEAVWDPPFHSgppcc | 1042 | 1052 | 129.8 | 0.000910486 |
| 1473 | Q96JE9 | MAP6 | Microtubule-associated protein 6 | frawpLPRRGDHPWIPkpvqi | 165 | 175 | 129.8 | 0.000910769 |
| 1476 | O43164 | PJA2 | E3 ubiquitin-protein ligase Praja-2 | rhersLGRAGDDYEVLelddv | 56 | 66 | 129.8 | 0.000911335 |
| 1474 | P07384 | CAPN1 | Calpain-1 catalytic subunit | qfkirLDETDDPDDYGdresg | 401 | 411 | 129.8 | 0.000911335 |
| 1475 | Q8TE85 | GRHL3 | Grainyhead-like protein 3 homolog | svdsyLLPTTDMYDNGslnsl | 164 | 174 | 129.8 | 0.000911335 |
| 1477 | Q9UNA1 | ARHGAP26 | Rho GTPase-activating protein 26 | pchpnLHLLFDRPEEAvheds | 739 | 749 | 129.8 | 0.000911713 |
| 1478 | Q8IW50 | FAM219A | Protein FAM219A | kplvaLDTDSDDDFDMsryss | 111 | 121 | 129.8 | 0.000912184 |
| 1479 | P0CE71 | OCM2 | Putative oncomodulin-2 | diaaaLQECQDPDTFEpqkff | 16 | 26 | 129.8 | 0.000912656 |
| 1480 | Q9ULF5 | SLC39A10 | Zinc transporter ZIP10 | lsdhkLNNTPDSDWLQlkpla | 550 | 560 | 129.8 | 0.000912656 |
| 1481 | P54819 | AK2 | Adenylate kinase 2, mitochondrial | itgepLIRRSDDNEKAlkirl | 172 | 182 | 129.8 | 0.000913034 |
| 1482 | Q5VWP3 | MLIP | Muscular LMNA-interacting protein | qqteeLCATIDKVLQDslsmh | 264 | 274 | 129.8 | 0.000913034 |
| 1483 | O15067 | PFAS | Phosphoribosylformylglycinamidine synthase | rglapLHWADDDGNPTeqypl | 1257 | 1267 | 129.8 | 0.000913789 |
| 1485 | Q5VWG9 | TAF3 | Transcription initiation factor TFIID subunit 3 | krprlLSTKGDTLDVVllear | 198 | 208 | 129.8 | 0.000914638 |
| 1484 | Q6ZMZ3 | SYNE3 | Nesprin-3 | pksgfLINPMDPIPRHrrran | 779 | 789 | 129.8 | 0.000914638 |
| 1486 | P08151 | GLI1 | Zinc finger protein GLI1 | mvgsgLNPYMDFPPTDtlgyg | 718 | 728 | 129.8 | 0.00091511 |
| 1487 | P18583 | SON | Protein SON | astlsLVNKYDVDLSLttqdt | 1622 | 1632 | 129.8 | 0.00091511 |
| 1488 | Q8IWI9 | MGA | MAX gene-associated protein | vglaeLPSSMDTEFPGdarra | 2847 | 2857 | 129.8 | 0.000916431 |
| 1490 | A2A3L6 | TTC24 | Tetratricopeptide repeat protein 24 | nylhaLQAARDSGDMKgqwqa | 340 | 350 | 129.8 | 0.000916714 |
| 1489 | O00255 | MEN1 | Menin | rypmaLGNLADLEELEptpgr | 285 | 295 | 129.8 | 0.000916714 |
| 1491 | Q15052 | ARHGEF6 | Rho guanine nucleotide exchange factor 6 | savnvLTQHSDELEQFmenqg | 345 | 355 | 129.8 | 0.000920016 |

| | | | | | | | | |
|---|---|---|---|---|---|---|---|---|
| 1495 | O75309 | CDH16 | Cadherin-16 | dtytvLVEAQDTDEPRlsasa | 635 | 645 | 129.8 | 0.000920394 |
| 1493 | Q9BQE6 | LBHD1 | LBH domain-containing protein 1 | tfnagLSWGQDQDEEDacwil | 150 | 160 | 129.8 | 0.000920394 |
| 1494 | Q9HBJ7 | USP29 | Ubiquitin carboxyl-terminal hydrolase 29 | pvadsLMDQGDISLPVmyedg | 651 | 661 | 129.8 | 0.000920394 |
| 1492 | Q9NQZ8 | ZNF71 | Endothelial zinc finger protein induced by tumor necrosis factor alpha | mkeLDPKNDISEDKlsvvg | 4 | 14 | 129.8 | 0.000920394 |
| 1496 | Q96PV7 | FAM193B | Protein FAM193B | lddvfLPKDMDGVEMDetdre | 850 | 860 | 129.8 | 0.000921243 |
| 1497 | P46013 | MKI67 | Proliferation marker protein Ki-67 | ekaraLEDLVDFKELFsapgh | 2556 | 2566 | 129.8 | 0.000921432 |
| 1498 | Q7Z494 | NPHP3 | Nephrocystin-3 | dfgdvLWDIHDEQEQMetfqq | 481 | 491 | 129.8 | 0.00092247 |
| 1499 | P50222 | MEOX2 | Homeobox protein MOX-2 | igaatLQQTGDSIANEdshds | 280 | 290 | 129.8 | 0.000922753 |
| 1500 | Q9NRD9 | DUOX1 | Dual oxidase 1 | ltpqrLQCPMDTDPPQeirrr | 986 | 996 | 129.8 | 0.000923508 |
| 1502 | O60239 | SH3BP5 | SH3 domain-binding protein 5 | malknLEMISDEIHERrssa | 250 | 260 | 129.7 | 0.00092398 |
| 1501 | Q9P2M7 | CGN | Cingulin | kelqnMKRLLDQGEDLrhgle | 421 | 431 | 129.7 | 0.00092398 |
| 1503 | O75449 | KATNA1 | Katanin p60 ATPase-containing subunit A1 | rvkaeLLVQMDGVGGTsendd | 333 | 343 | 129.7 | 0.000924074 |
| 1504 | Q5UIP0 | RIF1 | Telomere-associated protein RIF1 | aetnmLTAEMDNFVCDtvems | 2059 | 2069 | 129.7 | 0.000925489 |
| 1505 | Q8WZ42 | TTN | Titin | tiklrLSVRGDTIKVKagepv | 16328 | 16338 | 129.7 | 0.000927188 |
| 1507 | B3KU38 | IQCJ-SCHIP1 | IQCJ-SCHIP1 readthrough transcript protein | rlqnpLEQVNDGKYSFenhql | 13 | 23 | 129.7 | 0.000928886 |
| 1506 | Q1A5X6 | IQCJ | IQ domain-containing protein J | rlqnpLEQVNDGKYSFenhql | 13 | 23 | 129.7 | 0.000928886 |
| 1508 | Q92979 | EMG1 | Ribosomal RNA small subunit methyltransferase NEP1 | hksilLKNGRDPGEARpdith | 74 | 84 | 129.7 | 0.000929736 |
| 1509 | O75553 | DAB1 | Disabled homolog 1 | kavtqLELFGDMSTPPditsp | 279 | 289 | 129.7 | 0.000929924 |
| 1510 | O60641 | SNAP91 | Clathrin coat assembly protein AP180 | tappaLDIFGDLFESTpevaa | 561 | 571 | 129.7 | 0.000930396 |
| 1511 | O14529 | CUX2 | Homeobox protein cut-like 2 | kleekLQAQSDYEEIKtelsi | 356 | 366 | 129.7 | 0.000931245 |
| 1512 | Q8NFP4 | MDGA1 | MAM domain-containing glycosylphosphatidylinositol anchor protein 1 | rgsdtLSHSQDNGVDIyeply | 176 | 186 | 129.7 | 0.000932944 |
| 1516 | O95292 | VAPB | Vesicle-associated membrane protein-associated protein B/C | rcvfeLPAENDKPHDVeinki | 125 | 135 | 129.7 | 0.000933133 |
| 1513 | P16989 | YBX3 | Y-box-binding protein 3 | nprkyLRSVGDGETVEfdvve | 132 | 142 | 129.7 | 0.000933133 |
| 1514 | P67809 | YBX1 | Y-box-binding protein 1 | nprkyLRSVGDGETVEfdvve | 100 | 110 | 129.7 | 0.000933133 |
| 1517 | Q9NQ48 | LZTFL1 | Leucine zipper transcription factor-like protein 1 | nmkeiLTKKNDQIKDLrkrla | 278 | 288 | 129.7 | 0.000933133 |
| 1515 | Q9Y2T7 | YBX2 | Y-box-binding protein 2 | nprkfLRSVGDGETVEfdvve | 135 | 145 | 129.7 | 0.000933133 |
| 1518 | Q00722 | PLCB2 | 1-phosphatidylinositol 4,5-bisphosphate phosphodiesterase beta-2 | ceqdpLIAKADAQESRl | 1174 | 1184 | 129.7 | 0.000934076 |
| 1519 | O43157 | PLXNB1 | Plexin-B1 | ilpssLDYQYDTPGLWeleea | 891 | 901 | 129.7 | 0.000934642 |
| 1520 | Q8N1M1 | BEST3 | Bestrophin-3 | sdmlyLMENLDTKETDiieln | 645 | 655 | 129.7 | 0.000936152 |
| 1521 | Q86VR2 | RETREG3 | Reticulophagy regulator 3 | egsedLDGHSDPEESFardlp | 316 | 326 | 129.7 | 0.00093719 |
| 1522 | Q8N573 | OXR1 | Oxidation resistance protein 1 | nevgtLCHKTDLNNLEmaike | 409 | 419 | 129.7 | 0.000937473 |
| 1523 | Q8TAQ9 | SUN3 | SUN domain-containing protein 3 | allrdMKDGMDNNHNWnthgd | 140 | 150 | 129.7 | 0.000938889 |
| 1525 | P48740 | MASP1 | Mannan-binding lectin serine protease 1 | vkhttLHPQYDPNTFEndval | 540 | 550 | 129.7 | 0.000939926 |
| 1524 | Q01082 | SPTBN1 | Spectrin beta chain, non-erythrocytic 1 | germpLATSTDHGHNLqtvql | 1507 | 1517 | 129.7 | 0.000939926 |
| 1526 | Q9C0F1 | CEP44 | Centrosomal protein of 44 kDa | isedtLSPITDVNEAVdvsdl | 191 | 201 | 129.7 | 0.00094087 |
| 1527 | Q9H972 | C14orf93 | Uncharacterized protein C14orf93 | qrlteLCYHLDANSKHgtkan | 455 | 465 | 129.7 | 0.00094087 |
| 1528 | P43026 | GDF5 | Growth/differentiation factor 5 | aviqtLMNSMDPESTPptccv | 452 | 462 | 129.7 | 0.000941342 |
| 1529 | Q6NVH7 | SWSAP1 | ATPase SWSAP1 | hfshrLGPGRDCGLMValqtq | 129 | 139 | 129.7 | 0.000944739 |
| 1530 | Q8IVL1 | NAV2 | Neuron navigator 2 | vthstLETTFDTNVTTemsgr | 777 | 787 | 129.6 | 0.000944833 |
| 1531 | Q6V0I7 | FAT4 | Protocadherin Fat 4 | gsaslLVTLEDINDNGpmltv | 3497 | 3507 | 129.6 | 0.000945399 |
| 1532 | Q9HBM0 | VEZT | Vezatin | peeqeLEAYVDDIDIDsdfrk | 543 | 553 | 129.6 | 0.000945399 |
| 1533 | O60941 | DTNB | Dystrobrevin beta | rpptdLSFNFDANKQQrqlia | 426 | 436 | 129.6 | 0.000945494 |
| 1534 | Q96S90 | LYSMD1 | LysM and putative peptidoglycan-binding domain-containing protein 1 | teprdLFNGLDSEEEKdgeek | 93 | 103 | 129.6 | 0.000945494 |
| 1535 | Q05682 | CALD1 | Caldesmon | qeeesLGQVTDQVEVNaqnsv | 59 | 69 | 129.6 | 0.000945966 |
| 1536 | Q8IWD4 | CCDC117 | Coiled-coil domain-containing protein 117 | varrkLQEIEDRIIDEdeeve | 149 | 159 | 129.6 | 0.000946343 |
| 1537 | A8K0R7 | ZNF839 | Zinc finger protein 839 | ltslgLSMPADPCEGGarscl | 256 | 266 | 129.6 | 0.000946532 |
| 1538 | Q16621 | NFE2 | Transcription factor NF-E2 45 kDa subunit | peeyaLQQAADGTIFLvprgt | 352 | 362 | 129.6 | 0.000948796 |
| 1539 | P28324 | ELK4 | ETS domain-containing protein Elk-4 | tpsppLSSHPDIDTDIdsvas | 272 | 282 | 129.6 | 0.000948985 |
| 1540 | Q15884 | FAM189A2 | Protein FAM189A2 | pgllhLQSCGDLHTFTpagrp | 409 | 419 | 129.6 | 0.000949079 |
| 1541 | Q9UPN7 | PPP6R1 | Serine/threonine-protein phosphatase 6 regulatory subunit 1 | apcqaLVSIGDLQATFhgirs | 798 | 808 | 129.6 | 0.000949457 |
| 1542 | Q2LD37 | KIAA1109 | Transmembrane protein KIAA1109 | neelnLLRNVDANNTEnsttv | 4772 | 4782 | 129.6 | 0.000950306 |
| 1543 | Q16514 | TAF12 | Transcription initiation factor TFIID subunit 12 | kklqdLVREVDPNEQLdedve | 66 | 76 | 129.6 | 0.000950589 |
| 1544 | Q86TG7 | PEG10 | Retrotransposon-derived protein PEG10 | ecpedLPEKFDGNPDMlapfm | 82 | 92 | 129.6 | 0.000950589 |
| 1545 | Q8IXQ3 | C9orf40 | Uncharacterized protein C9orf40 | lppidLADIEDLSEDTlteat | 165 | 175 | 129.6 | 0.000950589 |
| 1546 | Q9NUQ3 | TXLNG | Gamma-taxilin | agicgLGVKADMLCNSqsndi | 48 | 58 | 129.6 | 0.000950589 |
| 1547 | Q6WRI0 | IGSF10 | Immunoglobulin superfamily member 10 | yreltLQRRGDSTHRRfrenr | 726 | 736 | 129.6 | 0.000950872 |
| 1548 | Q8TE56 | ADAMTS17 | A disintegrin and metalloproteinase with thrombospondin motifs 17 | nktttLVNDSDCPQASrpepq | 837 | 847 | 129.6 | 0.000950872 |
| 1549 | Q14CX7 | NAA25 | N-alpha-acetyltransferase 25, NatB auxiliary subunit | pndrrLRPIYDYLDNGnnkma | 15 | 25 | 129.6 | 0.000951438 |
| 1550 | Q13164 | MAPK7 | Mitogen-activated protein kinase 7 | lrhpfLAKYHDPDDEPdcapp | 347 | 357 | 129.6 | 0.00095191 |
| 1551 | O00321 | ETV2 | ETS translocation variant 2 | pdsqaLPWSGDWTDMActawd | 85 | 95 | 129.6 | 0.000952193 |
| 1552 | Q9P0U4 | CXXC1 | CXXC-type zinc finger protein 1 | fddhgLPWMSDTEESPfldpa | 304 | 314 | 129.6 | 0.000952571 |
| 1553 | Q9H6I2 | SOX17 | Transcription factor SOX-17 | pwaesLSPIGDMKVKGeapan | 36 | 46 | 129.6 | 0.000953137 |
| 1554 | O60315 | ZEB2 | Zinc finger E-box-binding homeobox 2 | hnneiLQASVDGPEEMkedyd | 103 | 113 | 129.6 | 0.000953797 |
| 1555 | Q9NXG2 | THUMPD1 | THUMP domain-containing protein 1 | eayslLNEYGDDMYGPekftd | 65 | 75 | 129.6 | 0.000953797 |
| 1556 | O75970 | MPDZ | Multiple PDZ domain protein | kqeaeLMSREDVTKDAdlspv | 470 | 480 | 129.6 | 0.000954175 |
| 1557 | Q92949 | FOXJ1 | Forkhead box protein J1 | gelepLKGNFDWEAIFdagtl | 305 | 315 | 129.6 | 0.000958138 |
| 1558 | Q9H9Y2 | RPF1 | Ribosome production factor 1 | tspkiLITTSDRPHGRtvrlc | 145 | 155 | 129.6 | 0.000958987 |
| 1560 | Q8WVB6 | CHTF18 | Chromosome transmission fidelity protein 18 homolog | dyeqeLCGVEDDFHNQfaael | 8 | 18 | 129.6 | 0.000959365 |
| 1559 | Q96RW7 | HMCN1 | Hemicentin-1 | appptLTWYKDGHPLTssdkv | 2809 | 2819 | 129.6 | 0.000959365 |
| 1561 | P51003 | PAPOLA | Poly(A) polymerase alpha | lndssLDLSMDSDNSMsvpsp | 523 | 533 | 129.6 | 0.000959742 |
| 1562 | Q68CR7 | LRRC66 | Leucine-rich repeat-containing protein 66 | pfekpLISAPDSGMYKthlen | 777 | 787 | 129.6 | 0.00096012 |
| 1563 | Q14524 | SCN5A | Sodium channel protein type 5 subunit alpha | frrrdLGSEADFADDEnstag | 537 | 547 | 129.6 | 0.000961063 |
| 1564 | Q9Y4K1 | CRYBG1 | Beta/gamma crystallin domain-containing protein 1 | skslvLENVTDTAQDIpttvd | 420 | 430 | 129.6 | 0.000961063 |
| 1566 | Q86XP3 | DDX42 | ATP-dependent RNA helicase DDX42 | hnlglLHGDMDQSERNkvisd | 528 | 538 | 129.6 | 0.000961535 |
| 1565 | Q9H6P5 | TASP1 | Threonine aspartase 1 | krkleLAERVDTDFMQlkkrr | 205 | 215 | 129.6 | 0.000961535 |
| 1567 | P35527 | KRT9 | Keratin, type I cytoskeletal 9 | emeqnLRQGVDADINGlrqvl | 249 | 259 | 129.6 | 0.000962007 |
| 1568 | Q8TF72 | SHROOM3 | Protein Shroom3 | qslpsLCSTSDPDTPLgapst | 1466 | 1476 | 129.6 | 0.000962101 |
| 1569 | Q9BXT5 | TEX15 | Testis-expressed protein 15 | slsfdLSRNTDVNHTSenqns | 860 | 870 | 129.6 | 0.000962101 |
| 1570 | O75376 | NCOR1 | Nuclear receptor corepressor 1 | prthrLITLADHICQIitqdf | 2048 | 2058 | 129.6 | 0.000963045 |
| 1572 | P48595 | SERPINB10 | Serpin B10 | qmaqvLQFNRDQGVKCdpese | 58 | 68 | 129.6 | 0.000963422 |
| 1571 | Q99708 | RBBP8 | DNA endonuclease RBBP8 | tatkkLHTHGDKQDKVkqkaf | 761 | 771 | 129.6 | 0.000963422 |
| 1573 | Q9H3D4 | TP63 | Tumor protein 63 | ytnlgLLNSMDQQIQNgssst | 116 | 126 | 129.6 | 0.000963517 |
| 1574 | Q8WUQ7 | CACTIN | Cactin | lsrqqLQVTGDASESAediff | 592 | 602 | 129.6 | 0.0009638 |
| 1575 | P28715 | ERCC5 | DNA repair protein complementing XP-G cells | daedsLHEWQDINLEEletle | 725 | 735 | 129.6 | 0.000966064 |
| 1576 | P49756 | RBM25 | RNA-binding protein 25 | rlkefLEDYDDDRDDPkyyrg | 500 | 510 | 129.6 | 0.000966064 |
| 1577 | O60941 | DTNB | Dystrobrevin beta | vpprpLTNMNDTMVSHmssgv | 339 | 349 | 129.6 | 0.00096748 |
| 1579 | Q14151 | SAFB2 | Scaffold attachment factor B2 | rrpydLDRRDDAYWPEgkrva | 734 | 744 | 129.6 | 0.000967763 |
| 1578 | Q15424 | SAFB | Scaffold attachment factor B1 | rrpydLDRRDDAYWPEakraa | 716 | 726 | 129.6 | 0.000967763 |

| | | | | | | | | |
|---|---|---|---|---|---|---|---|---|
| 1580 | Q8TAQ2 | SMARCC2 | SWI/SNF complex subunit SMARCC2 | rhfeeLETIMDREREAleyqr | 909 | 919 | 129.5 | 0.000968423 |
| 1581 | Q96D09 | GPRASP2 | G-protein coupled receptor-associated sorting protein 2 | ntalkLRAQKDVDSDRvkqep | 364 | 374 | 129.5 | 0.000971065 |
| 1582 | P08514 | ITGA2B | Integrin alpha-IIb | gadnvLELQMDAANEGegaye | 662 | 672 | 129.5 | 0.000971632 |
| 1583 | Q32MZ4 | LRRFIP1 | Leucine-rich repeat flightless-interacting protein 1 | reikeLNELKDQIQDVegkym | 139 | 149 | 129.5 | 0.000971632 |
| 1584 | O75147 | OBSL1 | Obscurin-like protein 1 | kdgmaLDEVWDSSHFAlqpgr | 168 | 178 | 129.5 | 0.000972198 |
| 1585 | O95477 | ABCA1 | Phospholipid-transporting ATPase ABCA1 | pvnakLSPLNDEDEDVrrerq | 1885 | 1895 | 129.5 | 0.000972387 |
| 1587 | Q9NY74 | ETAA1 | Ewing's tumor-associated antigen 1 | ddierLTQQQDIRKDSktses | 620 | 630 | 129.5 | 0.000972953 |
| 1586 | Q9NY74 | ETAA1 | Ewing's tumor-associated antigen 1 | ehgakLTQQQDIRKDSktses | 649 | 659 | 129.5 | 0.000972953 |
| 1588 | O14511 | NRG2 | Pro-neuregulin-2, membrane-bound isoform | sdsppLCPAADSRTYYsldsh | 812 | 822 | 129.5 | 0.000973425 |
| 1590 | P15924 | DSP | Desmoplakin | keierLKQLIDKETNDrkcle | 1486 | 1496 | 129.5 | 0.000973425 |
| 1589 | Q5C9Z4 | NOM1 | Nucleolar MIF4G domain-containing protein 1 | sssvpLSFARDGLDYIlgale | 209 | 219 | 129.5 | 0.000973425 |
| 1592 | Q9Y266 | NUDC | Nuclear migration protein nudC | eeaerLQLEIDQKKDAenhea | 115 | 125 | 129.5 | 0.000973708 |
| 1591 | Q9Y3X0 | CCDC9 | Coiled-coil domain-containing protein 9 | grragLGSAGDMTLSMtgrer | 246 | 256 | 129.5 | 0.000973708 |
| 1593 | Q14966 | ZNF638 | Zinc finger protein 638 | ftldeLIDQDDCISHSepkdv | 1648 | 1658 | 129.5 | 0.000974179 |
| 1594 | Q6NSI3 | FAM53A | Protein FAM53A | kslcsLNYEDDDEDDTpvktv | 305 | 315 | 129.5 | 0.000974557 |
| 1595 | Q6UB99 | ANKRD11 | Ankyrin repeat domain-containing protein 11 | pqqeeLPLSSDMVEKQtgkkd | 15 | 25 | 129.5 | 0.000976067 |
| 1597 | Q12955 | ANK3 | Ankyrin-3 | nvpetMNEVLDMSDDEvrkan | 840 | 850 | 129.5 | 0.000976538 |
| 1596 | Q9NZN5 | ARHGEF12 | Rho guanine nucleotide exchange factor 12 | dngfgLTVSGDNPVFVqsvke | 84 | 94 | 129.5 | 0.000976538 |
| 1598 | O94916 | NFAT5 | Nuclear factor of activated T-cells 5 | hmmsaLSTNEDMQMQCelfss | 942 | 952 | 129.5 | 0.000976821 |
| 1599 | Q9UHA3 | RSL24D1 | Probable ribosome biogenesis protein RLP24 | kmvqqLQEDVDMEDAP | 153 | 163 | 129.5 | 0.000976916 |
| 1600 | Q9Y421 | FAM32A | Protein FAM32A | rhldtLTEHYDIPKVSwtk | 99 | 109 | 129.5 | 0.000977388 |
| 1601 | Q6ZSJ9 | SHISA6 | Protein shisa-6 | skyssLKRLTDKEADEyymrr | 314 | 324 | 129.5 | 0.000977671 |
| 1603 | P68104 | EEF1A1 | Elongation factor 1-alpha 1 | asgttLLEALDCILPPtrptd | 228 | 238 | 129.5 | 0.000977954 |
| 1602 | Q5VTE0 | EEF1A1P5 | Putative elongation factor 1-alpha-like 3 | asgttLLEALDCILPPtrptd | 228 | 238 | 129.5 | 0.000977954 |
| 1604 | Q9UHX1 | PUF60 | Poly(U)-binding-splicing factor PUF60 | stvmvLRNMVDPKDIDddleg | 466 | 476 | 129.5 | 0.000979086 |
| 1605 | Q6PL18 | ATAD2 | ATPase family AAA domain-containing protein 2 | taeavLQKMDDMKKMRrqrmr | 185 | 195 | 129.5 | 0.000979181 |
| 1608 | P48788 | TNNI2 | Troponin I, fast skeletal muscle | elckqLHAKIDAAEEEkydme | 68 | 78 | 129.5 | 0.000979935 |
| 1606 | Q05DH4 | FAM160A1 | Protein FAM160A1 | vmvyrLCAEKDSEDMKdsqee | 647 | 657 | 129.5 | 0.000979935 |
| 1607 | Q5JR59 | MTUS2 | Microtubule-associated tumor suppressor candidate 2 | qqnedLKARIDQNTVVtrqls | 1287 | 1297 | 129.5 | 0.000979935 |
| 1609 | Q8N163 | CCAR2 | Cell cycle and apoptosis regulator protein 2 | aldpeLLLLRDDGEEEfagak | 652 | 662 | 129.5 | 0.000980596 |
| 1612 | A0A075B6I7 | IGLV5-48 | Probable non-functional immunoglobulin lambda variable 5-48 | pprylLNYYSDSDKHQgsgvp | 69 | 79 | 129.5 | 0.00098069 |
| 1611 | Q9H8G2 | CAAP1 | Caspase activity and apoptosis inhibitor 1 | raikaLMKAGDIKKPA | 351 | 361 | 129.5 | 0.00098069 |
| 1610 | Q9NQB0 | TCF7L2 | Transcription factor 7-like 2 | gandeLISFKDEGEQEekse | 18 | 28 | 129.5 | 0.00098069 |
| 1613 | O00321 | ETV2 | ETS translocation variant 2 | fcfpdLALQGDTPTATaetcw | 35 | 45 | 129.5 | 0.000981162 |
| 1614 | Q9UKV5 | AMFR | E3 ubiquitin-protein ligase AMFR | dlsprLEETLDFGEVEvepse | 545 | 555 | 129.5 | 0.00098154 |
| 1615 | O60716 | CTNND1 | Catenin delta-1 | dprrrLRSYEDMIGEEvpsdq | 318 | 328 | 129.5 | 0.000981823 |
| 1616 | Q5VVM6 | CCDC30 | Coiled-coil domain-containing protein 30 | qkrklLYQNVDELHRQvrtlq | 410 | 420 | 129.5 | 0.000981823 |
| 1617 | Q9NQX1 | PRDM5 | PR domain zinc finger protein 5 | tdtelLIGYLDSDMEAeeeeq | 120 | 130 | 129.5 | 0.000983899 |
| 1619 | Q9ULJ7 | ANKRD50 | Ankyrin repeat domain-containing protein 50 | vvqvlLEHGADPNHADqfgrt | 1065 | 1075 | 129.5 | 0.00098437 |
| 1618 | Q9Y4B5 | MTCL1 | Microtubule cross-linking factor 1 | dglssLFNIIDHSPVVqdpfq | 1765 | 1775 | 129.5 | 0.00098437 |
| 1626 | P0DMU7 | CT45A6 | Cancer/testis antigen family 45 member A6 | ippsqLDSQIDDFTGFskdgm | 69 | 79 | 129.5 | 0.00098522 |
| 1629 | P0DMU8 | CT45A5 | Cancer/testis antigen family 45 member A5 | ippsqLDSQIDDFTGFskdgm | 69 | 79 | 129.5 | 0.00098522 |
| 1625 | P0DMU9 | CT45A10 | Cancer/testis antigen family 45 member A10 | ippsqLDSQIDDFTGFskdgm | 69 | 79 | 129.5 | 0.00098522 |
| 1623 | P0DMV0 | CT45A7 | Cancer/testis antigen family 45 member A7 | ippsqLDSQIDDFTGFskdgm | 69 | 79 | 129.5 | 0.00098522 |
| 1624 | P0DMV1 | CT45A8 | Cancer/testis antigen family 45 member A8 | ippsqLDSQIDDFTGFskdrm | 69 | 79 | 129.5 | 0.00098522 |
| 1628 | P0DMV2 | CT45A9 | Cancer/testis antigen family 45 member A9 | ippsqLDSQIDDFTGFskdrm | 69 | 79 | 129.5 | 0.00098522 |
| 1621 | Q5DJT8 | CT45A2 | Cancer/testis antigen family 45 member A2 | ippsqLDSQIDDFTGFskdrm | 69 | 79 | 129.5 | 0.00098522 |
| 1622 | Q5HYN5 | CT45A1 | Cancer/testis antigen family 45 member A1 | ippsqLDSQIDDFTGFskdrm | 69 | 79 | 129.5 | 0.00098522 |
| 1627 | Q8NHU0 | CT45A3 | Cancer/testis antigen family 45 member A3 | ippsqLDSQIDDFTGFskdrm | 69 | 79 | 129.5 | 0.00098522 |
| 1620 | Q9HBT8 | ZNF286A | Zinc finger protein 286A | thmnsLSEETDHKHDVywksf | 183 | 193 | 129.5 | 0.00098522 |
| 1630 | O94827 | PLEKHG5 | Pleckstrin homology domain-containing family G member 5 | rlprgLRFDHDSWEEEydede | 348 | 358 | 129.5 | 0.000985974 |
| 1631 | Q13387 | MAPK8IP2 | C-Jun-amino-terminal kinase-interacting protein 2 | esepdLSEDADSPWLLsnlvs | 341 | 351 | 129.5 | 0.000987012 |
| 1632 | O75152 | ZC3H11A | Zinc finger CCCH domain-containing protein 11A | slkerLGMSADPDNEDatdkv | 335 | 345 | 129.5 | 0.000987296 |
| 1633 | Q14721 | KCNB1 | Potassium voltage-gated channel subfamily B member 1 | pllpvLGMYHDPLRNRgsaaa | 688 | 698 | 129.5 | 0.000987579 |
| 1634 | Q8WXI7 | MUC16 | Mucin-16 | eilatLAATTDIETIHpsink | 4522 | 4532 | 129.5 | 0.000988333 |
| 1635 | O95373 | IPO7 | Importin-7 | tiieaLRGTMDPALREaaerq | 10 | 20 | 129.5 | 0.000988428 |
| 1636 | Q8TBC3 | SHKBP1 | SH3KBP1-binding protein 1 | lamwdLTTAMDGLGQApaggl | 578 | 588 | 129.5 | 0.000988428 |
| 1637 | P29353 | SHC1 | SHC-transforming protein 1 | gkeppLGGVVDMRLREgaapg | 360 | 370 | 129.5 | 0.000988805 |
| 1638 | O95810 | CAVIN2 | Caveolae-associated protein 2 | lhtvdLSSDDDLPHDEealed | 202 | 212 | 129.5 | 0.000989183 |
| 1639 | Q9Y6Q6 | TNFRSF11A | Tumor necrosis factor receptor superfamily member 11A | ssenyLQKEVDSGHCPhwaas | 422 | 432 | 129.5 | 0.000989371 |
| 1640 | Q8TCU4 | ALMS1 | Alstrom syndrome protein 1 | lsfapLRGIPDKSEDTewssr | 254 | 264 | 129.5 | 0.000989655 |
| 1641 | Q8TDN6 | BRIX1 | Ribosome biogenesis protein BRX1 homolog | krqrkMKQRMDSGKTK | 343 | 353 | 129.5 | 0.000989749 |
| 1643 | P55196 | AFDN | Afadin | dsggtLRIYADSLKPNipykt | 250 | 260 | 129.4 | 0.000990598 |
| 1642 | Q8NF91 | SYNE1 | Nesprin-1 | qkeelLKSIEDIEERTdkerl | 1991 | 2001 | 129.4 | 0.000990598 |
| 1644 | P01282 | VIP | VIP peptides | pdqvsLKEDIDMLQNAlaend | 54 | 64 | 129.4 | 0.000991164 |
| 1645 | Q9BV36 | MLPH | Melanophilin | neqlpLQYLADVDTSDeesir | 305 | 315 | 129.4 | 0.000991164 |
| 1646 | Q93074 | MED12 | Mediator of RNA polymerase II transcription subunit 12 | kiegtLGVLYDQPRHVqyath | 722 | 732 | 129.4 | 0.000992014 |
| 1648 | Q8IX90 | SKA3 | Spindle and kinetochore-associated protein 3 | edrtsLVLNSDTCFENltdps | 330 | 340 | 129.4 | 0.000992297 |
| 1647 | Q8WXK1 | ASB15 | Ankyrin repeat and SOCS box protein 15 | MDTNDDPDEDHltsyd | 1 | 11 | 129.4 | 0.000992297 |
| 1649 | Q8WZ75 | ROBO4 | Roundabout homolog 4 | aelggLHWGQDYEFKVrpssg | 313 | 323 | 129.4 | 0.000992485 |
| 1650 | Q9BX66 | SORBS1 | Sorbin and SH3 domain-containing protein 1 | knasgLVLPTDMDLTKictgk | 53 | 63 | 129.4 | 0.00099324 |
| 1651 | O60266 | ADCY3 | Adenylate cyclase type 3 | rrrlrLQDLADRVVDAsedeh | 569 | 579 | 129.4 | 0.000994089 |
| 1652 | Q96MH2 | HEXIM2 | Protein HEXIM2 | nttqfLMNDRDPEEPNldvph | 147 | 157 | 129.4 | 0.000994184 |
| 1654 | Q00872 | MYBPC1 | Myosin-binding protein C, slow-type | kprpeLTWKKDGAEIDknqin | 871 | 881 | 129.4 | 0.000994373 |
| 1653 | Q16236 | NFE2L2 | Nuclear factor erythroid 2-related factor 2 | pseysLQQTRDGNVFLvpksk | 583 | 593 | 129.4 | 0.000994373 |
| 1655 | P25440 | BRD2 | Bromodomain-containing protein 2 | tappaLPTGYDSEEEEesrpm | 627 | 637 | 129.4 | 0.000994467 |
| 1656 | Q86UU0 | BCL9L | B-cell CLL/lymphoma 9-like protein | gmgaqLRGPMDVQDPMqlrgg | 580 | 590 | 129.4 | 0.000994844 |
| 1657 | Q8N680 | ZBTB2 | Zinc finger and BTB domain-containing protein 2 | isdseLQHISDSPIIDgqqqs | 320 | 330 | 129.4 | 0.000996165 |
| 1658 | Q5VTT5 | MYOM3 | Myomesin-3 | skaseLVVMGDHDAARrktei | 464 | 474 | 129.4 | 0.000996449 |
| 1659 | Q92630 | DYRK2 | Dual specificity tyrosine-phosphorylation-regulated kinase 2 | klrtnLAQMTDANGNIqqrtv | 580 | 590 | 129.4 | 0.000997015 |
| 1660 | Q9ULE0 | WWC3 | Protein WWC3 | dhinkLTQIEDPREQWrreqe | 7 | 17 | 129.4 | 0.00099777 |
| 1661 | P67936 | TPM4 | Tropomyosin alpha-4 chain | rtvakLEKTIDDLEEKlaqak | 213 | 223 | 129.4 | 0.000998147 |

**Supporting Dataset S2. Homologs of OspB by genus and species**

| Genus | Species | Designation | Coverage (%) | Identity (%) | E value | Length (aa) | Accession No. |
|---|---|---|---|---|---|---|---|
| *Shigella* | *S. flexneri* | OspB | 100 | 99.3 | 0 | 288 | WP_258249761.1 |
| | *S. sonnei* | OspB | 100 | 99.3 | 0 | 288 | WP_052978102.1 |
| | *S. dysenteriae* | OspB | 100 | 99.0 | 0 | 288 | WP_001046940.1 |
| | *S. boydii* | OspB | 100 | 98.6 | 0 | 288 | WP_171700986.1 |
| *Escherichia* | *E. coli* | OspB | 90 | 43.5 | 6e-63 | 281 | WP_096968747.1 |
| | *E. coli* | EspS | 95 | 33.3 | 4e-33 | 291 | WP_289397300.1 |
| | *E. fergusonii* | OspB | 87 | 43.5 | 2e-62 | 278 | WP_333001558.1 |
| | *E. albertii* | hypothetical protein | 93 | 35.5 | 6e-33 | 291 | WP_059275292.1 |
| | *Escherichia sp. MOD1-EC7003* | hypothetical protein, partial | 43 | 36.4 | 8e-11 | 263 | WP_105273972.1 |
| *Salmonella* | *S. enterica* | hypothetical protein | 98 | 31.5 | 6e-27 | 295 | WP_080150472.1 |
| | *S. bongori* | hypothetical protein | 83 | 35.9 | 1e-27 | 294 | WP_257709789.1 |
| | *Salmonella sp. ZJCDC-24* | hypothetical protein | 82 | 32.0 | 6e-26 | 240 | WP_410964612.1 |
| *Enterobacter* | *E. quasiroggenkampii* | EspS | 86 | 34.1 | 2e-26 | 291 | WP_417930567.1 |
| | *Enterobacter* sp. DC4 | hypothetical protein | 40 | 0.0 | 30.65 | 346 | WP_032664481.1 |
| *Citrobacter* | *C. braakii* | EspS | 85 | 35.3 | 8e-32 | 292 | WP_200108966.1 |
| | *C. rodentium* | hypothetical protein | 82 | 34.4 | 7e-27 | 291 | WP_012904699.1 |
| *Providencia* | *P. alcalifaciens* | hypothetical protein, partial | 85 | 33.1 | 1e-25 | 285 | WP_036983279.1 |
| | *P. alcalifaciens* | hypothetical protein | 85 | 32.7 | 8e-25 | 412 | WP_224051083.1 |
| | *P. rustigianii* | hypothetical protein | 78 | 28.9 | 9e-13 | 244 | WP_407901958.1 |
| *Grimontia* | *G. hollisae* | OspB | 83 | 35.4 | 1e-34 | 280 | WP_158174806.1 |
| *Vibrio* | *V. parahaemolyticus* | OspB | 93 | 33.7 | 5e-33 | 295 | WP_276204163.1 |
| | *V. cholerae* | hypothetical protein | 92 | 34.0 | 5e-32 | 281 | WP_000493227.1 |
| | *V. harveyi group* | OspB | 88 | 35.3 | 5e-32 | 289 | WP_080540165.1 |
| | *V. mimicus* | hypothetical protein | 86 | 31.9 | 2e-21 | 280 | WP_158139385.1 |
| | *V. ichthyoenteri* | hypothetical protein, partial | 70 | 31.9 | 3e-18 | 206 | WP_039949299.1 |
| | *V. navarrensis* | RHS repeat domain-containing prote | 92 | 23.3 | 0.003 | 980 | WP_172566106.1 |
| *Photobacterium* | *P. damselae* | hypothetical protein | 86 | 33.7 | 4e-26 | 272 | WP_086368607.1 |
| *Shewanella* | *Shewanella sp. S23-S33* | hypothetical protein | 84 | 32.1 | 1e-24 | 302 | WP_393942610.1 |
| | *S. surugensis* | hypothetical protein | 46 | 29.5 | 0.049 | 396 | WP_248938446.1 |
| *Burkholderia* | *B. ubonensis* | hypothetical protein | 91 | 27.6 | 9e-19 | 292 | WP_143136225.1 |
| *Chromobacterium* | *C. violaceum* | hypothetical protein | 84 | 27.7 | 9e-13 | 276 | WP_139794061.1 |
| | *C. amazonense* | hypothetical protein | 84 | 27.7 | 6e-12 | 276 | WP_274773425.1 |
| *Kosakonia* | *Kosakonia sp. H7A* | hypothetical protein | 80 | 24.5 | 4e-09 | 312 | WP_107147045.1 |
| | *K. radicincitans* | hypothetical protein | 80 | 24.5 | 4e-09 | 312 | WP_072439455.1 |
| | *K. oryzae* | hypothetical protein | 50 | 25.0 | 0.011 | 314 | WP_227122271.1 |
| *Erwinia* | *unclassified Erwinia* | hypothetical protein | 79 | 24.8 | 1e-08 | 312 | WP_099707632.1 |
| *Pseudomonas* | *P. baetica* | hypothetical protein | 80 | 22.8 | 4e-05 | 325 | WP_310868890.1 |
| | *Pseudomonas sp. Irchel 3E19* | hypothetical protein | 80 | 22.4 | 1e-04 | 326 | WP_141233178.1 |
| *Aeromonas* | *A. salmonicida* | hypothetical protein | 51 | 26.6 | 3e-04 | 357 | WP_271787573.1 |
| *Pseudovibrio* | *Pseudovibrio sp. Alg231-02* | OspB | 51 | 29.3 | 0.002 | 329 | WP_199914859.1 |
| | *P. ascidiaceicola* | OspB | 51 | 26.6 | 0.019 | 296 | WP_093523678.1 |
| *Iodobacter* | *Iodobacter sp. LRB* | hypothetical protein, partial | 50 | 28.8 | 0.016 | 217 | WP_337680879.1 |
| Bacteriophages | *Escherichia phage 2B8* | Ibe | 80 | 35.7 | 3e-28 | 244 | YP_009909863.1 |
| | *Stx2-converting phage 1717* | Ibe | 57 | 35.7 | 4e-20 | 201 | YP_002274285.1 |
| | *Enterobacteria phage BP-4795* | Ibe | 58 | 37.2 | 1e-20 | 201 | YP_001449317.1 |

**Extended Data Table S3. Bacterial and yeast strains used in this study**

| Strain | Description | Source |
|---|---|---|
| *E. coli* | | |
| DH10B | *str. K-12 F*$^{-}$, *mcrA*$^{-}$, Δ(*mrr*, *hsdRMS*, *mcrBC*), Φ80 *lacZΔM15*, Δ*lacX74*, *recA1*, *endA1*, *araD139Δ(ara, leu)7697*, *galU*, *galK*, λ$^{-}$, *rpsL*, *nupG* | Invitrogen (ElectroMAX), #18290015 |
| DB3.1 | *gyrA462 endA1* Δ*(sr1-recA) mcrB mrr hsdS20 supE44 ara14 galK2 lacY1 proA2 rpsL20 xyl5 leuB6 mtl1* | Invitrogen, #11782-018 |
| *S. cerevisiae* | | |
| BY4741 | MATa *his3Δ1 leu2Δ0 met15Δ0 ura3Δ0* | Laboratory collection |
| BY4741 *tco89*Δ | MATa *his3Δ1 leu2Δ0 met15Δ0 ura3Δ0 tco89::KanMX4* | [1] |
| MaV103 | MATa. *Leu2-3, 112 trp-901 his3D200 ade2-1 gal4D gal80D SPAL10::URA3 GAL1::lacZ GAL1::HIS3-@LYS2 can1*$^{R}$ *cyh2*$^{R}$.Yeast host strain for two-hybrid assays. | Laboratory collection |
| MaV103 *tco89*Δ*ate1*Δ | MATa. *Leu2-3, 112 trp-901 his3D200 ade2-1 gal4D gal80D SPAL10::URA3 GAL1::lacZ GAL1::HIS3-@LYS2 can1*$^{R}$ *cyh2*$^{R}$ *ate1::kanMX4 tco89::hphMX6* | This study |

### References


1 Wood, T. E. et al. *mBio*, e0127022 (2022).

**Extended Data Table S4. Plasmids used in this study.**

| Plasmid | Purpose | Source |
|---|---|---|
| pCMV-RFP-$FLAG_3$ | Transfection vector expressing RFP-$FLAG_3$ | This study |
| pCMV-RFP-Tco89p(2-347)-$FLAG_3$ | Transfection vector expressing RFP-Tco89p(2-347)-$FLAG_3$ | This study |
| pCMV-RFP-Tco89p(254-347)-$FLAG_3$ | Transfection vector expressing RFP-Tco89p(254-347)-$FLAG_3$ | This study |
| pCMV-RFP-Tco89p(287-347)-$FLAG_3$ | Transfection vector expressing RFP-Tco89p(287-347)-$FLAG_3$ | This study |
| pCMV-RFP-Tco89p(305-347)-$FLAG_3$ | Transfection vector expressing RFP-Tco89p(305-347)-$FLAG_3$ | This study |
| pCMV-RFP-Tco89p(287-495)-$FLAG_3$ | Transfection vector expressing RFP-Tco89p(287-495)-$FLAG_3$ | This study |
| pCMV-RFP-Tco89p(295-309)-$FLAG_3$ | Transfection vector expressing RFP-Tco89p(295-309)-$FLAG_3$ | This study |
| pCMV-RFP-Tco89p(295-309)(S296A)-$FLAG_3$ | Transfection vector expressing RFP-Tco89p(295-309)(S296A)-$FLAG_3$ | This study |
| pCMV-RFP-Tco89p(295-309) (L296A)-$FLAG_3$ | Transfection vector expressing RFP-Tco89p(295-309)(L296A)-$FLAG_3$ | This study |
| pCMV-RFP-Tco89p(295-309) (V298A)-$FLAG_3$ | Transfection vector expressing RFP-TCO89(295-309)(V298A)-$FLAG_3$ | This study |
| pCMV-RFP-Tco89p(295-309) (K299A)-$FLAG_3$ | Transfection vector expressing RFP-Tco89p(295-309)(K299A)-$FLAG_3$ | This study |
| pCMV-RFP-Tco89p(295-309) (A300A)-$FLAG_3$ | Transfection vector expressing RFP-Tco89p(295-309)(A300A)-$FLAG_3$ | This study |
| pCMV-RFP-Tco89p(295-309) (G301A)-$FLAG_3$ | Transfection vector expressing RFP-Tco89p(295-309)(G301A)-$FLAG_3$ | This study |
| pCMV-RFP-Tco89p(295-309) (D302A)-$FLAG_3$ | Transfection vector expressing RFP-Tco89p(295-309)(D302A)-$FLAG_3$ | This study |
| pCMV-RFP-Tco89p(295-309) (N303A)-$FLAG_3$ | Transfection vector expressing RFP-Tco89p(295-309)(N303A)-$FLAG_3$ | This study |
| pCMV-RFP-Tco89p(295-309) (I304A)-$FLAG_3$ | Transfection vector expressing RFP-Tco89p(295-309)(I304A)-$FLAG_3$ | This study |
| pCMV-RFP-Tco89p(295-309) (S305A)-$FLAG_3$ | Transfection vector expressing RFP-Tco89p(295-309)(S305A)-$FLAG_3$ | This study |

| | | |
|---|---|---|
| pCMV-RFP-Tco89p(295-309)(E306A)-FLAG$_3$ | Transfection vector expressing RFP-Tco89p(295-309)(E306A)-FLAG$_3$ | This study |
| pCMV-RFP-Tco89p(295-309)(A307A)-FLAG$_3$ | Transfection vector expressing RFP-Tco89p(295-309)(A307A)-FLAG$_3$ | This study |
| pCMV-RFP-Tco89p(295-309)(P308A)-FLAG$_3$ | Transfection vector expressing RFP-Tco89p(295-309)(P308A)-FLAG$_3$ | This study |
| pCMV-RFP-Tco89p(295-309)(Y309A)-FLAG$_3$ | Transfection vector expressing RFP-Tco89p(295-309)(Y309A)-FLAG$_3$ | This study |
| pCMV-Myc | Transfection vector expressing Myc tag; negative control for OspB expression); | This study |
| pCMV-Myc-OspB | Transfection vector expressing Myc-OspB | This study |
| pCMV-Myc-OspB(C184S) | Transfection vector expressing Myc-OspB(C184S) catalytic mutant | This study |
| pCMV-RFP-FLAG$_3$ | Transfection vector expressing negative control for RFP-FLAG$_3$ fusion constructs | This study |
| pCMV-RFP-GGA1(184-198)-FLAG$_3$ | Transfection vector expressing RFP-GGA1(184-198)-FLAG$_3$ | This study |
| pCMV-RFP-VPS41(735-750)-FLAG$_3$ | Transfection vector expressing RFP-VPS41(735-750)-FLAG$_3$ | This study |
| pCMV-RFP-VPS41(735-854)-FLAG$_3$ | Transfection vector expressing RFP-VPS41(735-854)-FLAG$_3$ | This study |
| pCMV-RFP-VPS41(735-750)(E735A)-FLAG$_3$ | Transfection vector expressing RFP-VPS41(735-750)(E735A)-FLAG$_3$ | This study |
| pCMV-RFP-VPS41(735-750)(I736A)-FLAG$_3$ | Transfection vector expressing RFP-VPS41(735-750)(I736A)-FLAG$_3$ | This study |
| pCMV-RFP-VPS41(735-750)(P737A)-FLAG$_3$ | Transfection vector expressing RFP-VPS41(735-750)(P737A)-FLAG$_3$ | This study |
| pCMV-RFP-VPS41(735-750)(N738A)-FLAG$_3$ | Transfection vector expressing RFP-VPS41(735-750)(N738A)-FLAG$_3$ | This study |
| pCMV-RFP-VPS41(735-750)(L739A)-FLAG$_3$ | Transfection vector expressing RFP-VPS41(735-750)(L739A)-FLAG$_3$ | This study |
| pCMV-RFP-VPS41(735-750)(R740A)-FLAG$_3$ | Transfection vector expressing RFP-VPS41(735-750)(R740A)-FLAG$_3$ | This study |
| pCMV-RFP-VPS41(735-750)(D741A)-FLAG$_3$ | Transfection vector expressing RFP-VPS41(735-750)(D741A)-FLAG$_3$ | This study |
| pCMV-RFP-VPS41(735-750)(S742A)-FLAG$_3$ | Transfection vector expressing RFP-VPS41(735-750)(S742A)-FLAG$_3$ | This study |
| pCMV-RFP-VPS41(735-750)(L743A)-FLAG$_3$ | Transfection vector expressing RFP-VPS41(735-750)(L743A)-FLAG$_3$ | This study |

| | | |
|---|---|---|
| pCMV-RFP-VPS41(735-750)(V744A)-$FLAG_3$ | Transfection vector expressing RFP-VPS41(735-750)(V744A)-$FLAG_3$ | This study |
| pCMV-RFP-TLN1(1565-1576)-$FLAG_3$ | Transfection vector expressing RFP-TLN1(1565-1576)-$FLAG_3$ | This study |
| pCMV-RFP-TFAP2(2-13)-$FLAG_3$ | Transfection vector expressing RFP-TFAP2(2-13)-$FLAG_3$ | This study |
| pCMV-RFP-IQGA1(1531-1542)-$FLAG_3$ | Transfection vector expressing RFP-IQGA1(1531-1542)-$FLAG_3$ | This study |
| pCMV-RFP-SPTN1(2290-2301)-$FLAG_3$ | Transfection vector expressing RFP-SPTN1(2290-2301)-$FLAG_3$ | This study |
| pCMV-RFP-IQGA1(1034-1045)-$FLAG_3$ | Transfection vector expressing RFP-IQGA1(1034-1045)-$FLAG_3$ | This study |
| pCMV-RFP-RPTOR(1010-1021)-$FLAG_3$ | Transfection vector expressing RFP-RPTOR(1010-1021)-$FLAG_3$ | This study |
| pCMV-RFP-TFAP2(193-204)-$FLAG_3$ | Transfection vector expressing RFP-TFAP2(193-204)-$FLAG_3$ | This study |
| pCMV-RFP-TLN1(1565-1576)-GST-$FLAG_3$ | Transfection vector expressing RFP-TLN1(1565-1576)-GST-$FLAG_3$ | This study |
| pRS316GAL-OspB-$FLAG_3$-$His_6$ | Yeast expression vector to produce OspB with a C-terminal triple FLAG tag and hexahistidine tag from the inducible pGAL promoter. | This study |
| pRS316GPD-OspB-$FLAG_3$-$His_6$ | Yeast expression vector to produce OspB with C-terminal triple FLAG tag and hexahistidine tag from the constitutive pGPD promoter, with CYC1 terminator. | This study |
| pRS314GPD-OspB | Yeast plasmid to produce OspB from a constitutive GPD promoter. | This study |
| pFHTFG-WT | Parental yeast vector containing STE5 promoter, FAS membrane anchor, codon-optimized $HA_3$, Tco89p(287-347), $FLAG_3$, and GAL4 transcriptional activator. | This study |
| pFHTFG-D302A | Yeast vector containing STE5 promoter, FAS membrane anchor, codon-optimized $HA_3$, Tco89p(287-347)(D302A), $FLAG_3$, and GAL4 transcriptional activator. | This study |
| pFHTFG-p6 | Yeast vector containing STE5 promoter, FAS membrane anchor, codon-optimized $HA_3$, Tco89p(287-347)(codon 297=NNK), $FLAG_3$, and GAL4 transcriptional activator. | This study |
| pFHTFG-p5 | Yeast vector containing STE5 promoter, FAS membrane anchor, codon-optimized $HA_3$, Tco89p(287-347)(codon 298=NNK), $FLAG_3$, and GAL4 transcriptional activator. | This study |
| pFHTFG-p4 | Yeast vector containing STE5 promoter, FAS membrane anchor, codon-optimized $HA_3$, Tco89p(287-347)(codon 299=NNK), $FLAG_3$, and GAL4 transcriptional activator. | This study |

| | | |
|---|---|---|
| pFHTFG-p3 | Yeast vector containing STE5 promoter, FAS membrane anchor, codon-optimized $HA_3$, Tco89p(287-347)(codon 300=NNK), $FLAG_3$, and GAL4 transcriptional activator. | This study |
| pFHTFG-p2 | Yeast vector containing STE5 promoter, FAS membrane anchor, codon-optimized $HA_3$, Tco89p(287-347)(codon 301=NNK), $FLAG_3$, and GAL4 transcriptional activator. | This study |
| pFHTFG-p1 | Yeast vector containing STE5 promoter, FAS membrane anchor, codon-optimized $HA_3$, Tco89p(287-347)(codon 302=NNK), $FLAG_3$, and GAL4 transcriptional activator. | This study |
| pFHTFG-p1’ | Yeast vector containing STE5 promoter, FAS membrane anchor, codon-optimized $HA_3$, Tco89p(287-347)(codon 303=NNK), $FLAG_3$, and GAL4 transcriptional activator. | This study |
| pFHTFG-p2’ | Yeast vector containing STE5 promoter, FAS membrane anchor, codon-optimized $HA_3$, Tco89p(287-347)(codon 304=NNK), $FLAG_3$, and GAL4 transcriptional activator. | This study |
| pFHTFG-p3’ | Yeast vector containing STE5 promoter, FAS membrane anchor, codon-optimized $HA_3$, Tco89p(287-347)(codon 305=NNK), $FLAG_3$, and GAL4 transcriptional activator. | This study |
| pFHTFG-p4’ | Yeast vector containing STE5 promoter, FAS membrane anchor, codon-optimized $HA_3$, Tco89p(287-347)(codon 306=NNK), $FLAG_3$, and GAL4 transcriptional activator. | This study |
| pFHTFG-p5’ | Yeast vector containing STE5 promoter, FAS membrane anchor, codon-optimized $HA_3$, Tco89p(287-347)(codon 307=NNK), $FLAG_3$, and GAL4 transcriptional activator. | This study |
